# Inhibiting a promiscuous GPCR: iterative discovery of bitter taste receptor ligands

**DOI:** 10.1101/2022.11.24.517821

**Authors:** Fabrizio Fierro, Lior Peri, Harald Hübner, Alina Tabor-Schkade, Lukas Waterloo, Stefan Löber, Tara Pfeiffer, Dorothee Weikert, Tamir Dingjan, Eitan Margulis, Peter Gmeiner, Masha Y Niv

## Abstract

The human GPCR family comprises circa 800 members, activated by hundreds of thousands of compounds. Bitter taste receptors, TAS2Rs, constitute a large and distinct subfamily, expressed orally and extra-orally and involved in physiological and pathological conditions. TAS2R14 is the most promiscuous member, with over 150 agonists and 3 antagonists known prior to this study. Due to the scarcity of inhibitors and to the importance of chemical probes for exploring TAS2R14 functions, we aimed to discover new ligands for this receptor, with emphasis on antagonists. To cope with the lack of experimental structure of the receptor, we used a mixed experimental/computational methodology which iteratively improved the performance of the predicted structure. The increasing number of active compounds, obtained here through experimental screening of FDA-approved drug library, and of chemically synthesized flufenamic acid derivatives, enabled the refinement of the binding pocket, which in turn improved the structure-based virtual screening reliability. This mixed approach led to the identification of 10 new antagonists and 200 new agonists of TAS2R14, illustrating the untapped potential of rigorous medicinal chemistry for TAS2Rs. The iterative framework suggested residues involved in the activation process, is suitable for expanding bitter and bitter-masking chemical space, and is applicable to other promiscuous GPCRs lacking experimental structures.

**Graphical abstract:** 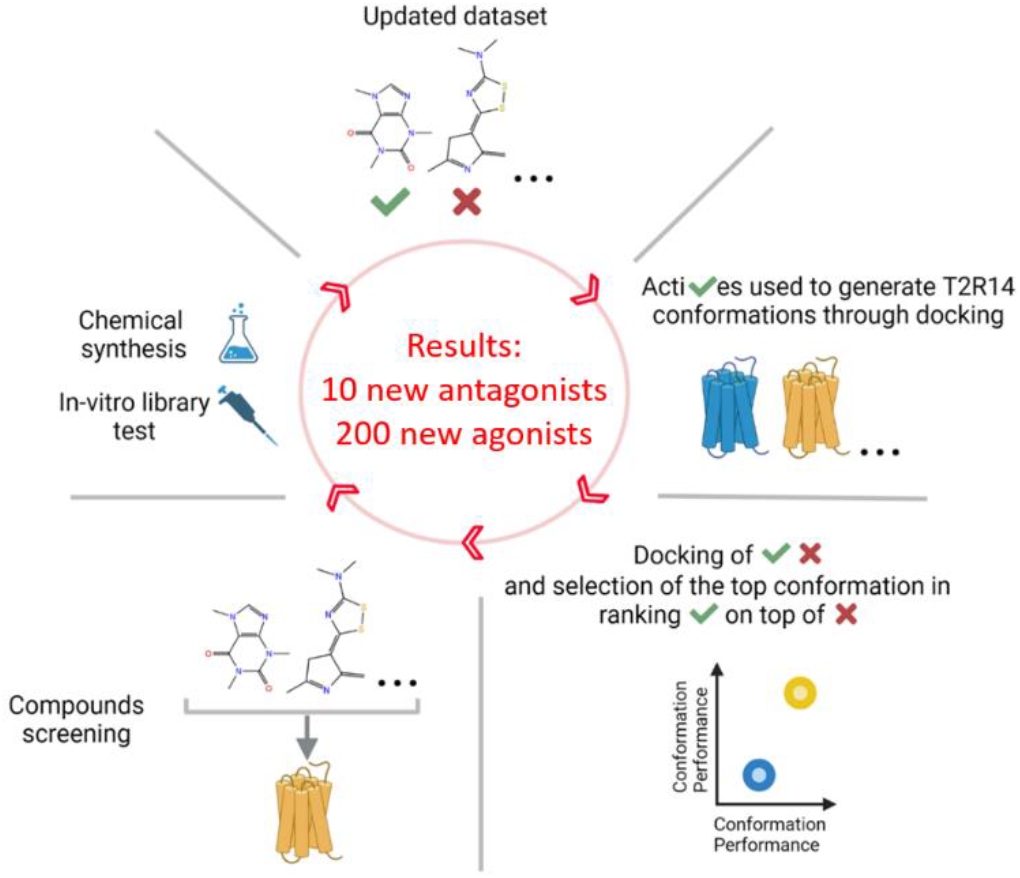

## Introduction

G-protein coupled receptors (GPCRs) are the largest family of membrane proteins, with ~800 members in humans [1]. GPCRs are involved in countless cellular processes as their main function is related to translation of extracellular signals, usually chemicals, into intracellular stimuli. A total of ~180,000 different ligands known to target human GPCRs are reported in the GPCRdb [2], defining a broad ligand spectrum for many members belonging to this family. Indeed, over 200 receptors are targeted by more than 100 ligands and ~100 by more than 1000, according to ChEMBL [3]. The long list of GPCRs targeting molecules includes over 450 drugs approved by the FDA, representing one third of the FDA-approved drugs (December 2022) [2, 4]. Interestingly, 52% of these GPCRs targeting drugs are antagonists [2].

The 25 human bitter taste receptors (TAS2Rs) belong to the GPCR family and are known to recognize over 1000 natural or synthetic bitter molecules [5], including several drugs [6, 7]. The TAS2Rs expressed within the taste buds on the tongue are responsible for the bitterness perception, which is a valued property for selected foods and beverages, but a major obstacle in many foods and in drug compliance [8–11]. The extra-oral expression of bitter taste receptors is now established [12], and their potential as new pharmacological targets (i.e. for lungs disease treatment [13, 14]) or as off-target [15] is often debated. Therefore, potent agonists and antagonists are required as biochemical probes for studying the roles of ectopic TAS2Rs, and for developing novel modulators of taste properties in foods and drugs. The most promiscuous bitter taste GPCR in humans is TAS2R14 [16, 17], which is responsible for the detection of a great variety of bitter molecules. Extra-oral expression of TAS2R14 has been confirmed in different tissues, including heart, lungs, thyroid gland and the brain [18–21] and linked with several physiological and pathological conditions [22]. For example, TAS2R14 in the human placental tissue has been suggested as contributor to the immunological regulation at the maternal-fetal interface [23]; activation of TAS2R14 expressed within the airway smooth muscle induces significant bronchodilation; increased TAS2R14 expression was associated with worse survival in adrenocortical carcinoma and esophageal adenocarcinoma [24], but prolonged overall survival in pancreatic ductal adenocarcinoma [25].

To unravel the physiological roles of TAS2Rs and TAS2R14 in particular, antagonists are of great importance. However, despite multiple experimental studies or ligand-based and structure-based guided screens [6, 26], only a few antagonists of TAS2R14 were discovered so far [27].

While this study was performed, no experimental structure was available for any of the TAS2Rs. Modeling was challenging due to low sequence identity with available structural templates (~11%) [28, 29] and subsequent lacking details on ligand binding and receptor activation mechanism.

To overcome the lack of structural data, we applied a mixed computational/experimental approach to discover TAS2R14 antagonists and agonists in an iterative methodology. In each iteration, we identified new compounds through cell-based screen of drug-like molecules and/or through synthesis of derivatives of the known agonist flufenamic acid. The new obtained dataset of known active molecules was employed to improve the performance of TAS2R14 models in their ability to discriminate actives from decoys (compounds supposed not to bind the receptor) through Induced Fit Docking (IFD) followed by Retrospective Virtual Screening (RVS). IFD allows a rearrangement of the binding pocket side chains and, consequently, new TAS2R14 conformations generation. RVS computes the IFD conformations’ performance relying on docking score-based ranking of docked actives and decoys. The top two performing 3D models, one dedicated to agonists and one to antagonists’ detection, were selected in each iteration and employed for virtual screening (VS). Specifically, two cycles of structure-based and mixed structure and ligand based screening of a multi-million compounds library [30] allowed the identification of 3 antagonists and 6 agonists, a remarkable result considering the difficulties in finding TAS2Rs antagonist and the low number (n=29) of computationally predicted compounds that were experimentally tested. In addition, the RVS performances of the AlphaFold model [31] and of a homology model based on the TAS2R46 CryoEM structure [32] (both not available at the beginning of this study) were also evaluated.

Including both experimental and computationally-driven procedures, we found a total of 200 new agonists and 10 new antagonists and highlighted new TAS2R14 structural features related to ligands binding. Comparison of TAS2R14 agonists vs antagonists dedicated models suggested new hypotheses pertaining to TAS2R14 activation.

## Results

The 3D models of TAS2R14 were shown to be useful for finding agonists through a structure-based approach [17, 33]. Starting from a model generated with I-TASSER [34] defined as IT0 throughout the manuscript, two iterations of structure refinement, each one followed by structure-based and mixed structure and ligand-based detection of TAS2R14 ligands, were applied. The models nomenclature is “IT” followed by the number of the iteration and by a “+” symbol if the model is for agonists screening, or a “-” symbol for antagonists’ screening. The ligands employed in each iteration are summarized in Table 1. Each iteration starts with a pure experimental strategy to increase the number of known active ligands that are then used within the computational part of the methodology to identify additional new actives. A third iteration was performed to get the final models.

**Table 1.** Agonists and antagonists available for each computational iteration and source details (Flu derived: flufenamic acid derivatives; BitterDB: compounds retrieved within the literature, not within this study; Comp screen: active compounds found in this work by using the computational methodology; FDA library: agonists identified through the experimental screening of the FDA-approved drug library (DiscoveryProbe™, ApexBio)). Within each iteration output, the number of output ligands includes the ones used as input for that iteration plus the ones discovered through the computational screening. For each data source, the number of new ligands compared to the previous iteration step is indicated with a “+” symbol and the total cumulative number retrieved through that source is specified within parenthesis.

| Compounds | Iteration 1 |  | Iteration 2 |  | Iteration 3 |
| --- | --- | --- | --- | --- | --- |
| Agonists | Input: **209**<br>BitterDB: 170<br>Flu derived: 39 | Output: **212**<br>Comp screen: +3 (3) | Input: **391**<br>BitterDB: +24 (194)<br>Flu derived: +14 (53)<br>FDA library: +141 (141)<br>Comp screen: +0 (3) | Output: **394**<br>Comp screen: +3 (6) | Input: **394**<br>Same ligands as iteration 2 |
| Antagonist | Input: **10**<br>BitterDB: 3<br>Flu derived: 7 | Output: **11**<br>Comp screen: +1 (1) | Input: **12**<br>BitterDB: +1 (4)<br>Flu derived: +0 (7)<br>Comp screen: +0 (1) | Output: **14**<br>Comp screen: +2 (3) | Input: **14**<br>Same ligands as iteration 2 |

### Iteration 1: chemical synthesis of new active molecules

Following a previous study [33], several flufenamic acid derivatives were synthesized and tested for their ability to activate or inhibit the receptor. Among them, 39 were agonists (Supplementary File S1 and S2), while 7 were antagonists (Table 2 and Fig. 1a). These compounds were combined with the previously known TAS2R14 actives, including 17 flufenamic acid derivatives from [35].

**Table 2.** Pharmacological data for tested compounds and the reference agonists flufenamic acid and aristolochic acid obtained by measuring Gσ_qi_ mediated accumulation of IP1^a^.

| agonistic effect |  |  |  | inhibitory effect<sup>b</sup> |  |  |  |  |  |
| --- | --- | --- | --- | --- | --- | --- | --- | --- | --- |
|  | EC<sub>50</sub> [μM]<sup>c</sup> | E<sub>max</sub> [%]<sup>d</sup> | n<sup>e</sup> | Flufenamic acid as agonist |  |  | Aristolochic acid as agonist |  |  |
|  |  |  |  | IC<sub>50</sub> [μM]<sup>f</sup> | E<sub>max</sub> [%]<sup>g</sup> | n<sup>e</sup> | IC<sub>50</sub> [μM]<sup>h</sup> | E<sub>max</sub> [%]<sup>g</sup> | n<sup>e</sup> |
| Flufen. acid | 0.43 ± 0.04 | 100 | 10 | -- | -- | -- | -- | -- | -- |
| Aristol. acid | 0.83±0.2 | 100 | 4 | -- | -- | -- | -- | -- | -- |
| LW 118 | -- | <5 | 2 | 4.0 ± 0.49 | 83 ± 4 | 8 | 2.7 ± 4.1 | 82 ± 17 | 3 |
| LW 129 | -- | <5 | 3 | 3.7 ± 0.62 | >95 | 5 | -- | -- | -- |
| LW 131 | -- | <5 | 2 | 4.1 ± 1.6 | >95 | 2 | -- | -- | -- |
| LW 145 | 3.2 ± 1.2 | -14 ± 5 | 6/11 | 3.8 ± 0.64 | 88 ± 6 | 5 | -- | -- | -- |
| LW 148 | -- | <5 | 6 | 4.4 ± 1.6 | >95 | 6 | -- | -- | -- |
| LW 209 | 0.48 ± 0.1 | -19 ± 3 | 4 | 4.1 ± 1.3 | 96 ± 4 | 3 | -- | -- | -- |
| LWYW22 | -- | <5 | 1 | 3.8 ± 1.5 | >95 | 2 | -- | -- | -- |
| LF1 | -- | -8 | 2 | 6.8 ± 3.2 | 75 ± 7 | 5 | 11 ± 6.1 | >95 | 3 |
| LF14 | -- | -15±22 | 2 | 22 ± 16 | 76 ± 17 | 3 | 14.3 ± 5.8 | 53 ± 10 | 3 |
| LF22 | -- | -29 ± 5.6 | 2 | 7.2 ± 3.7 | 63 ± 13 | 3 | 4.3 ± 2.6 | 59 ± 16 | 3 |

<sup>a</sup>Measurement of G-protein signaling was performed applying the IP-One assay<sup>®</sup> (Cisbio) in HEK293T cells transiently co-transfected with the human TAS2R14 receptor and the hybrid G-protein $G\alpha_{qi}$ . <sup>b</sup>Inhibition of the agonist effect of 1 μM of flufenamic acid/aristolochic acid. <sup>c</sup>Potency of TAS2R14 activation as mean value in μM±SEM. <sup>d</sup>Maximum efficacy in %±SEM relative to the full effect of flufenamic acid. <sup>e</sup>Number of individual experiments all performed in duplicates/triplicates. <sup>f</sup>Potency to inhibit the effect of 1 μM flufenamic acid as mean value in μM±SEM. <sup>g</sup>Maximum inhibitory efficacy was analyzed by subtracting the effect at the highest concentrations of test compound suitable for inhibition from 100% and is displayed as %±SEM. <sup>h</sup>Potency to inhibit the effect of 1 μM aristolochic acid as mean value in μM±SEM.

**Fig. 1.**
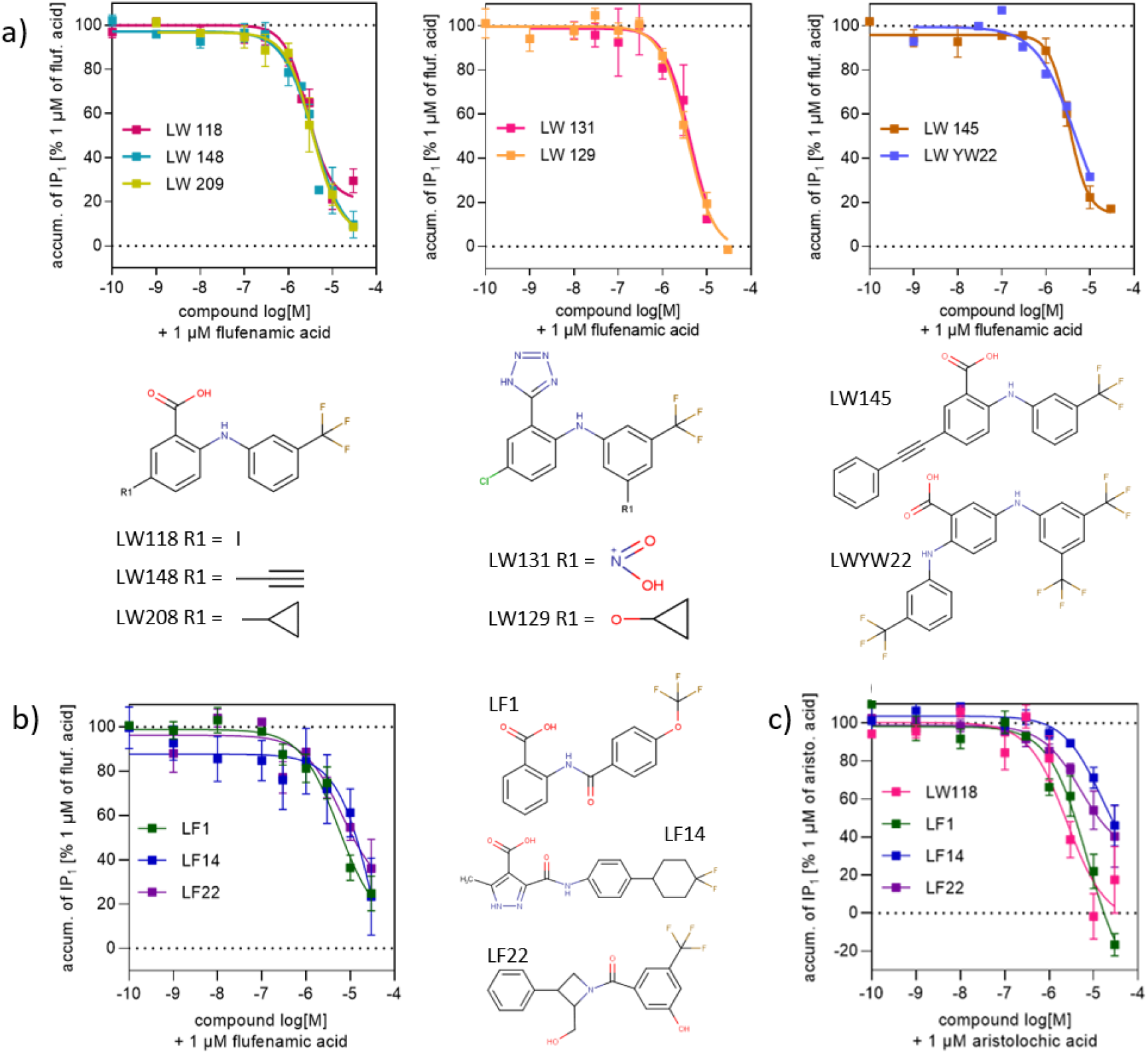
Dose dependency of the 10 TAS2R14 antagonists identified in this study and their 2D structure. Values represent the means, error bars indicate SEM of 2-11 independent experiments, each treatment performed in three replicates. **a)** Derivatives of flufenamic acid showing antagonistic effect on TAS2R14 in the presence of 1μM of flufenamic acid. **b)** Antagonists identified through the computational methodology having a different scaffold compared to the flufenamic acid derivatives. **c)** Dose dependency of the inhibitory effect of 4 TAS2R14 antagonists identified in this study in the presence of 1μM aristolochic acid.

### Iteration 1: conformations generation and selection

The dataset of TAS2R14 active compounds included 209 agonists and 10 antagonists (Table 1). These molecules were used to create circa 1900 conformations of the protein by applying IFD (Schrödinger Release 2021-1: Induced Fit Docking protocol, Schrödinger, LLC, New York, NY, 2021) to the IT0 starting model. The same ligands were also employed, together with a set of decoys, to evaluate the conformations quality through RVS. The model with the highest Enrichment Factor 10% (EF10%) and Area Under the Curve (AUC) for antagonists was chosen for further antagonists screening (model name: IT1-) and the model with the highest EF1% and AUC for agonist discrimination was chosen for further agonists screening (model name: IT1+) (see paragraph “Iteration 3: final models, top performances” for models performances).

### Iteration 1: virtual screening and ligands’ clustering strategies

The Enamine REAL compound library, a drug-like dataset of molecules complying the “rule of 5” and lacking PAINS and toxic compounds, comprised ~23 million compounds when this study started [36]. The compounds were docked to both IT1- and IT1+, ~6000 compounds with high docking scores to both models were selected. Indeed, while IT1-prioritized antagonists only over decoys, IT1+ prioritized both agonists and antagonists over decoys, a feature that can be exploited to increase the chances of new antagonists’ identification (Supplementary Fig. S1). Among the ~6000 compounds, 7 were selected for purchase and testing. Two of them showed agonistic activity (LF6 and LF9, Supplementary Table S1), while no antagonists were found.

The structure-based virtual screening results were also used as a filter for a ligand-based strategy: the ~6000 compounds were clustered, together with the active ligands, using a hierarchical clustering algorithm. Three Enamine compounds that clustered together with known actives were selected for purchase and experimental testing, finding one new antagonist (LF1) (Fig. 1b and Table 2) and one new agonist (LF3) (Supplementary Table S1).

### Iteration 2: cell-based screening of drug-like compounds and new chemical synthesis

In parallel to the first iteration of the computational methodology, a large cell-based in vitro screening of an FDA-approved drug library (DiscoveryProbe™, ApexBio) was performed, using the IP -One HTRF^®^ assay (384-well format) to asses TAS2R14 activation. The library included 1791 compounds with well-characterized bioactivity, bioavailability, and safety profiles and was primary screened at a final single concentration of 30 μM [37, 38]. With this criterion, 161 library compounds were identified as active (141 previously unknown) (Supplementary File S3). The active library compounds from the primary screen were further tested for their activity at 3 μM and thereafter at 1 μM and 0.3 μM, resulting in 38, 24 and 9 active compounds, respectively. These numbers take into account the results of a counter-screen against TAS2R14 uninfected HEK293T cells (mock cells), which suggested five false positives compounds that were excluded from the list of agonists due to unspecific accumulation of inositol monophosphate (IP) (Supplementary File S3).

The inactive compounds from the FDA screening were docked to the IT2-model and 10 top ranking compounds were selected for antagonism test, but none showed antagonistic activity.

Additionally, 14 new agonists were identified through chemical modification of the flufenamic acid (some of them presented in reference [35] and some within this study) (Supplementary File S1 and S2). The FDA library compounds and the new flufenamic acid derivatives were combined with the updated list of actives retrieved within the literature and with the compounds obtained in iteration 1, and used as input set for iteration 2 (Table 1).

### Iteration 2: conformations generation and selection

IT2+ and IT2-were generated starting from the top performing model of iteration 1. The active dataset now included 391 agonists and 12 antagonists (Table 1), and was employed for IFD models’ generation and RVS for performance evaluation. Selection of the IT2 models was performed as for iteration 1. Unlike iteration 1, the best agonist discriminating model did not get satisfying results in antagonist detection and thus a VS strategy differing from the one applied in iteration 1 was employed.

### Iteration 2: virtual screening and tanimoto similarity + clustered docking strategies

The same Enamine library employed for VS during iteration 1 was docked to the IT2- and IT2+ models. Seven compounds were chosen for *in-vitro* test based on the IT2-: one was experimentally found to be an antagonist (LF22, Table 2 and Fig. 1b), and two were found to be agonists (LF25 and LF26, Supplementary Table S1). Five compounds were selected based on a high docking score in the IT2+ and no agonists nor antagonists were identified.

An additional approach for candidates’ selection was based on the assumption that, usually, similar ligands bind in a similar way to the same receptor. Hence, we first looked for active-2D similar compounds through Tanimoto score within the Enamine REAL library. Then, among these selected molecules, the ones with a binding pose similar to the topologically similar active molecule were chosen. Particularly, by docking an active compound to all of the iteration 2 IFD conformations and by clustering the 10 binding poses that got the highest docking score, we obtained one representative pose for the active. Similarly, we got a representative binding pose for the similar candidates found within the library. A structural comparison was then performed and some candidates selected based on binding pose similarity with the docking of the active. This strategy was applied to the available antagonists, with 4 Enamine REAL compounds reaching the *in-vitro* test. One antagonist (LF14, similar to the flufenamic acid derivative agonist lw209) (Table 2 and Fig. 1b) and 1 agonist (LF11, similar to the antagonist LF1 found in iteration 1) (Supplementary Table S1) were found. The same strategy was also applied to the 14 agonists that activated TAS2R14 at 0.3μM, the lowest concentration tested in the FDA library screening. Three compounds similar to the known agonists (two to butoconazole nitrate, one to sulconazole nitrate) having a similar predicted binding orientation or interaction pattern with the receptor when compared to the docking of the agonist, were selected for testing, resulting in no hits.

### Characterization of antagonists

Among the antagonists with novel scaffolds discovered through the computational study, LF1, LF14, and LF22 were able to reduce the activity of the receptor and block flufenamic acid induced TAS2R14 activity with a half-maximal inhibitory concentration of 6.8±3.2, 22±16 and 7.2±3μM, respectively (Table 2 and Fig. 1b).

In order to investigate the antagonistic effect of the discovered compounds, another TAS2R14 reported agonist, aristolochic acid, was tested. This is a natural bitter compound known to activate 3 different TAS2Rs (TAS2R14, TAS2R43, TAS2R44) [39]. In an extensive TAS2R14 mutagenesis study, Nowak et al suggested that flufenamic acid and aristolochic acid bind differently in the receptor-binding pocket [17]. A derivative of flufenamic acid (LW118) and the compounds having different scaffolds derived from the computational study (LF1/LF14/LF22) inhibit both flufenamic and aristolochic acid in dose response manner (Fig. 1b and 1c). At the maximal tested concentration of the antagonists (30 μM), IP1 accumulation was reduced to basal level by LF14 and LF22, or to complete signal blockage by LF1 and LW118.

### Iteration 3: final models, top performances

The full set of ligands (394 agonists and 14 antagonists (Table 1)), was employed to build the last models, IT3- and IT3+. New IFD conformations (>1500) were generated starting from the model used for iteration 2 and the top performing ones in terms of RVS were selected. Evaluating all the models generated through the different iterations using the most updated dataset of ligands shows an improvement in their VS performances (Fig. 2). A substantial increase in the EF values is evident for both agonist and antagonist models when comparing the first iteration with iteration 3 results, while AUC values have a different behavior according to the model. For the antagonist, a constant increase is observed, and for the agonist a slight decrease (circa -0.03) is registered. This trend can be explained by the presence of the Extracellular Loop 2 (ECL2) acting as a lid on top of the binding site and limiting the docking of bulky compounds in our models (Supplementary Fig. S2). The largest TAS2R14 antagonist known so far has a molecular weight (MW) of 390 g/mol, while the largest agonist is almost 1000 g/mol. High MW compounds are probably most affected by the lid feature, thus influencing the agonist model performances.

Only 6 agonists were identified by IT0 within the top 168 (1%) RVS screened compounds, while 52 were identified using the IT3+ model. Five antagonists out of 14 identified by IT0 within the top 59 (10%) screened ligands, increased to 12 in IT3-model (Fig. 2a and 2b, bottom panels).

**Fig. 2.**
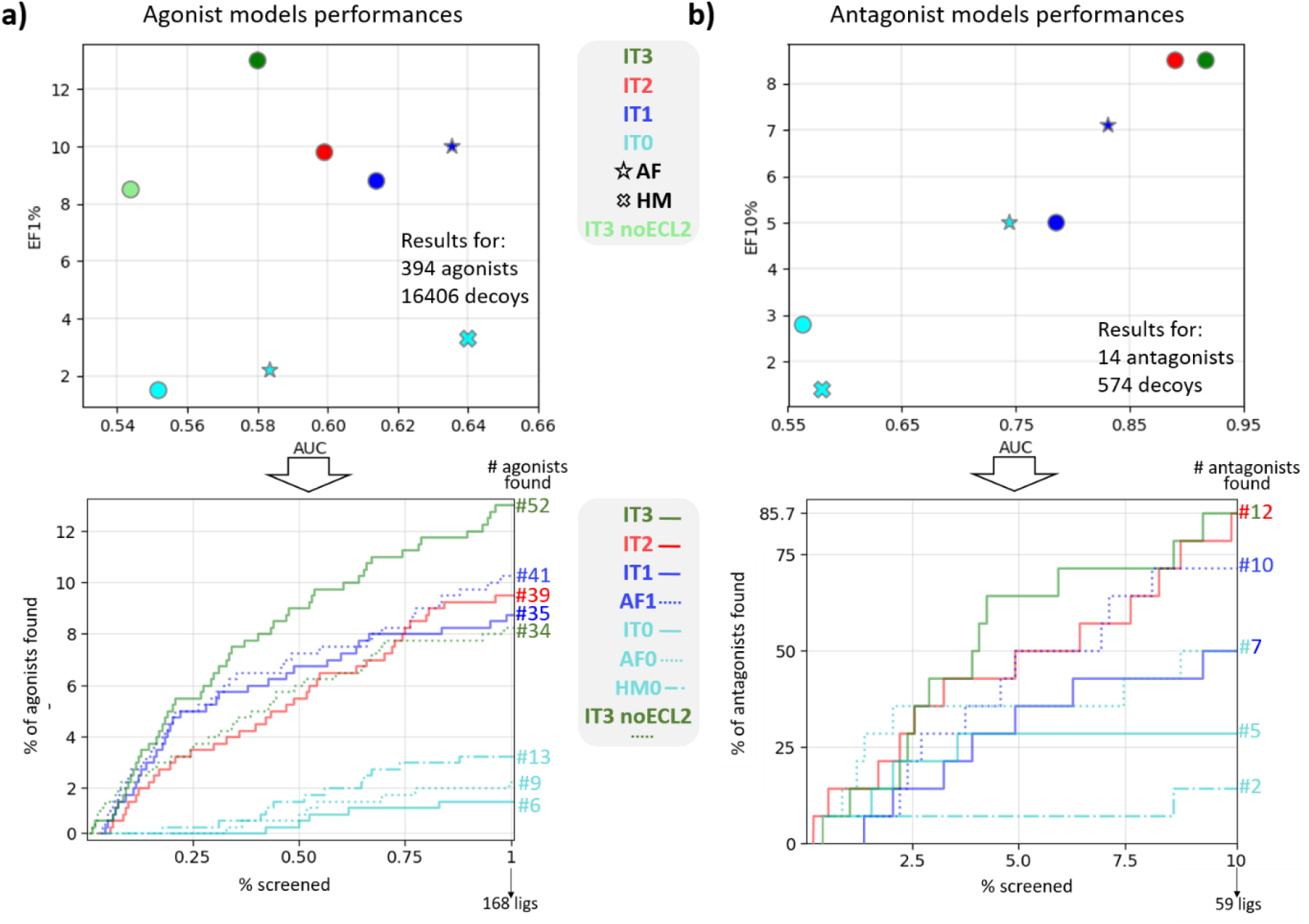
Models’ evaluation across iterations. **a)** Models’ performances in terms of AUC and EF1% calculated by using the final dataset of agonists and decoys. Circles represents the models derived from IT0, different iterations are represented with different colors. All the models within this panel are intended to be “+”. IT#=model at iteration #; AF=AlphaFold model (at iteration 0: cyan star symbol; at iteration 1: blue star symbol); HM0=homology model based on the TAS2R46 template (cyan “X” symbol: iteration 0). In the bottom panel, the AUC showing the performances obtained for the top 1% of the results, including the number of agonists identified in the top 1% of the docked results ranked according to docking score. **b)** Same as a) for the antagonists (models “-”) with EF10% on y-axis. AF0=original AlphaFold model; AF1=AlphaFold model after one iteration.

A comparison between the first and the last models generated in this work shows a massive rearrangement of the side chains within the binding site (Fig. 3a). The flufenamic acid docked to the IT3+ clashes with the F186^5.46^ of the superposed IT0. Indeed, the binding site is slightly shifted towards the intracellular side in the IT3+. As a consequence, also other residues involved in binding, such as W89^3.32^ and N93^3.36^, have different orientations. N93^3.36^ and Y240^6.48^ evolve through the generation of a H-bond in the IT3+, an interaction missing within IT0.

**Fig. 3.**
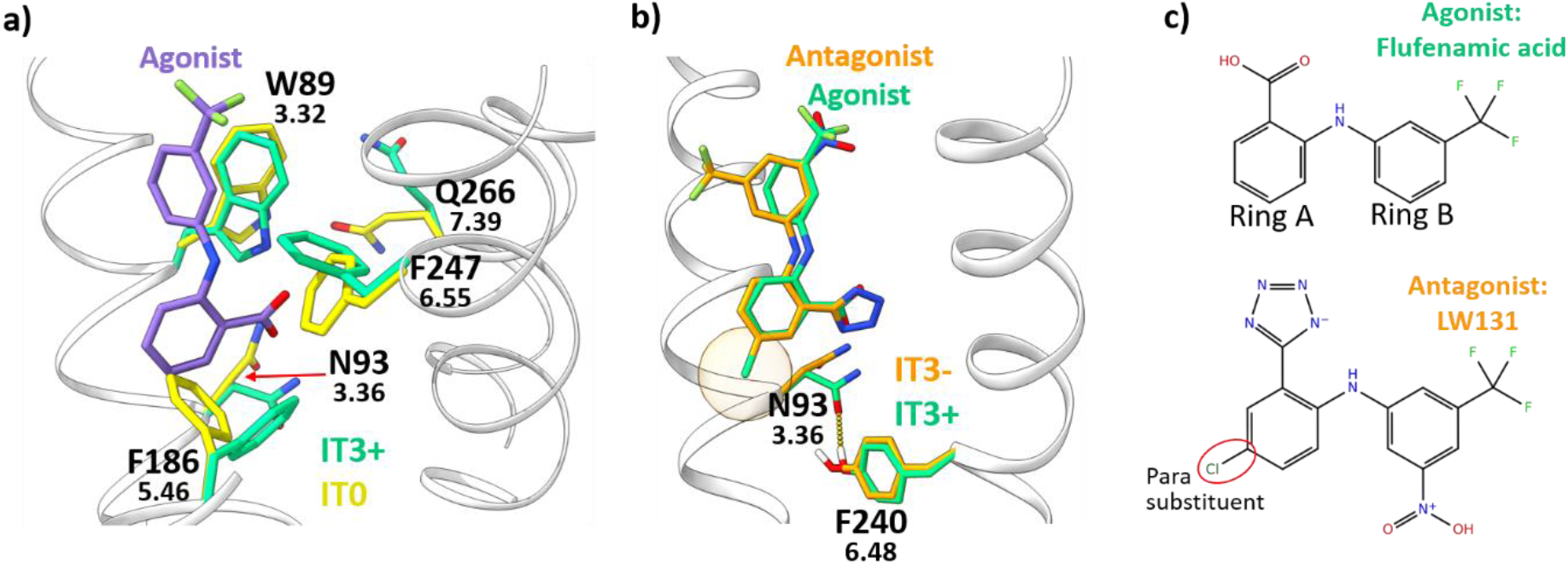
**a)** Superposition between IT0 (yellow licorice) and IT3+ (green licorice). In purple licorice the flufenamic acid (agonist) docked to the IT3+. **b)** Superposition between IT3+ in complex with flufenamic acid (both in green licorice) and IT3-in complex with the flufenamic acid derivative antagonist LW131 (both in orange licorice). The two ligands overlap within the binding site. The H-bond interaction observed only in the IT3+ between 3.36 and 6.48 is shown as a yellow dotted line and it is potentially affected by the presence of the para substituent on ring B (orange transparent circle) of the antagonist. **c)** 2D structure of the ligands from panel B. The typical arrangement for flufenamic acid derivative antagonists – substituent in position para on ring A – is shown.

#### Agonist/antagonist model comparison

There are only subtle differences between agonist and antagonist models IT3+ and IT3-, in line with the high similarity between agonists and antagonists themselves. The main structural divergence involves two positions associated with activation in several class A GPCRs, namely N93^3.36^ and Y240^6.48^ (Fig. 3b). The interaction between these two positions observed within the IT3+ is lacking in the IT3-due to a different arrangement of the N93^3.36^ side chain. In several GPCRs like MC4R, S1PR1, AGTR1 or CNR1, a similar orientation rearrangement of position 3.36 and/or 6.48 is involved in receptor activation, driving the typical TM6 movement characterizing the GPCR active state [40, 41]. The interaction between the two residues is often regulated by the nature of the binding ligand, adjusting the contact between the two residues [42–48]. Our final agonist model IT3+, when compared to IT3-, shows an additional H-bond between N93^3.36^ and Y240^6.48^ due to a displacement of N93, while Y240 remains almost unaltered. Experimental mutagenesis data on TAS2R14 show an increase in flufenamic acid EC_50_ by at least 2 orders of magnitude for both Y240A and N93A mutants [17], supporting the hypothesis that the interaction 3.36-6.48-ligand is essential for receptor activation regulation. Details regarding the role of the ligand in the 3.36-6.48 switch are reported in paragraph “Agonists/antagonist similarity and selectivity”.

#### AlphaFold model

AlphaFold, the artificial intelligence program, was a milestone in protein structure prediction [49]. While the project was in its final stages, we downloaded the AlphaFold model of TAS2R14 (Q9NYV8 (TAS2R14_HUMAN)) from the EMBL-EBI database [31], named here AF0. One of the main differences between AF0 and IT3+ model is the absence of a long extracellular loop 2 (ECL2) in AF0 (Supplementary Fig. S2). Our models have an ECL2 of circa 23 residues, potentially acting as a lid on top of the binding site (as already discussed in “Iteration 3: final models, top performances”), while AlphaFold model shows a short ECL2 made of 3 residues. AF0 and AF1+, the latter derived from one iteration of IFD and RVS by using the final set of active ligands - the same dataset used for IT3 models – exhibits better AUC values than the in-house models for the same iteration (Fig. 2a and 2b). A possible explanation is that the lid-ECL2 of our models impede the fitting of several ligands within the reduced-size binding pocket, affecting the performances. To reduce the spatial constraints, the ECL2 from the IT3+ was removed. Though this did not lead to IT3+ AUC improvement (Fig. 2a) the number of docked agonists increased by ~22% compared to the standard IT3+, suggesting that another iteration –starting from ECL2-lacking conformations could lead to improved values.

Another major difference is that AF0 lacks an interaction between 3.36 and 6.48 due to the placement of 6.48 one helix turn below compared to our I-TASSER derived models (Supplementary Fig. S2), a distance that does not allow any direct contact.

The EF results obtained with the AF models do not reach the high standard obtained with the IT3 models, suggesting IT3 as the best candidates for structure-based virtual screening.

#### Homology model based on the TAS2R46 structure

The cryoEM structure of TAS2R46 has been recently released [32]. This increases the sequence identity of TAS2R14 with available structural models from circa 11% at the beginning of this study to the current ~45% with the new TAS2R46 structure. We used the PDBid 7XP6 as a template for homology model of TAS2R14 using SwissModel [50]. The single homology model (HM0) was evaluated in its ability to discriminate agonists and antagonists from decoys, obtaining EF value for agonists discrimination slightly better than AF0 and IT0 (Fig. 2) and the top AUC registered within the models of this study. On the other hand, poor results are obtained for the antagonist model.

Similar to the AlphaFold model, this homology model space residues 3.36 and 6.48, not showing any type of interaction between these two positions.

### Agonists/antagonist similarity and selectivity

The similarity between the agonist flufenamic acid and its derivatives that act as antagonists allows a direct comparison. Small modifications to active ligands altering the molecule’s effect on the receptor have been already observed for TAS2R14 with the agonist flavanone and the 3 flavanone derivatives blockers [27], as well as for other GPCRs [51–54]. The similarity between TAS2R14 agonists and antagonists also explains why some compounds turned out to be experimental agonists, though found computationally while looking for antagonists or vice versa. Interestingly, all of the 7 antagonists found through testing of derivatives share the presence of substituents on position para of ring A (Fig. 1 and Fig. 3C). We hypothesize that the presence of a substituent in this position is responsible for the change in orientation of side chain 3.36 discussed within paragraph “Agonist/antagonist model comparison” (see Fig. 2B), affecting the activation status of the receptor.

The selectivity of the new antagonists was tested with SwissTargetPrediction, a webserver for prediction of probable protein targets of small molecules [55]. For each of the potential targets, the probability to bind it ranges from 0 to 1. The only two molecules with at least one predicted protein target - no TAS2R - with a probability higher than 0.25 are LW118 (max value=0.42) and LW209 (max value=0.22), while the average probability for all of the 292 potential targets found by the algorithm for the 14 antagonists is limited to 0.11. In addition, all the ligands found within this study underwent BitterMatch, a machine learning algorithm for prediction of TAS2R subtypes binding bitter molecules with 80% precision in prospective testing [56]. The majority (131 out of 210, ~62%) of the compounds are predicted to be selective for TAS2R14. The algorithm identifies TAS2R14 as the only human TAS2R target for all of the 10 inhibitors, potentially paving the way for specific inactivation of the receptor. All of the 53 flufenamic acid derivatives acting as agonists are predicted to be selective for TAS2R14, as well as the 6 LF agonists discovered through the computational methodology (LF3, LF6, LF9, LF11, LF25, LF26). Among the 141 new agonists discovered through the FDA library screening, 62 are predicted to be selective for TAS2R14, other 62 to activate at least one additional receptor (TAS2R10, TAS2R39, or TAS2R46), 3 are selective for TAS2R46 and 14 are predicted as not binding any TAS2R.

### TAS2R14 ligands chemical space

The final dataset of TAS2R14 active molecules includes, as of December 2022, 394 agonists and 14 antagonists. A Principle Components Analysis (PCA) of these dataset shows that the flufenamic acid derivatives (LW compounds), as well as the molecules discovered via the computational study (LF compounds) represent only a restricted portion of the chemical space (Fig. 4a). The library including the LF molecules was subject to Enamine REAL filters related to MW (≤ 500), SlogP (≤ 5), number of H-Bonds Acceptors (≤ 10) and H-Bonds Donors (≤ 5), number of rotatable bonds (≤ 10) and Topological Polar Surface Area (≤ 140), limiting the topology spectrum of ligands that can be discovered. Therefore, new possibilities for different ligands can still be explored by using different ligands libraries. The main descriptors influencing the Principal Components (PC) are related to aliphatic (PC1) and aromatic (PC2) features (Fig. 4b). Increasing the presence of these structures and, therefore, the ligand MW, would lead to exploration of new areas of the chemical space.

**Fig. 4.**
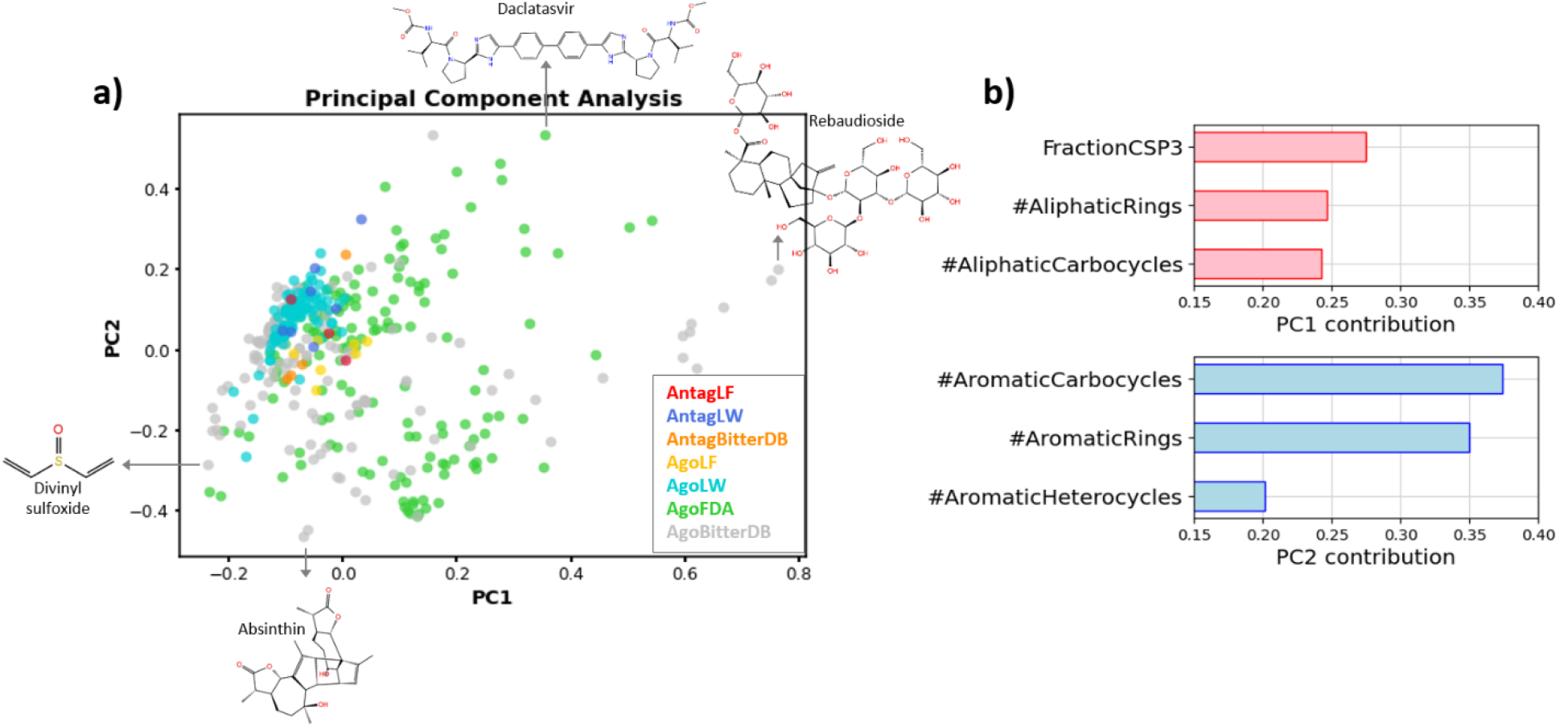
**a)** PCA showing the chemical space occupied by the TAS2R14 ligands. The molecules are colored according to source and type of activity. For each PC, the molecule with highest and lowest value are shown. **b)** For each PC, the 3 main descriptors affecting the PC are reported.

## Discussion

In this study, a mixed experimental and computational approach was applied to find 10 antagonists and 200 agonists and of a broadly tuned bitter taste GPCR, the human TAS2R14, for which no experimental 3D structure is available. The receptors 3D models obtained by means of IFD procedure provide hypotheses on ligands’ binding and on residues involved in activation.

### Methodology and models performance

The methodology is iterative and, since the novel chemicals identified in each iteration are used to further refine the receptor models, potentially represents a perpetual procedure toward VS performance improvement for promiscuous GPCRs. Applied to agonists or antagonists, this technique allows to generate different models tailored for screening one category of ligands or the other. Each iteration comprised several steps: expansion of the set of actives through experimental methodologies; generation of receptor conformations through induced fit docking using the known ligands; conformations selection using performance evaluation through retrospective virtual screening; expansion of the set of actives through testing of structure and ligand-based predictions.

Two iterations were applied in this study, followed by a refinement step with the most updated list of ligands which was performed to generate the final models. While IT0, the original model, is able to capture 1 agonist per 28 decoys within the top 1% of the retrospective virtual screening results, our final agonist-discriminating model captures an agonist per 3.2 decoys, 8.6 times more efficient. The possibility to locate an antagonist within the top 10% of the RVS results improves by 1.7 folds. These models can be further used for prospective screening of libraries of interest.

### Ligands discovered

All of the 10 antagonists and 121 agonists – representing >60% of all the agonists here discovered – are predicted by BitterMatch to be selective for TAS2R14 over other bitter taste receptors. The antagonists could open new possibilities for lead compounds generation and provide probes in the study of TAS2Rs, but further cell-based studies will be required to prove it.

### Receptor activation

The high similarity within the binding sites of the IT3+ agonist model and the IT3-antagonist one is not surprising, given other GPCRs [57] and agonist-antagonist 2D similarities found here. N93^3.36^ is the only residue located at the bottom of the binding site that modifies its orientation between the two models, forming an H-bond with Y240^6.48^ in the IT3+ agonist model. In several GPCRs, the modulation of this interaction by binding ligands regulates the shift of the TM6 characterizing the GPCR state [42–48]. Experimental mutagenesis data support the theory of a pivotal role for these two residues in TAS2R14 [17]. Comparison between the agonist flufenamic acid and the derived antagonists suggests that the addition of a substituent in position para of ring A could alter the orientation of N93^3.36^ (Fig. 3b and 3c), determining receptor activation through the regulation of the H-bond with Y240^6.48^.

However, the homology model derived from the recently published TAS2R46 structure, as well as the AlphaFold model, do not show the 3.36-6.48 interaction, since in these models the TM6 is shifted by one helix turn compared to our models, increasing the distance between N93^3.36^ and Y240^6.48^ (Supplementary Fig. S2).

### Additional remarks

As a noteworthy member of the GPCR family, TAS2R14 is a promiscuous receptor that recognizes hundreds of different molecules. It belongs to the ~80% of GPCRs for which the experimental 3D structure has not been solved yet (December 2022) [2]. Cryo-EM has opened up new possibilities in GPCR structural biology, but stabilization of the receptor by ligands remain essential for experimental structural discovery [58, 59]. Therefore, the strategy employed within this study can be applied to a large cluster of receptors with the same challenges, facilitating ligand discovery and structural stabilization, while improving the structural knowledge of the receptor and discovering new ligands.

The systematic exploration of ligand modifications by medicinal chemistry, which is still underemployed for TAS2Rs targeting compounds, showed very promising results. Similar exploration starting from ligands other than the flufenamic acid should be investigated in the future. Among the compounds identified within this study, novel scaffolds for bitter inhibitors have been identified, providing the basis for developing more stable, effective, synthetically accessible blockers.

## Materials and Methods

### Structure based approach

#### Generation of TAS2R14 conformations: Induced Fit Docking

Generation of TAS2R14 conformations was performed for each iteration and by means of IFD.

##### Iteration 1

the initial receptor structure, IT0, was generated with I-TASSER [34]. The receptor was prepared with the protein preparation wizard (Schrödinger Release 2021-1: Protein preparation wizard, Schrödinger, LLC, New York, NY, 2021) available within the Schrödinger Suite (Schrödinger Release 2021-1, LLC, New York, NY, 2021). The TAS2R14’s active ligands known while performing this step were prepared with the ligprep tool (Schrödinger Release 2021-1: Ligprep, Schrödinger, LLC, New York, NY, 2021). The glide grid was generated based on the SiteMap results (Schrödinger Release 2021-1: SiteMap, Schrödinger, LLC, New York, NY, 2021), and all the active ligands docked to the receptor using the OPLS3e force field [60] with the IFD protocol available within Schrödinger (Schrödinger Release 2021-1: Induced fit docking protocol, Schrödinger, LLC, New York, NY, 2021). Up to 5 poses for each ligand were retained. Prime refinement was conducted for each pose of each of the known ligands, using an implicit membrane defined as spanning the transmembrane helical portions of the protein structure.

##### Iteration 2

the starting structure was the IT1+. The list of TAS2R14 ligands was updated with the flufenamic acid derivatives and the drug-like compounds retrieved within this study, as well as the active chemicals detected in iteration 1. These ligands were prepared with ligprep and the same procedure as iteration 1 was applied, with the maximum number of IFD poses limited to 3 per ligand.

##### Iteration 3

the third iteration of IFD was performed by following the same procedure as for iteration 2, by adding to the list of agonists and antagonists the active compounds from the second iteration.

A single round of IFD was also applied to the original AlphaFold model by using the complete list of active ligands from iteration 3.

#### Retrospective virtual screening and models’ selection

In each of the three iterations, evaluation of the IFD conformations VS performances was accomplished through RVS. An agonist discriminating model and an antagonist discriminating model were selected in two different docking experiments within the same set of conformations generated within that iteration. The set of actives (agonists or antagonists) was docked to the respective iteration conformations together with a set of decoys. The decoys for the agonist discriminating conformation selection differ from the one used to select the antagonist models. In both cases, they were retrieved with DUD-E http://dude.docking.org/ [61] by using as input the list of corresponding active compounds.

##### Iteration 1

a total of 10,967 decoys for the agonists and 499 for the antagonists were generated, after removals of duplicated chemicals within the same list.

##### Iteration 2

same approach employed for iteration 1 using as input for DUD-E the compounds available after iteration 1. Out of 16,280 DUD-E compounds, 1511 were randomly chosen and replaced with the inactive compounds tested in the FDA approved drugs TAS2R14 screening. A total of 538 DUD-E decoys were used for the iteration 2 antagonists model selection.

##### Iteration 3

the 16,407 decoys for the agonist model employed in iteration 3 were retrieved as in iteration 2. A total of 840 decoys were suggested by DUD-E for the third iteration of antagonist model selection. These were screened through BitterIntense, a machine learning software that, based on the chemical structure of the input molecules, predict their level of bitterness [8]. The 266 molecules with the highest scores in terms of bitter intensity were deleted from the list of decoys, leaving 574 DUD-E decoys.

The final set of active ligands and decoys employed for iteration 3 were also used for RVS of the AlphaFold models evaluation, as well as for RVS of the TAS2R14 homology model based on the TAS2R46 template (not underwent IFD) that was generated using SwissModel [50] and selecting 7XP6 as template.

As previously done for the active compounds, in each iteration the decoys were prepared with the LigPrep tool. The Glide Grid for each conformation was generated through the Cross Docking tool available within Schrödinger, by setting the grid center for each model at the centroid of the associated docked ligand. The grids were then used as input for the Schrödinger Virtual Screening Workflow tool together with the prepared active and decoy ligands in order to perform the docking on any IFD model. The Glide HTVS algorithm was selected. For all the iterations, the results of docking to each conformation were analyzed by means of the Schrödinger Enrichment Calculator tool that, giving as input the list of actives, the number of decoys and the docking file with the ligands ordered according to their docking score, calculate several parameters useful for performance evaluation. Particularly, the evaluation of the IFD models’ ability to discriminate agonists was performed by using the EF1% and the AUC parameters. The EF1% corresponds to the concentration of active ligands found within the top 1% of the results sorted by docking score compared to their concentration within the whole set of actives plus decoys. Therefore, an EF1% equal to 10 means 10 times higher chances to find an active compound within the top 1% results than randomly. The EF1% was replaced by the EF10% in the evaluation for the antagonist model due to the reduced dataset of active compounds available for the blockers. The AUC, similarly to the EF, is highly used in retrospective analysis of VS. It plots the true positive fractions versus the false positive fractions for all the compounds in the sorted dataset. EF was prioritized over AUC for models’ selection. Indeed, AUC curves summarize the ability of the method to rank the whole set of docked ligands [62], whereas in our virtual screening we focus on a subset of the screened compounds, particularly the top ranked, since our final aim is to conduct experimental test for a selection within the top ranked compounds. AUC, EF1% and EF10% values throughout the manuscript are always calculated by using the final dataset of active ligands in order to compare the different models.

#### Virtual screening

Once a model has been selected, the VS of compounds library was performed for the first two iterations. Slightly different procedures were employed for each iteration for compounds selection, in order to adapt the available data to the methodology that better fit. Again, the Enamine REAL library [36] compounds that was screened against the top IFD models was prepared with the LigPrep tool, while the receptors with the Protein Preparation tool.

##### Iteration 1

The ~23mln compounds of the Enamine REAL library were docked to the top agonist and antagonist discriminating models from iteration 1. The Virtual Screening Workflow tool was employed. This application uses the Glide program and allows to perform the docking in the 3 following steps. For any of the two receptors, the entire REAL dataset was docked using the HTVS docking algorithm. The top 20% ranked compounds according to the Glide score underwent step 2 and were docked for a second time by using the SP algorithm. The top 20% were selected for the final docking step performed with the XP algorithm. Since the agonist discriminating model from iteration 1 showed good performances in antagonists’ discrimination, the compounds to test experimentally were chosen among the 5797 ligands that reached the XP stage in both the models, in order to increase the chances to find antagonists. Glide score and visual inspection of the interactions with the receptor were used to select the top candidates for the experimental test.

##### Iteration 2

The Enamine REAL library was docked to both the best agonists and antagonists discriminating models from iteration 2 using the HTVS docking algorithm through the Virtual Screening Workflow. The top 1% of ranked compounds was then docked with the XP algorithm to the same receptors and the compounds that underwent experimental validation were selected from the top ranked compounds according to the Glide score, followed by visual inspection of the 2D chemical structure and receptor/ligand interactions.

### Structure/ligand-based approaches

#### Binary fingerprints-based hierarchical clustering

Molecular binary fingerprints allow the structural description of a molecule through a series of binary digits specifying the presence or absence of certain substructure within the compound. Fingerprint comparison is therefore useful for similarity calculation between molecules. Among the available set of fingerprints, the MolPrint2D were calculated within Canvas (Schrödinger Release 2021-3: Canvas, Schrödinger, LLC, New York, NY, 2021) for 5797 Enamine molecules reaching the final docking stage of iteration 1 and for the known active compounds. The MolPrint2D were selected because, in a fingerprint comparative study, were identified as the best ones in retrieving actives [63]. Based on the calculated fingerprint, the compounds were clustered together by using the hierarchical clustering tool available in Canvas. Enamine compounds falling within the same cluster of active molecules were selected.

#### Tanimoto similarity and clustered docking

Assuming that, in most of the cases, similar ligands bind in an analogues manner to the same receptor, the Enamine REAL library was screened to find compounds similar to known blockers.

The MolPrint2d binary fingerprints were calculated for the antagonists and for the Enamine REAL library using Schrödinger. For each antagonist, the Tanimoto score based on the fingerprint was calculated with any of the REAL compounds by means of the same software. Only compounds exceeding the threshold of 0.3 were selected for further studies.

The binding of each Enamine candidates was compared to that of the similar antagonist. For each of the known antagonists, the 10 conformations from RVS iteration 2 obtaining the top docking scores were selected. Out of these binding poses, a representative receptor/ligand structure was chosen by clustering using the conformer_cluster script available in Schrödinger. The complex with the best docking score within the most populated cluster was selected. Similarly, the Enamine compounds retrieved because of their Tanimoto score, were docked to the same 10 receptors as the antagonist they are similar to, then clustered and a representative pose selected. The selection of the compounds to test was based on a comparison of the binding pose obtained of each antagonist with the candidate molecules similar to it. Compounds with orientation within the pocket and receptor/ligands interaction similar to their antagonist counterpart were chosen. The same approach described for the antagonists was applied to the FDA drug-like compounds showing activation at the lowest concentration. Compounds similar to these 14 molecule were found through Tanimoto score within the Enamine library, and the clustered docking strategy was applied for candidates’ selection.

### Predictions with BitterMatch

BitterMatch algorithm was applied to the list of agonists and antagonists according to the protocol described in [56]. Briefly, the 3D structures of the compounds were prepared using LigPrep (Schrödinger Release 2021-1: LigPrep, Schrödinger, LLC, New York, NY, 2021) and Epik (Schrödinger Release 2021-1: Epik, Schrödinger, LLC, New York, NY, 2021) in pH = 7.0±0.5. The 3D structures were used to calculates chemical properties using Canvas (Schrödinger Release 2019–1: Canvas, Schrödinger, LLC, New York, NY, 2019) and similarities of the ligands to the training set based on linear and MolPrint2D fingerprints. All the features were inputted into the trained model to obtain the predictions.

### Principal component analysis

To explore the conformational space occupied by the known TAS2R14 ligands, we performed PCA with an in-house python script adapted from https://chem-workflows.com/articles/2019/07/02/exploring-the-chemical-space-by-pca/. A total of 43 descriptors were calculated with RDKit and the data normalized with the MinMaxScaler. The 67% of variance is explained by the top 2 PC (49% and 19%).

### Cell-based system

#### Cell culture

All the experiments were performed using HEK293T cells. Cells were cultured in plates with 10% Dulbecco’s Modified Eagle’s (DMEM) Medium, containing 10% Fetal Bovine Serum (FBS), 1% L-glutamine amino acid, and 0.2% penicillin streptomycin. Every 2-3 days, when plates were at 80% confluence, the medium was removed, and cells were transfected into a fresh growth medium. Cells were kept in the incubator, in 37 °C and 5% CO_2_.

#### Compounds

Compounds tested were obtained from Sigma-Aldrich (St. Louis, MO, USA) and Enamine (Kyiv, Ukraine). Compounds were dissolved in DMSO. Serial dilutions of compounds were prepared in the stimulation buffer of the kit (containing 50 mM LiCl to prevent IP1 degradation) at the desired working concentration on the day of the experiment (Cisbio IP-ONE-Gq KIT).

#### IP-One assay

For functional expression of the human bitter taste receptor TAS2R14, HEK293-T cells were grown at about 50-80% confluence and transiently transfected with 2 μg of a plasmid (pcDNA3.1) encoding N-terminally modified TAS2R14 (N-terminal addition of a cleavable HA-signal peptide, followed by a FLAG-tag and the first 45 amino acids of the rat somatostatin receptor 3) [33, 64] and 1 μg of a plasmid encoding the chimeric Gα_qi5_ protein (Gα_q_ protein with the last five amino acids at the C-terminus replaced by the corresponding sequence of Gα_i_, gift from The J. David Gladstone Institutes, San Francisco, CA and from Bruce Conklin (Addgene plasmid # 24501 ; http://n2t.net/addgene:24501 ; RRID:Addgene_24501) [65] employing Mirus TransIT-293 (Peqlab) in a 1:3 DNA to reagent ratio.

24 hours after transfection, the transfected cells were suspended in 10% DMEM and seeded onto a 384-well culture micro plate (Greiner). The plate was then kept in an incubator for 24 hours to obtain cell adherence. The next day, cell culture medium supernatant was removed from the plates and stimulation buffer was added to each well. Cells were then treated by the addition of the tested compounds. Plates were incubated for 150 minutes, to allow IP1 accumulation inside the cell. For investigation TAS2R14 inhibition, the cells were first pre-incubated for 30 minutes with potential inhibitor, followed by addition of 1 μM of flufenamic acid/aristolochic acid and continuing incubation for further 150 minutes.

Accumulation of second messenger was stopped by adding detection reagents (IP1-d2 conjugate and Anti-IP1cryptate TB conjugate) (dissolved in kit lysis buffer). FRET-ratios were calculated as the ratio of emission intensity of the FRET acceptor (665/10 nm) divided by the FRET donor intensity (620/10 nm). Raw FRET-ratios were normalized to buffer conditions (0%) and the maximum effect of flufenamic acid (100%) and the obtained responses were analyzed using the equation for sigmoid concentration-response curves (four-parameter) implemented in GraphPad Prism 9.3 for Windows (GraphPad Software, La Jolla, USA) to derive the maximum effect (Emax, relative to flufenamic acid/aristolochic acid) and the ligand potency (EC50). For antagonist properties the maximum effect at 1 μM flufenamic acid/aristolochic acid was normalized to 100%. Per compound, two to eleven independent experiments were performed, with each concentration in duplicate/triplicates.

### FDA-approved Drug Library Screening

#### Accumulation of inositol monophosphate (IP) as functional assay in 384-well format for TAS2R14 activation

The determination of G-protein stimulated TAS2R14 activation was measured by applying the IP-One HTRF^®^ assay (Cisbio, Codolet, France) according to the manufacturer’s protocol and as described previously [1]. In brief, HEK293T cells were grown to a confluency of approximately 80% and transiently co-transfected with the cDNAs of the hybrid G protein Gα_qi5-HA_ (Gαq protein with the last five amino acids at the C-terminus replaced by the corresponding sequence of Gα_i_; a gift from J. David Gladstone Institutes San Francisco, CA) [4] and a slightly modified human TAS2R14 (this modification included the fusion of a haemagglutinin (HA) signal followed by a Flag-tag and the first 45 amino acids of rat somatostatin receptor 3 (SSTR3) to TAS2R14 in front of its N-terminus) [5] applying TransIT-293 Mirus transfection reagent (MoBiTec, Goettingen, Germany). The next day, cells were seeded into black 384-well plates (Greiner Bio-One, Frickenhausen, Germany) and maintained for 24 h at 37 °C. After incubation with test compounds (single concentration for screening experiments; a final range of concentration from 100 pM up to 30 μM for dose-response experiments) dissolved in stimulation buffer (total V = 20 μl), at 37 °C for 150 min (unless otherwise stated), the detection reagents were added (IP1-d2 conjugate and Anti-IP1cryptate TB conjugate, each dissolved in lysis buffer), and incubation was continued at rt for 60 min. Time-resolved fluorescence resonance energy transfer (HTRF) was measured using the Clariostar plate reader (BMG, Ortenberg, Germany).

#### Description of the FDA-approved Drug Library and Control Compounds

The DiscoveryProbe™ FDA-approved drug library (L1021-100, APExBIO, USA), consisting of 1971 FDA-approved drugs with known bioactivity and safety data in humans, was purchased from Biotrend Chemikalien GmbH, Cologne, Germany. The library was provided as 10 mM stock solutions in DMSO (100 μL), arrayed in 96-well format sample storage tube boxes with screw caps (23 boxes in total) and was stored at −80°C until use. Flufenamic acid was prepared as a 10 mM stock in DMSO (Sigma-Aldrich, Steinheim, Deutschland) and was stored at −80°C until use.

#### Primary Screening Procedure of the FDA-approved Drug Library – 30 μM Final Concentration

The FDA-approved drug library was screened to assess TAS2R14 activation in an endpoint 384-well format assay, performing the IP-One HTRF® assay (Cisbio, Codolet, France). Each library compound (176 compounds/assay plate) was tested at a final single concentration of 30 μM and 0.3 % (v/v) DMSO in the assay buffer on duplicate assay plates, running in parallel. Every 384-well plate in the screen included 24 wells each for stimulated (30 μM Flufenamic acid (0.3 % (v/v) DMSO)) and unstimulated (0.3 %(v/v) DMSO) controls. 0.3 μM Flufenamic acid (0.3 % (v/v) DMSO) was added as an additional control (16 wells/assay plate). Electronically adjustable tip spacing multichannel equalizer pipettes (Thermo Fisher Scientific GmbH, Dreieich, Germany) were used for seeding cells and for pipetting, diluting and mixing steps during the screening procedure. On each day of screening, library sample storage tube boxes were thawed, equilibrated to room temperature and centrifuged before use. Then, the library compounds and controls, dissolved in DMSO, were prediluted to 2-fold final concentration into 96 deep-well plates (nerbe plus, Winsen/Luhe, Germany) containing stimulation buffer (Cisbio, Codolet, France) in a total volume of 333 μL and were subsequently transferred into the assay plates containing cells (total V = 20 μl/well). The assay was run and data was collected as described above for TAS2R14 expressing HEK293T cells.

#### Confirmation Screening Procedures - 3 μM, 1 μM and 0.3 μM Final Concentration

The resulting selected “hit” library compounds from the primary screening assay were further tested for their activity at 3 μM and thereafter at 1 μM and 0.3 μM. Therefore, hit library compounds from the primary screen (30 μM) were rearrayed and serially diluted from the 96-vial boxes stocks (10 mM in DMSO) into 96-well microplates (V-bottom, PS, clear, Greiner Bio one, Frickenhausen, Germany) in DMSO with a concentration of 1 mM and 330 μM, 100 μM, respectively. Plates were sealed and stored at −80°C until use, performing the same screening assay format and protocol as described above for TAS2R14 expressing HEK293T cells. In addition, for counter screens (specificity) mock-transfected cells (empty pcDNA 3.1) were used.

#### Data Analysis and Hit Selection Criteria

All raw data were processed and analyzed using Microsoft Excel (Microsoft Corp., Redmond, WA) and PRISM 6.0 (GraphPad Software, San Diego, CA). In each of the 384-well assay plates, the stimulated (30 μM Flufenamic acid and 0.3 % (v/v) DMSO, *n*=24, 100 % activation) and unstimulated (0.3% (v/v) DMSO, *n*=24, 0% activation) control wells were averaged (mean) and the standard deviation (S.D.) was calculated. Library compound-treated wells were normalized to the percentage activation (% activation) of those controls. Library compounds were considered as active hits when the activation (%) > (mean norm. _(unstim. control.)_ + 3 S.D. _(unstim. control._) on each of the duplicate assay plates [37, 38]. Unpaired two-tailed Student’s t-tests were used for statistical significance.

### Synthesis of flufenamic acid derivatives

The procedure for synthesis of flufenamic acid derivatives is detailed within the Supplementary File S1.

## Acknowledgments

MYN, LP, EM, and FF are members of COST action ERNEST (CA18133).

## Funding

This research was partially supported by the German Research Foundation Grants Gm 13/12 (to PG and MYN), and ISF 494/16 (to MYN). Lady Davis to FF and Excellence Fellowship from the Hebrew University Center for Nanoscience and Nanotechnology to FF, EM, and LP are gratefully acknowledged.

## Competing Interests

The authors have no relevant financial or non-financial interests to disclose.

## Author Contributions

Study conception and design: MYN and PG. Computational methodologies: FF, TD, EM. Experimental test for compounds suggested by the computational methodology: LP. Chemical synthesis of flufenamic acid derivatives: LW. Activation essays of the flufenamic acid derivatives: HH, DW. Screening of the FDA approved drugs library: ATS, DW. Analysis and interpretation of results: FF, LP, MYN, HH. Draft manuscript preparation: FF. Draft manuscript review and editing: FF, LP, TP, SL, MYN, PG.

## Data availability

The updated TAS2R14 ligands dataset and the TAS2R14 models generated within this study are included within the supplementary material and will be incorporated within the next release of BitterDB https://bitterdb.agri.huji.ac.il/dbbitter.php

## Notes

### Competing Interest Statement

The authors have declared no competing interest.

### Summary of Updates

Updated introduction, fixed typos

