## Supplementary Information for "Inhibiting a promiscuous GPCR: iterative discovery of bitter taste receptor ligands"

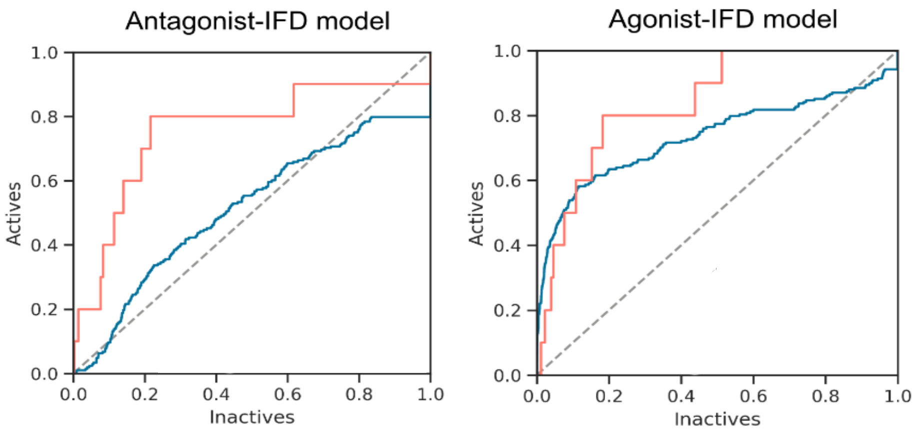


**Supplementary Fig. S1** Enrichment results for iteration 1. The antagonist model prioritized antagonists (orange curves) only over decoys (inactives), the agonist model prioritized both agonists (blue curves) and antagonists over decoys.

**Supplementary Table S1** Functional properties of TAS2R14 activated by the agonists derived from the computational study.

| | agonist effect[a] | | |
|---|---|---|---|
| | $EC_{50}$ [µM][b] | $E_{max}$ [%][c] | N[d] |
| LF3 | 909 ± 778 | 77 ± 1 | 3 |
| LF6 | 5.1 ± 2 | 23 ± 6 | 2 |
| LF9 | 0.084 | 23.7 | 1 |
| LF11 | 5.2±5 | 23 ± 20 | 2 |
| LF25 | 16.1 ± 11 | 63 ± 3 | 3 |
| LF26 | 2.8 ± 1.5 | 67 ± 1 | 3 |

[a]Measurement of G-protein signaling was performed applying the IP-One assay® (Cisbio) in HEK293T cells transiently co-transfected with the human TAS2R14 receptor and the hybrid G-protein G□$_{qi}$. [b]Potency of TAS2R14 activation as mean value in µM±SEM. [c]Maximum efficacy in %±SEM relative to the full effect of flufenanmic acid. [d] Number of individual experiments all performed in triplicates.

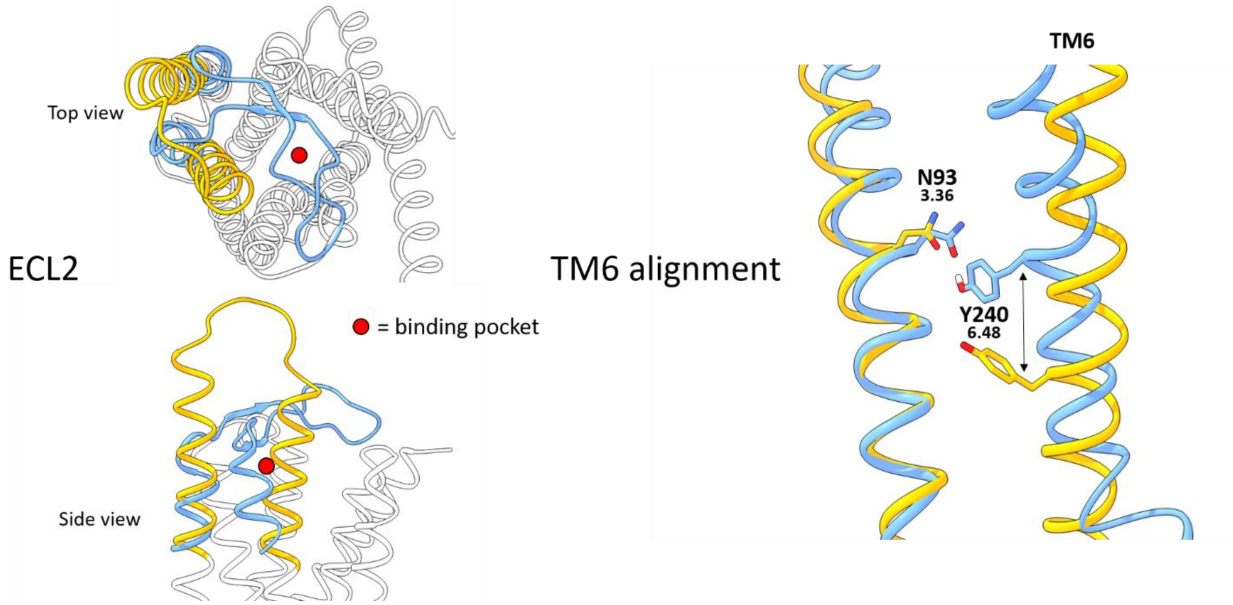


**Supplementary Fig. S2** Structural alignment between AlphaFold (orange) and IT3+ (blue) showing the differences between the two models. Within the active structure of the TAS2R46 template used to generate a homology model of TAS2R14 (not shown within this picture), the ECL2 is not solved. Therefore, our model is also missing the same loop. TM6 structural alignment of the homology model based on TAS2R46 overlap with the AlphaFold one.
