## Supplementary File 1 for "Inhibiting a promiscuous GPCR: iterative discovery of bitter taste receptor ligands"

### Supplementary File 1: synthesis of flufenamic acid derivatives

#### Inhibiting a promiscuous GPCR: iterative discovery of bitter taste receptor ligands

Fabrizio Fierro[1], Lior Peri[1], Harald Hubner[2], Alina Tabor-Schkade[2], Lukas Waterloo[2], Stefan Löber[2], Tara Pfeiffer[2], Dorothee Weikert[2], Tamir Dingjan[1], Eitan Margulis[1], Peter Gmeiner[2], Masha Y Niv[1*]

1. The Institute of Biochemistry, Food Science and Nutrition, Robert H. Smith Faculty of Agriculture, Food and Environment, The Hebrew University of Jerusalem, Rehovot, Israel

2. Department of Chemistry and Pharmacy, Medicinal Chemistry, Friedrich-Alexander-Universität Erlangen-Nürnberg, Nikolaus-Fiebiger-Str. 10, 91058 Erlangen, Germany

##### Chemistry:

Synthesis procedures were conducted using standard equipment and devices. Reactions were performed under inert atmosphere using argon gas if not stated otherwise. Dry solvents were used for reactions if not stated otherwise.

##### Solvents

Solvents were purchased in the highest available purity grade from FLUKA, ACROS, SIGMA-ALDRICH, FISHER SCIENTIFIC and VWR and used without further purification. Dry solvents for synthetic procedures were purchased and stored, whenever possible, in sealed septum bottles over molecular sieves.

##### Chemicals

Chemicals for synthetic procedures were purchased from FLUKA, ACROS, SIGMA-ALDRICH, ALFA-AESAR, TCI, FISHER SCIENTIFIC, ABCR, CHEMPUR FEINCHEM, IRIS BIOTECH and THERMO FISHER in the highest available purity grade and used without further purification.

##### NMR-Spectroscopy

NMR spectra were either recorded independently in an open access mode or by Anke Seitz, Annette Hofmann, and Sonja Burkhardt under the supervision of Dr. Jürgen Einsiedel. NMR spectra were obtained on a Bruker Avance 400 ($^{1}$H at 400 MHz, $^{13}$C (DEPTQ) at 101 MHz, $^{19}$F at 377 MHz) or a Bruker Avance 600 ($^{1}$H at 600 MHz, $^{13}$C (DEPTQ) at 151 MHz, $^{19}$F at 565 MHz) spectrometer at 298K using the solvents indicated. Chemical shifts are reported relative to TMS (for $^{1}$H and $^{13}$C) or $CCl_3F$ (for $^{19}$F NMR) or the residual solvent peak.

##### High-Resolution Mass Spectrometry

ESI-TOF high mass accuracy and resolution experiments were performed on an AB Sciex Triple TOF660 Sciex, on a Bruker maXis MS or a Bruker timsTOF Pro.

##### Flash Column Chromatography

Purification by flash chromatography was performed using Silica Gel 60 (40-63 µm mesh) from Merck as a stationary phase.

**Thin Layer Chromatography**

TLC analyses were performed using Merck 60 F254 aluminum sheets and the spots were visualized under UV light (254 nm) and with reagents such as $KMnO_4$ or ninhydrin solutions.

**HPLC-MS**

Analytical LCMS was performed on:

Thermo Scientific Dionex Ultimate 3000 HPLC system using DAD detection (230 nm; 254 nm) equipped with either a Kinetex 2.6u mesh C8 100A (2.1 x 75 mm, 2.6 µm) HPLC column or a Zorbax Eclipse XDB-C8 (4.6 x 150 mm, 5 µm) HPLC column, using mass detection on a BRUKER amaZon SL mass spectrometer using ESI or APCI ionization source.

or

Waters e2695 HPLC system using DAD detection (220 nm; 240 nm) equipped with a C18 (2.1 x 50 mm x 2.6 µm) column, using a quantom diode array detector for mass analysis.

**Preparative RP-HPLC**

Purification by preparative RP-HPLC was performed on AGILENT 1260 Preparative Series equipped with a VWD detector (230 nm; 254 nm) using a Zorbax Eclipse XDB-C8 21.2 x150 mm column with 5 µm particles [C8], flow rate 10 mL/min, employing eluent systems as specified below.

**Analytical RP-HPLC**

HPLC purity analyses were performed with an Agilent binary gradient system using UV detection ($\lambda$ = 220, 254, and 280 nm) in combination with ChemStation software. A Zorbax Eclipse XDB-C8 (4.6 mm × 150 mm, 5 µm) column was used with a flow rate of 0.5 mL/min in reversed-phase mode.

eluent system 1: methanol / $H_2O$ + 0.1% $HCO_2H$

- gradient: 10% for 3 min, 10% to 100% in 15 min, 100% for 6 min, 100% to 5% in 3 min, 10% for 3 min

eluent system 2: acetonitrile / $H_2O$+0.1% $HCO_2H$

- gradient: 5% for 3 min, 5% to 95% in 15 min, 95% for 6 min, 95% to 5% in 3 min, 5% for 3 min

#### Syntheses of antagonists:

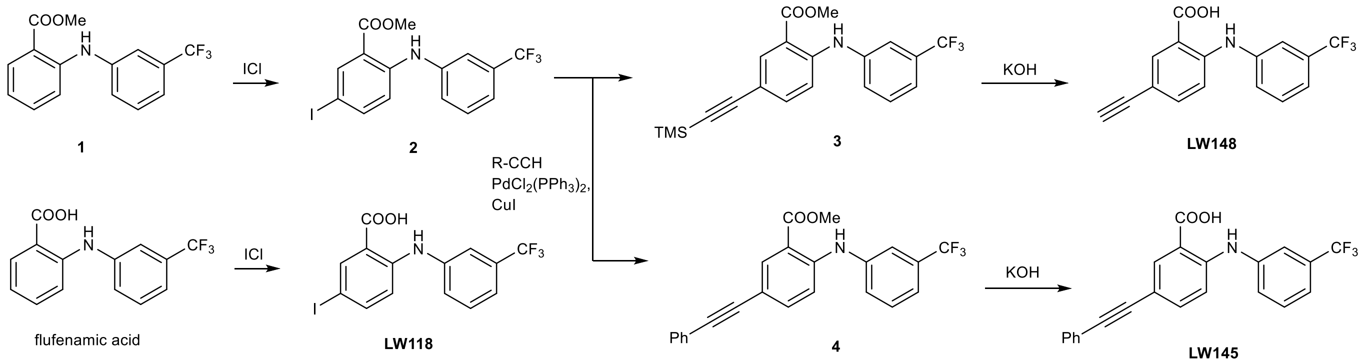


**Scheme 1**: Synthesis of target compounds **LW118**, **LW145** and **LW148**.

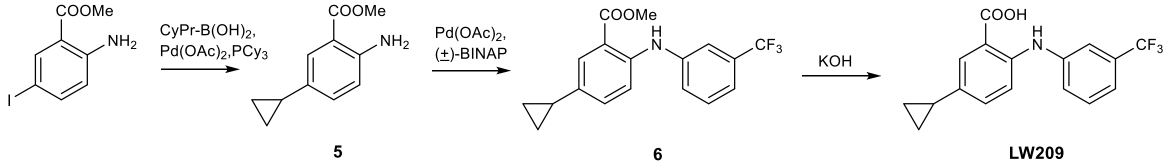


**Scheme 2**: Synthesis of target compound **LW209**.

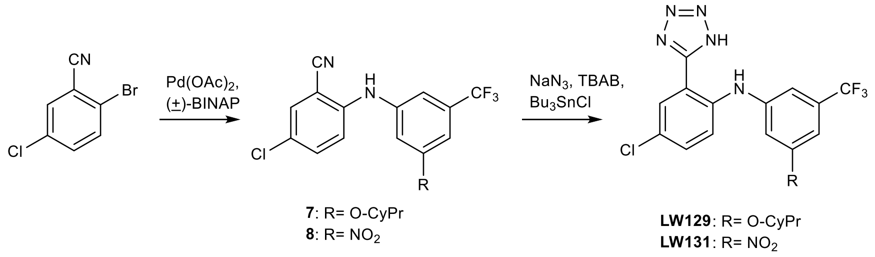


**Scheme 3**: Synthesis of target compounds **LW129** and **LW131**.

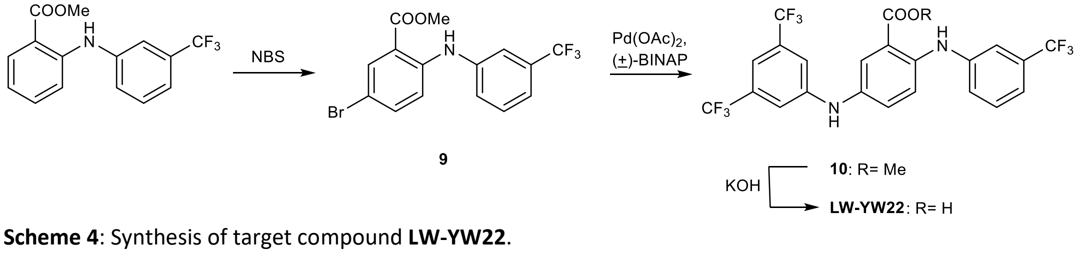


**Scheme 4**: Synthesis of target compound **LW-YW22**.

**Experimental procedures for the synthesis of antagonists:**

**5-Iodo-2-((3-(trifluoromethyl)phenyl)amino)benzoic acid (LW118)**

In a microwave tube, flufenamic acid (103 mg, 366 µmol) was dissolved in a mixture of $CH_2Cl_2$/AcOH (1mL). Then, ICl dissolved in $CH_2Cl_2$ (3.5 mL) was added to the mixture via syringe. The mixture was then stirred at rt for 15h. After the reaction was finished, solid $Na_2S_2O_5$ was added and the mixture was stirred until it turned orange. After filtration and evaporation of solvents, final purification by flash chromatography ($CH_2Cl_2$/MeOH 50:1 + 0.1% HCOOH) yielded 103 mg (253 µmol, 69%) of a greenish solid.

TLC: $R_f$ = 0.17 ($CH_2Cl_2$/MeOH 50:1 + 0.1% HCOOH)

ESI-MS: m/z 407.82 $[M+H]^+$

$^1$H NMR: (400 MHz, DMSO-$d_6$) δ 13.45 (s, 1H), 9.66 (s, 1H), 8.16 (d, *J* = 2.3 Hz, 1H), 7.71 (dd, *J* = 8.8, 2.2 Hz, 1H), 7.56 (d, *J* = 2.8 Hz, 3H), 7.42 – 7.35 (m, 1H), 7.11 (d, *J* = 8.9 Hz, 1H).

$^{13}$C NMR: (101 MHz, DMSO-$d_6$) δ 168.7, 145.7, 142.6, 141.8, 140.1, 131.1, 130.8 (q, *J* = 31.6 Hz), 124.9, 124.5 (q, *J* = 272.5 Hz), 119.7 (q, *J* = 3.8 Hz), 118.0 – 117.3 (m), 116.9, 80.3.

HR-MS (ESI): m/z $[M+H]^+$ found 407.9703, calcd. 407.9703 for $C_{14}H_{10}F_3INO_2$.

Purity: λ = 254: 96.9%, $t_R$ = 21.7 min (eluent system 1)

λ = 254: >99.9%, $t_R$ = 19.6 min (eluent system 2).

**Methyl 5-iodo-2-((3-(trifluoromethyl)phenyl)amino)benzoate (2)**

In a microwave tube, commercially available **1** (765 mg, 2.59 mmol) was dissolved in a mixture of $CH_2Cl_2$/AcOH (10mL). Then, ICl dissolved in $CH_2Cl_2$ (5 mL) was added to the mixture via syringe. The mixture was then stirred at rt for 16h. After the reaction was finished, solid $Na_2S_2O_5$ was added to the mixture and it was stirred until it turned colorless. After filtration and evaporation of solvents, final purification by flash chromatography (isohexane/$CH_2Cl_2$ 10:1) yielded 103 mg (253 µmol, 69%) of a greenish solid.

TLC: $R_f$ = 0.40 (isohexane/$CH_2Cl_2$ 10:1)

ESI-MS: m/z 421.79 $[M+H]^+$

$^1$H NMR: (600 MHz, $CDCl_3$) δ 9.56 (s, 1H), 8.27 (d, *J* = 2.2 Hz, 1H), 7.58 (dd, *J* = 8.9, 2.2 Hz, 1H), 7.47 – 7.43 (m, 2H), 7.39 – 7.36 (m, 1H), 7.35 – 7.32 (m, 1H), 7.02 (d, *J* = 8.9 Hz, 1H), 3.91 (s, 3H).

$^{13}$C NMR: (151 MHz, $CDCl_3$) δ 167.6, 146.5, 142.5, 140.9, 140.1, 132.0 (q, *J* = 32.4 Hz), 130.0, 125.1, 123.8 (q, *J* = 272.4 Hz), 120.3 (q, *J* = 3.8 Hz), 118.6 (q, *J* = 3.8 Hz), 116.2, 114.8, 78.4, 52.2.

**Methyl 2-((3-(trifluoromethyl)phenyl)amino)-5-((trimethylsilyl)ethynyl)benzoate (3)**

In a microwave tube, **2** (113 mg, 268 µmol), $PdCl_2(PPh_3)_2$ (9.00 mg, 13.1 µmol) and CuI (13.0 mg, 68.3 µmol) were dissolved in diethylamine (3 mL). The mixture was purged with argon while TMS-acetylene (72.3 µL, 523 µmol) was added. The tube was sealed with a crimp cap with rubber septum and the reaction was stirred at rt for 23h. After evaporation of solvent, purification by flash chromatography (isohexane/$CH_2Cl_2$ 6:1) yielded 80.0 mg (204 µmol, 78%) of a yellow oil.

TLC: $R_f$ = 0.22 (isohexane/$CH_2Cl_2$ 6:1)

ESI-MS: m/z 391.94 $[M+H]^+$

$^1H$ NMR: (400 MHz, $CDCl_3$) δ 9.75 (s, 1H), 8.14 (dd, *J* = 2.1, 0.4 Hz, 1H), 7.51 – 7.38 (m, 5H), 7.36 – 7.33 (m, 1H), 7.16 (d, *J* = 8.8 Hz, 1H), 3.92 (s, 3H), 0.25 (s, 9H).

$^{13}C$ NMR: (151 MHz, $CDCl_3$) δ 168.4, 147.0, 141.0, 137.6, 136.0, 132.2 (q, *J* = 32.4 Hz), 130.2, 125.5, 124.0 (q, *J* = 272.5 Hz), 120.6 (q, *J* = 3.9 Hz), 119.1 (q, *J* = 3.8 Hz), 114.0, 112.8, 112.6, 104.7, 92.9, 52.3, 0.2.

**Methyl 5-(phenylethynyl)-2-((3-(trifluoromethyl)phenyl)amino)benzoate (4)**

In a microwave tube, **2** (113 mg, 268 µmol), $PdCl_2(PPh_3)_2$ (10.0 mg, 14.3 µmol) and CuI (12.0 mg, 63.0 µmol) were dissolved in diethylamine (3 mL). The mixture was purged with argon while phenylacetylene (57.4 µL, 523 µmol) was added. The tube was sealed with a crimp cap with rubber septum and the reaction was stirred at rt for 23h. After evaporation of solvent, purification by flash chromatography (isohexane/$CH_2Cl_2$ 6:1) yielded 80.0 mg (204 µmol, 78%) of a yellow oil.

TLC: $R_f$ = 0.35 (isohexane/EtOAc 20:1)

ESI-MS: m/z 395.94 $[M+H]^+$

$^1H$ NMR: (400 MHz, $CDCl_3$) δ 9.77 (s, 1H), 8.20 (dd, *J* = 2.1, 0.4 Hz, 1H), 7.53 – 7.48 (m, 4H), 7.48 – 7.44 (m, 1H), 7.44 – 7.40 (m, 1H), 7.37 – 7.30 (m, 4H), 7.22 (d, *J* = 8.8 Hz, 1H), 3.94 (s, 3H).

$^{13}C$ NMR: (101 MHz, $CDCl_3$) δ 168.3, 146.6, 140.8, 137.1, 135.4, 132.5 – 131.3 (m), 131.4, 130.0, 128.3, 128.0, 125.3 – 125.2 (m), 123.8 (q, *J* = 272.4 Hz), 123.4, 120.4 (q, *J* = 3.8 Hz), 118.9 (q, *J* = 3.8 Hz), 113.9, 112.7, 112.5, 88.8, 88.2, 52.1.

**GP1: General procedure for the hydrolysis of phenylamino benzoates:**

In a microwave tube, the respective phenylamino benzoate was dissolved in a mixture of EtOH/$H_2O$ or EtOH/THF/$H_2O$. After addition of KOH or 2 N NaOH, the tube was sealed with a crimp cap with rubber septum and the mixture was stirred at rt or reflux. After completion, the mixture was cooled to rt and diluted with $H_2O$. The organic solvent was evaporated and the resulting aqueous fraction was acidified with 2 N HCl to pH 1. The formed precipitate was filtered and washed with $H_2O$ to give the final compound. In case of impurities, further means of purification are indicated for the respective compound.

**GP2: General procedure for the coupling of anilines with aryl halides:**

In a microwave tube, the respective aryl halide (if solid), aniline/aminopyridine/aminopyrimidine (if solid), $Pd(OAc)_2$ and (±)BINAP were dissolved in dry toluene. The mixture was purged with argon and the respective aryl halide (if liquid) and aniline/aminopyridine/aminopyrimidine (if liquid) were added under argon flow. After addition of $Cs_2CO_3$, the resulting mixture was purged for further two minutes with argon, sealed with a crimp cap with rubber septum and stirred at 120°C. After completion, the mixture was diluted with EtOAc, washed with 2 N HCl (3x), brine (3x), dried over $MgSO_4$, filtered and evaporated. The product was purified by flash chromatography.

**GP3: General procedure for the synthesis of tetrazoles from phenylamino benzonitriles:**

In a microwave tube, the respective phenylamino benzonitrile, TBAB, $NaN_3$ and $Bu_3SnCl$ were dissolved in dry DMF or toluene, purged with argon, sealed with a crimp cap with rubber septum and stirred at 165°C (DMF) or 120°C (toluene). After completion, the reaction was cooled to rt and transferred into a separatory funnel. Then, an excess of $H_2O$ was added and extracted with EtOAc (3x). The combined organic fractions were then washed with 2 N HCl (3x), brine (3x), dried over $MgSO_4$, filtered and evaporated. To remove traces of tin impurities, the resulting residue was then taken up in ACN and 2 N HCl and stirred for 1h. The organic solvent was evaporated and the precipitate was filtered and washed with $H_2O$ and isohexane. Final purification of the compound was achieved by column chromatography or preparative HPLC.

**5-(Phenylethynyl)-2-((3-(trifluoromethyl)phenyl)amino)benzoic acid (LW145)**

Compound **LW145** was synthesized according to the before mentioned general procedure GP1 using **LW138** (59.0 mg, 149 µmol) and KOH (24.0 mg, 428 µmol) in a mixture of EtOH (4 mL) and $H_2O$ (1 mL). The mixture was stirred at 80°C for 1h. Precipitation and filtration yielded 53.0 mg (135 µmol, 67%) of a yellowish solid.

ESI-MS: m/z 381.92 $[M+H]^+$

$^1$H NMR: (400 MHz, DMSO-$d_6$) δ 13.53 (s, 1H), 9.94 (s, 1H), 8.08 (d, $J$ = 2.1 Hz, 1H), 7.64 – 7.52 (m, 6H), 7.47 – 7.38 (m, 4H), 7.28 (d, $J$ = 8.8 Hz, 1H).

$^{13}$C NMR: (151 MHz, DMSO-$d_6$) δ 168.8, 145.9, 140.9, 136.6, 135.1, 131.1, 130.6, 130.3 (q, $J$ = 31.8 Hz), 128.7, 128.4, 125.1, 124.0 (q, $J$ = 272.5 Hz), 122.5, 119.7 (q, $J$ = 3.9 Hz), 117.8 (q, $J$ = 3.9 Hz), 114.5, 114.0, 111.7, 88.9, 88.0.

HR-MS (ESI): m/z $[M+H]^+$ found 382.1055, calcd. 382.1049 for $C_{22}H_{15}F_3NO_2$.

Purity: λ = 254: 97.5%, $t_R$ = 21.9 min (eluent system 1)

λ = 254: >99.9%, $t_R$ = 20.4 min (eluent system 2).

**5-Ethynyl-2-((3-(trifluoromethyl)phenyl)amino)benzoic acid (LW148)**

Compound **LW148** was synthesized according to the before mentioned general procedure GP1 using **LW137** (45.0 mg, 115 µmol) and KOH (29.0 mg, 517 µmol) in a mixture of EtOH (4 mL), THF (1 mL) and $H_2O$ (1 mL). The mixture was stirred at 80°C for 1h. Precipitation and filtration yielded 30.0 mg (98.3 µmol, 85%) of a beige solid.

ESI-MS: m/z 305.84 $[M+H]^+$

$^1$H NMR: (400 MHz, DMSO-$d_6$) δ 13.49 (s, 1H), 9.88 (s, 1H), 7.98 (d, *J* = 2.1 Hz, 1H), 7.59 (d, *J* = 2.9 Hz, 3H), 7.51 (dd, *J* = 8.7, 2.2 Hz, 1H), 7.46 – 7.40 (m, 1H), 7.23 (d, *J* = 8.7 Hz, 1H), 4.07 (s, 1H).

$^{13}$C NMR: (101 MHz, DMSO-$d_6$) δ 168.7, 146.0, 140.8, 136.9, 135.2, 130.5, 130.2 (q, *J* = 31.7 Hz), 125.0 – 124.9 (m), 123.9 (q, *J* = 272.4 Hz), 119.6 (q, *J* = 4.0 Hz), 117.7 (q, *J* = 3.9 Hz), 114.4, 113.7, 111.0, 82.8, 79.2.

HR-MS (ESI): m/z $[M+H]^+$ found 306.0738, calcd. 306.0736 for $C_{16}H_{11}F_3NO_2$.

Purity: λ = 254: >99.9%, $t_R$ = 20.9 min (eluent system 1)

λ = 254: 95.7%, $t_R$ = 18.6 min (eluent system 2).

**Methyl 2-amino-5-cyclopropylbenzoate (5)**[1]

In a microwave tube, methyl 2-amino-5-iodobenzoate (499 mg, 1.80 mmol), cyclopropylboronic acid (215 mg, 2.50 mmol), $PCy_3$ (53.0 mg, 189 µmol) and $K_3PO_4$ (1.34 g, 6.30 mmol) were dissolved in toluene (10 mL) and $H_2O$ (0.5 mL). After purging the mixture with argon, $Pd(OAc_2)_2$ (23.0 mg, 102 µmol) was added under constant argon flow. The tube was sealed with a crimp cap with rubber septum and stirred at 100°C for 3 days. After cooling to rt, the reaction was diluted with EtOAc, washed with 2 N HCl (3x), brine (3x), dried over $MgSO_4$, filtered and evaporated. Flash chromatography (isohexane/EtOAc 10:1) yielded an impure product, which was taken to the next step without further purification.

ESI-MS: m/z 191.95 $[M+H]^+$

$^1$H NMR: (400 MHz, $CDCl_3$) δ 7.60 (ddd, *J* = 2.3, 0.5 Hz, 1H), 7.04 (ddd, *J* = 8.5, 2.3, 0.5 Hz, 1H), 6.63 (d, *J* = 8.4 Hz, 1H), 5.74 (s, 2H), 3.87 (s, 3H).

$^{13}$C NMR: (101 MHz, $CDCl_3$) δ 168.5, 147.8, 132.4, 131.9, 128.2, 117.2, 111.0, 51.6, 14.5, 7.0.

**Methyl 5-cyclopropyl-2-((3-(trifluoromethyl)phenyl)amino)benzoate (6)**

Compound **6** was synthesized according to the before mentioned general procedure GP2 using **6** (110 mg, 575 µmol), 1-bromo-3-(trifluoromethyl)benzene (96.5 µL, 690 µmol), $Pd(OAc)_2$ (7.00 mg, 31.2 µmol), (±)-BINAP (34.0 mg, 54.6 µmol) and $Cs_2CO_3$ (267 mg, 819 µmol) in 5 mL dry toluene. The reaction was finished after 1h. Flash chromatography (isohexane/EtOAc 20:1) yielded an impure product, which was taken to the next step without further purification.

**ESI-MS:** m/z 336.10 [M+H]$^+$

**5-Cyclopropyl-2-((3-(trifluoromethyl)phenyl)amino)benzoic acid (LW209)**

Compound **LW209** was synthesized according to before mentioned general procedure GP1 using crude **LW206** and KOH (54.0 mg, 962 µmol) in a mixture of EtOH (2.5 mL) and $H_2O$ (0.5 mL). The mixture was stirred at 80°C for 1h. Precipitation, filtration and purification by preparative HPLC (gradient of 20% to 95% ACN in $H_2O$ with 0.3% HCOOH) yielded 10.0 mg (31.0 µmol, 3% over 3 steps) of a yellow solid.

ESI-MS: m/z 322.08 [M+H]$^+$

$^1$H NMR: (400 MHz, DMSO-$d_6$) δ 13.20 (s, 1H), 9.47 (s, 1H), 7.67 (d, *J* = 2.3 Hz, 1H), 7.54 – 7.44 (m, 3H), 7.30 – 7.23 (m, 2H), 7.19 (dd, *J* = 8.6, 2.3 Hz, 1H), 1.96 – 1.86 (m, 1H), 0.95 – 0.88 (m, 2H), 0.65 – 0.57 (m, 2H).

$^{13}$C NMR: (101 MHz, DMSO-$d_6$) δ 169.3, 142.5, 134.2, 131.0, 130.4, 130.1 (q, *J* = 31.5 Hz), 128.6, 124.0 (q, *J* = 272.4 Hz), 122.5, 117.62 (q, *J* = 4.0 Hz), 116.0, 115.2 – 115.0 (m), 115.1, 14.1, 8.5.

HR-MS (ESI): m/z [M+H]$^+$ found 322.1049, calcd. 322.1049 for $C_{17}H_{15}F_3NO_2$.

Purity: λ = 254: >99.9%, $t_R$ = 21.7 min (eluent system 1)

λ = 254: >99.9%, $t_R$ = 19.4 min (eluent system 2).

**5-Chloro-2-((3-cyclopropoxy-5-(trifluoromethyl)phenyl)amino)benzonitrile (7)**

Compound **7** was synthesized according to a slightly modified general procedure GP2 using 2-bromo-5-chlorobenzonitrile (188 mg, 869 µmol), 3-cyclopropoxy-5-(trifluoromethyl)aniline (184 mg, 847 µmol), $Pd(OAc)_2$ (9.00 mg, 40.1 µmol), (±)-BINAP (39.0 mg, 62.6µmol) and $Cs_2CO_3$ (341 mg, 1.05 mmol) in 5 mL dry toluene at 80°C. The reaction was finished after 23h. Purification by flash chromatography (isohexane/EtOAc 15:1) yielded 205 mg (581 µmol, 82%) of an off white solid.

TLC: $R_f$ = 0.33 (isohexane/EtOAc 15:1)

ESI-MS: m/z 352.91 [M+H]$^+$

$^1$H NMR: (600 MHz, $CDCl_3$) δ 7.51 (d, *J* = 2.5 Hz, 1H), 7.39 (ddd, *J* = 9.1, 2.5, 0.6 Hz, 1H), 7.22 (d, *J* = 9.0 Hz, 1H), 7.06 – 7.04 (m, 1H), 7.01 – 6.99 (m, 1H), 6.96 (dd, *J* = 2.1 Hz, 1H), 6.38 (s, 1H), 3.76 (dddd, *J* = 5.9, 3.1 Hz, 1H), 0.86 – 0.78 (m, 4H).

$^{13}$C NMR**:** (151 MHz, $CDCl_3$) δ 160.6, 144.7, 141.6, 134.4, 133.0 (q, *J* = 32.7 Hz), 132.3, 125.2, 123.6 (q, *J* = 272.7 Hz), 116.7, 116.0, 110.4, 109.9 (q, *J* = 3.8 Hz), 107.6 (q, *J* = 3.9 Hz), 101.1, 51.4, 6.3.

**2-((3-Nitro-5-(trifluoromethyl)phenyl)amino)benzonitrile (8)**

Compound **8** was synthesized according to the general procedure GP2 using 2-bromo-benzonitrile (151 mg, 829 µmol), 3-nitro-5-(trifluoromethyl)aniline (205 mg, 995 µmol), $Pd(OAc)_2$ (12.0 mg, 53.5 µmol), (±)-BINAP (42.0 mg, 67.5µmol) and $Cs_2CO_3$ (377 mg, 1.16 mmol) in 5 mL dry toluene. The reaction was finished after 4.5h. Purification by flash chromatography (isohexane/EtOAc 10:1) yielded 205 mg (667 µmol, 80%) of a deep yellow solid.

TLC: $R_f$ = 0.24 (isohexane/EtOAc 15:1)

ESI-MS: m/z 329.91 $[M+Na]^+$

$^1$H NMR: (400 MHz, $CD_3CN$) δ 8.04 (dd, *J* = 2.2 Hz, 1H), 8.01 – 7.99 (m, 1H), 7.74 (ddd, *J* = 7.8, 1.6, 0.5 Hz, 1H), 7.67 – 7.65 (m, 1H), 7.65 – 7.59 (m, 1H), 7.52 (s, 1H), 7.46 – 7.41 (m, 1H), 7.23 (ddd, *J* = 7.6, 1.0 Hz, 1H).

$^{13}$C NMR: (101 MHz, $CD_3CN$) δ 149.6, 145.2, 143.9, 134.5, 134.1, 132.1 (q, *J* = 33.7 Hz), 123.9, 123.2 (q, *J* = 272.1 Hz), 120.3, 119.3 (q, *J* = 3.7 Hz), 116.8, 114.9 – 114.9 (m), 112.2 (q, *J* = 4.0 Hz), 104.7.

**4-Chloro-*N*-(3-cyclopropoxy-5-(trifluoromethyl)phenyl)-2-(1*H*-tetrazol-5-yl)aniline (LW129)**

Compound **LW129** was synthesized according to the before mentioned general procedure GP3 using **7** (84.0 mg, 238 µmol), TBAB (8.00 mg, 24.8 µmol), $NaN_3$ (96.0 mg, 1.48 mmol) and $Bu_3SnCl$ (388 µL, 1.43 mmol) in 2 mL dry DMF. The reaction was finished after 14h. Purification by flash chromatography ($CH_2Cl_2$/MeOH 50:1 + 0.1% HCOOH) yielded 31.0 mg (78.3 µmol, 33%) of a white solid.

TLC: $R_f$ = 0.11 ($CH_2Cl_2$/MeOH 50:1 + 0.1% HCOOH)

ESI-MS: m/z 395.89 $[M+H]^+$

$^1$H NMR: (600 MHz, $CD_3CN$) δ 8.97 (s, 1H), 7.89 (d, *J* = 2.4 Hz, 1H), 7.47 (d, *J* = 8.9 Hz, 1H), 7.43 (dd, *J* = 9.0, 2.4 Hz, 1H), 7.11 (dd, *J* = 2.2 Hz, 1H), 7.09 – 7.06 (m, 1H), 6.99 – 6.97 (m, 1H), 3.83 (dddd, *J* = 6.0, 2.9 Hz, 1H), 0.83 – 0.79 (m, 2H), 0.73 – 0.69 (m, 2H).

$^{13}$C NMR: (151 MHz, $CD_3CN$) δ 161.7, 144.8, 142.0, 133.8 – 132.3 (m), 129.8, 125.8, 125.1 (q, *J* = 271.8 Hz), 120.1, 113.7, 110.5, 110.1 (q, *J* = 4.1 Hz), 107.1 (q, *J* = 3.9 Hz), 52.3, 6.7.

HR-MS (ESI): m/z $[M+H]^+$ found 396.0838, calcd. 396.0833 for $C_{17}H_{14}ClF_3N_5O$.

Purity: λ = 254: 97.2%, $t_R$ = 22.1 min (eluent system 1)

λ = 254: 98.2%, $t_R$ = 19.8 min (eluent system 2).

**4-Chloro-*N*-(3-nitro-5-(trifluoromethyl)phenyl)-2-(1*H*-tetrazol-5-yl)aniline (LW131)**

Compound **LW131** was synthesized according to a slightly modified general procedure GP3 using **8** (70.0 mg, 205 µmol), TBAB (9.00 mg, 27.9 µmol), $NaN_3$ (40.0 mg, 615 µmol) and $Bu_3SnCl$ (167 µL, 616 µmol) in 2 mL dry toluene at 120°C. The reaction was finished after 16h. Purification by flash chromatography ($CH_2Cl_2$/MeOH 50:1 + 0.1% HCOOH) and subsequent preparative HPLC (gradient of 20% to 95% ACN in $H_2O$ with 0.3% HCOOH) yielded 25.0 mg (65.0 µmol, 26%) of an off white solid.

TLC: $R_f$ = 0.13 ($CH_2Cl_2$/MeOH 50:1 + 0.1% HCOOH)

ESI-MS: m/z 384.82 $[M+H]^+$

$^1H$ NMR: (400 MHz, $CD_3CN$) δ 9.14 (s, 1H), 8.17 – 8.13 (m, 1H), 8.02 – 7.98 (m, 2H), 7.78 – 7.75 (m, 1H), 7.56 – 7.46 (m, 2H).

$^{13}C$ NMR: (151 MHz, $CD_3CN$) δ 150.7, 145.9, 140.3, 133.3 (q, *J* = 33.7 Hz), 132.8, 130.2, 127.9, 127.1 – 121.2 (m), 116.8, 116.5, 113.6 (q, *J* = 3.9 Hz).

HR-MS (ESI): m/z $[M+H]^+$ found 385.0425, calcd. 385.0422 for $C_{14}H_9ClF_3N_6O_2$.

Purity: λ = 254: 99.1%, $t_R$ = 21.5 min (eluent system 1)

λ = 254: 98.8%, $t_R$ = 19.0 min (eluent system 2).

**Methyl-5-bromo-2-((3-(trifluormethyl)phenyl)amino)benzoate (9)**

Diisopropylamine (0.09 ml; 65 mg; 0,64 mmol) was added to a solution of commercially available **1** (250 mg; 0.849 mmol) and NBS (304 mg; 2.58 mmol) in $CH_2CL_2$ (20 mL). After being stirred at room temperature for 65h, the solution washed with water (3x 20 mL). The organic layer was dried with $Na_2SO_4$ and evaporated under reduced pressure. Flash chromatography (isohexane 100%) yielded **9** (302 mg, 0.81 mmol, 95%) as a yellow liquid.

TLC: $R_f$ = 0.17 (isohexane)

ESI-MS: m/z 375.93 $[M+H]^+$

$^1H$ NMR: (600 MHz, $CDCl_3$) δ 9.54 (s, 1H), 8.10 (d, *J* = 2.5 Hz, 1H), 7.45 (d, *J* = 3.3 Hz, 2H), 7.43 (dd, *J* = 9.0, 2.4 Hz, 1H), 7.38 (d, *J* = 8.4 Hz, 1H), 7.33 (d, *J* = 7.7 Hz, 1H), 7.26 (s, 1H), 7.14 (d, *J* = 9.0 Hz, 1H), 5.30 (s, 2H), 3.92 (s, 3H).

$^{13}C$ NMR: (101 MHz, $CDCl_3$) δ 167.8, 145.9, 141.0, 137.0, 134.1, 130.1, 125.0, 120.3, 118.6, 116.0, 114.2, 109.6, 52.3, 29.7.

**Methyl-5-((3,5-bis(trifluormethyl)phenyl)amino)-2-((3-(trifluormethyl)phenyl)amino)benzoate (10)**

A solution of **9** (70.8 mg; 0.189 mmol), $Pd(OAc)_2$ (2.4 mg; 11 µmol), (±)-BINAP (9.8 mg; 16 µmol) and 3,5-bis(trifluoromethyl)aniline (75 µl; 110 mg; 480 µmol) in dry toluene (5 mL) was flushed with argon for 5 min. After the addition of $Cs_2CO_3$ (87.8 mg; 0.269 mmol) the mixture was flushed again with argon and subsequently stirred for 66h at reflux temperature. After evaporation of the solvent, the crude product was purified by flash chromatography (isohexane/EtOAc 15:1) to yield **10** (65.5 mg, 0.125 mmol, 66%) as a yellow liquid.

TLC: $R_f$ = 0.32 (isohexane/EtOAc = 10/1)

ESI-MS: m/z 523.02 $[M+H]^+$

$^1$H NMR: (400 MHz, DMSO-d6) δ 9.08 (s, 1H), 8.90 (s, 1H), 7.72 (dd, *J* = 2.1, 0.7 Hz, 1H), 7.55 - 7.45 (m, 3H), 7.39 (d, *J* = 12.5 Hz, 3H), 7.30 (q, *J* = 7.1, 6.5 Hz, 2H), 3.83 (s, 2H), 1.25 - 1.20 (m, 1H).

**Methyl-5-((3,5-Bis(trifluormethyl)phenyl)amino)-2-((3-(trifluormethyl)phenyl)amino)benzoate (LW-YW22)**

A solution of methyl ester **10** (66.1 mg; 125 µmol) and KOH (85%, 20.2 mg, 0.360 mmol) in ethanol/water (3/1, 3 mL) was heated for 1h at reflux temperature. After being cooled to room temperature the mixture was treated with 2N HCl to adjust a pH value of 1-2. The resulting precipitate was collected by filtration, washed with water and dried in vacuo to yield **LW-YW22** (34.3 mg, 67 µmol, 54%) as a green-yellowish solid.

TLC: $R_f$ = 0.10 (isohexane/EtOAc/HCOOH = 10/1/0.06)

HR-MS: calculated ($C_{22}H_{13}F_9N_20_2$): m/z 507,0833 $[M-H]^-$

found: m/z 507,0761 $[M-H]^-$

$^1$H NMR: (400 MHz, DMSO-d6) δ 13.33 (s, 1H), 9.46 (s, 1H), 8.88 (s, 1H), 7.75 (dd, *J* = 2.3, 0.9 Hz, 1H), 7.63 - 7.43 (m, 3H), 7.41 - 7.34 (m, 3H), 7.30 (dt, *J* = 5.1, 1.6 Hz, 2H), 1.20 (d, *J* = 25.7 Hz, 1H).

$^{13}$C NMR: (101 MHz, $CDCl_3$) δ 171.8, 146.7, 145.1, 141.0, 132.5, 131.1, 130.5, 130.1, 127, 127.1, 125.2, 125.2, 124.7, 122.2, 120.3, 118.7, 115.8, 115.8, 113.8, 113.7, 112.3, 112.2.

Purity: λ = 254: 99%, $t_R$ = 22.1 min (eluent system 1)

λ = 254: 99%, $t_R$ = 20.8 min (eluent system 2)

**<u>NMR spectra of antagonists:</u>**

**LW118**, $^{1}H$ NMR, DMSO-$d_6$, 400 MHz

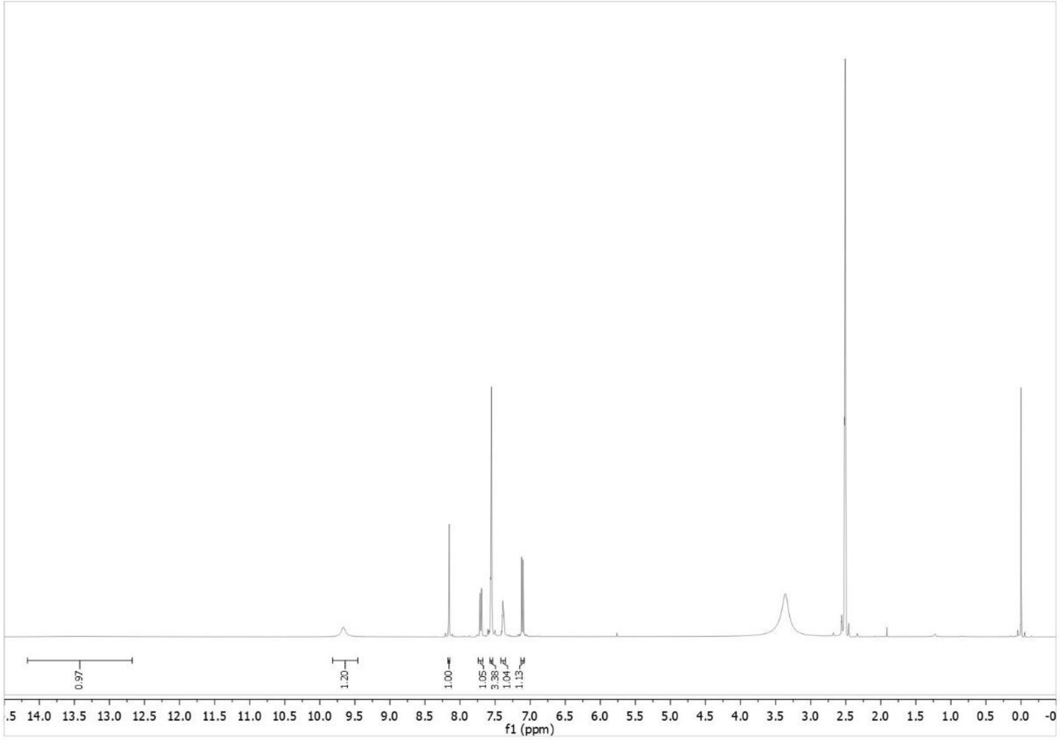


**LW118**, $^{13}C$ NMR, DMSO-$d_6$, 101 MHz

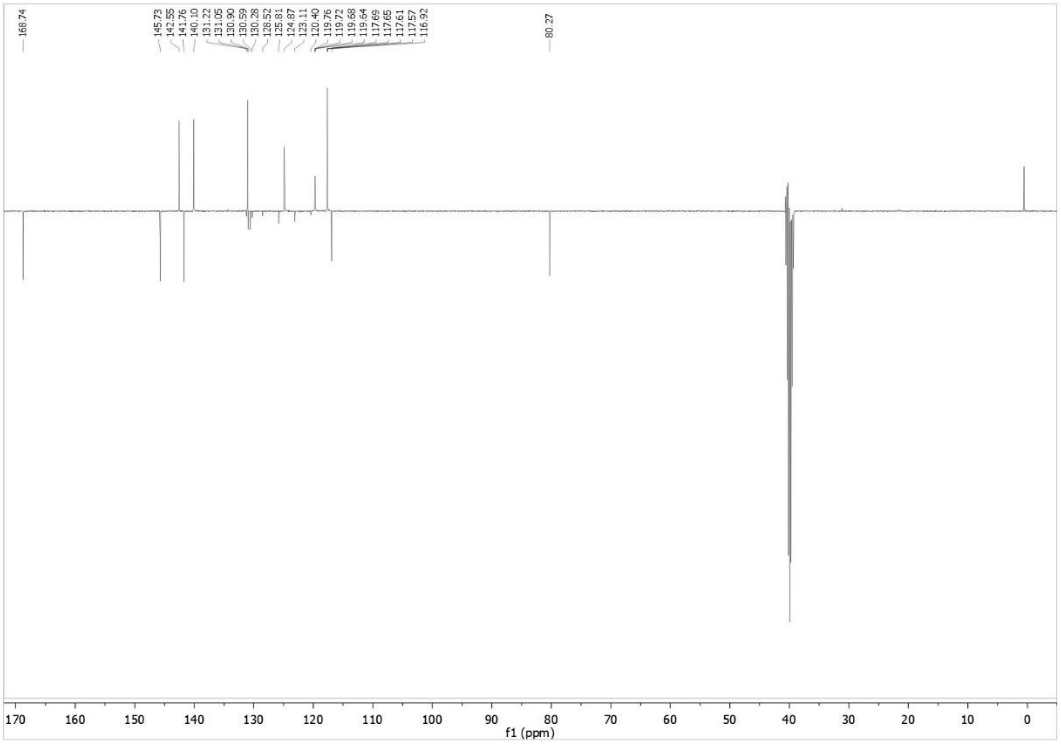

**LW129**, $^{1}$H NMR, $CD_3CN$, 600 MHz

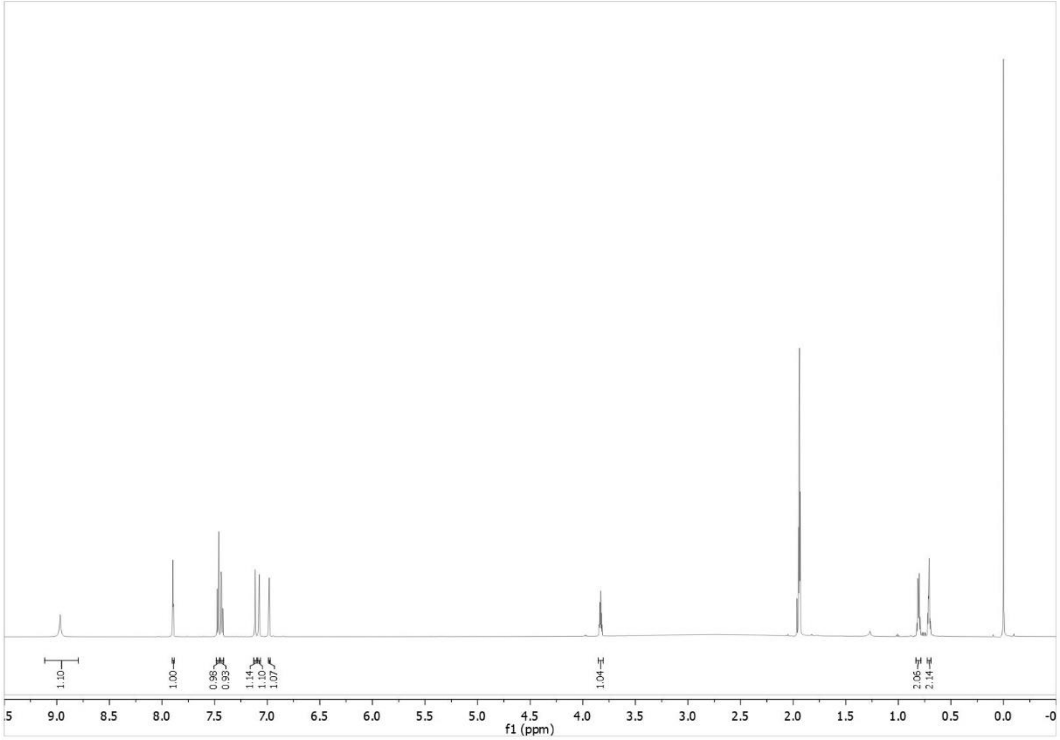

**LW129**, $^{13}$C NMR, $CD_3CN$, 101 MHz

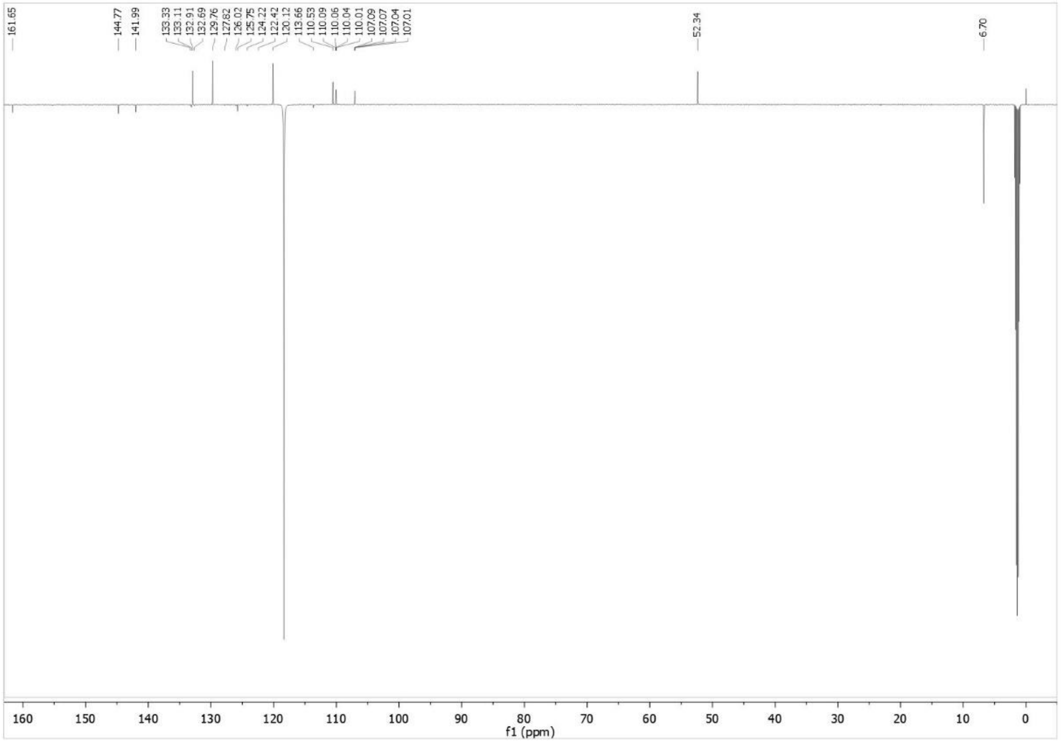

**LW131**, $^1$H NMR, $CD_3CN$, 400 MHz

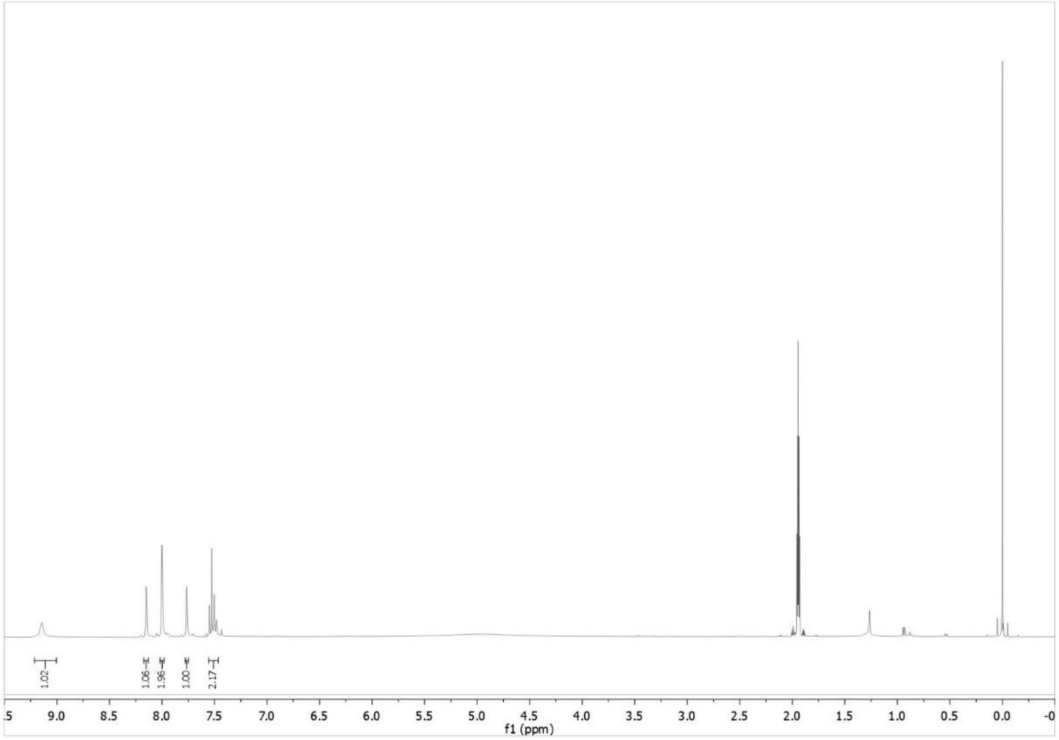

**LW131**, $^{13}$C NMR, $CD_3CN$, 151 MHz

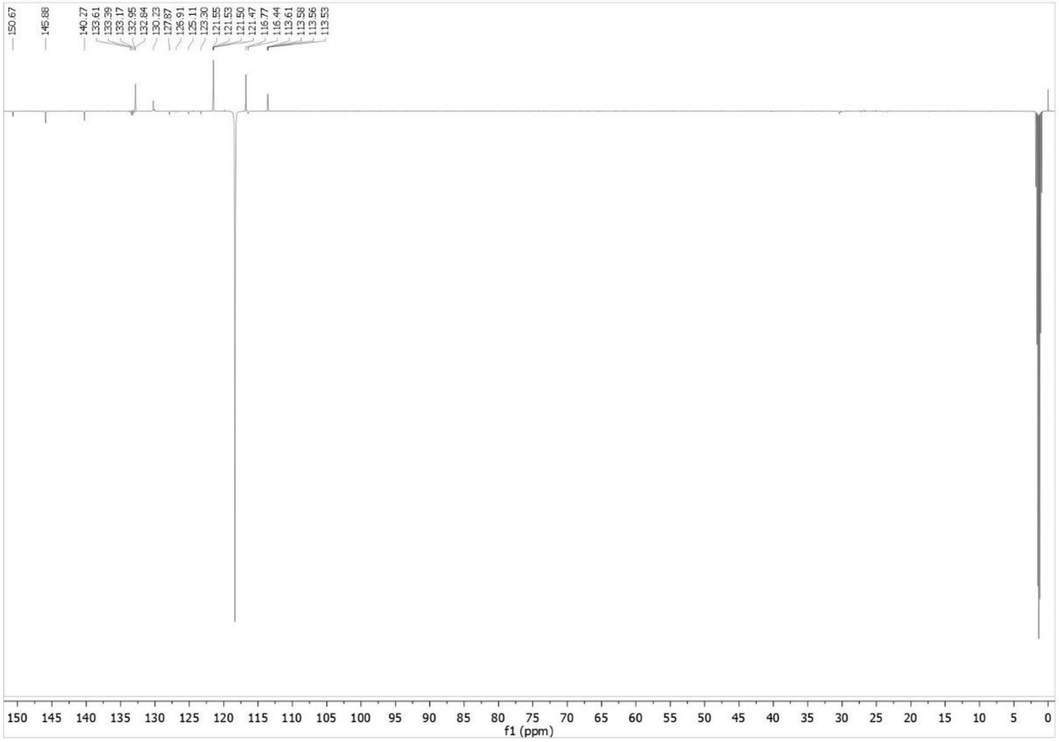

**LW145**, $^{1}$H NMR, DMSO-$d_6$, 400 MHz

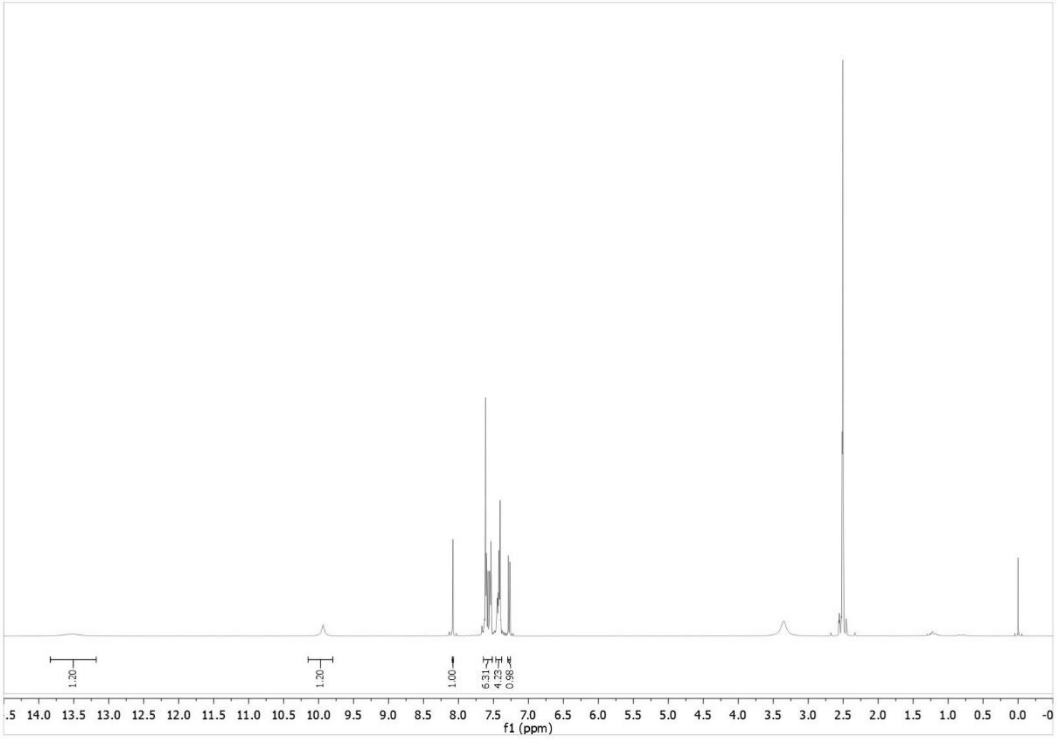

**LW145**, $^{13}$C NMR, DMSO-$d_6$, 151 MHz

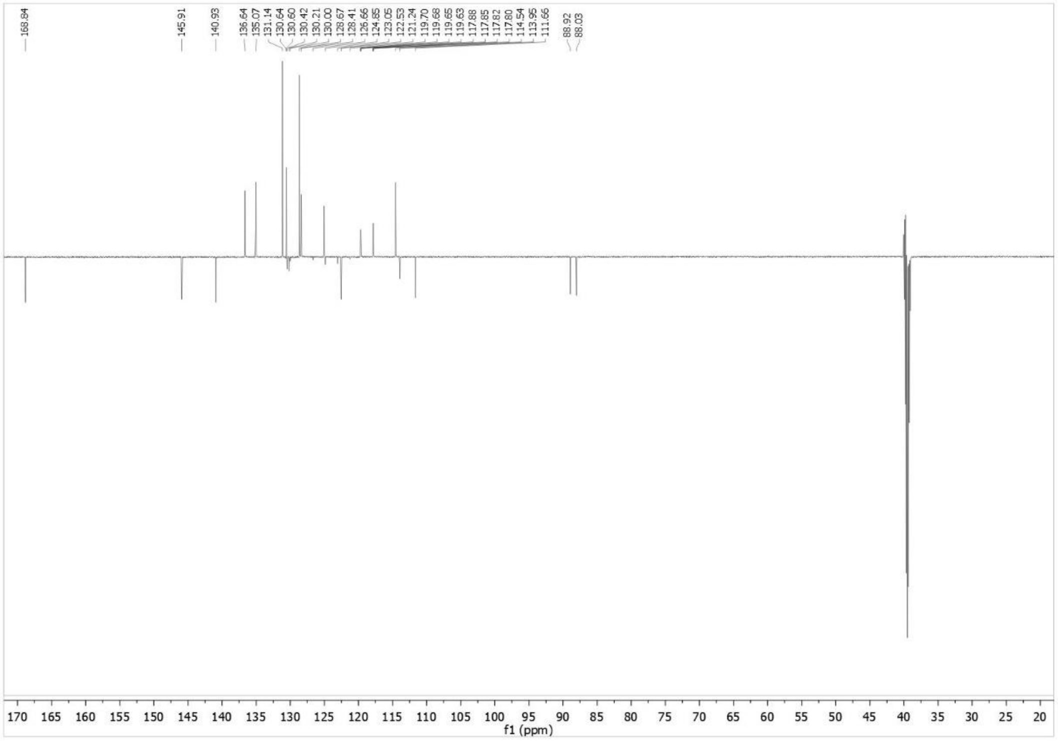

**LW148**, $^{1}$H NMR, DMSO-$d_6$, 400 MHz

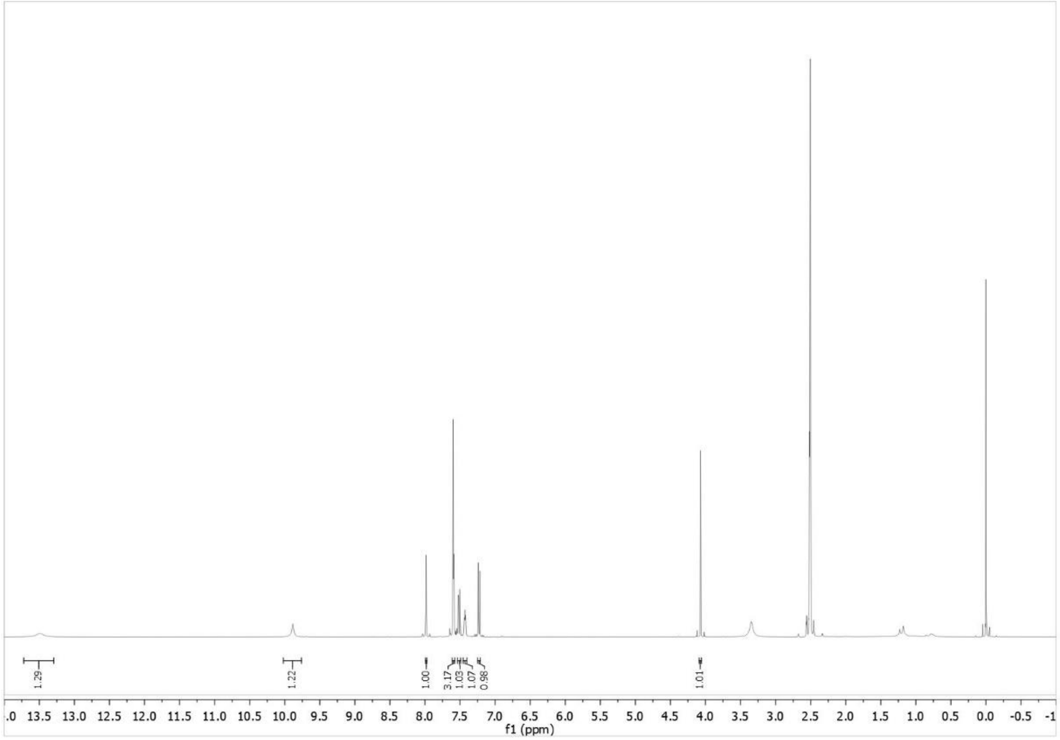

**LW148**, $^{13}$C NMR, DMSO-$d_6$, 101 MHz

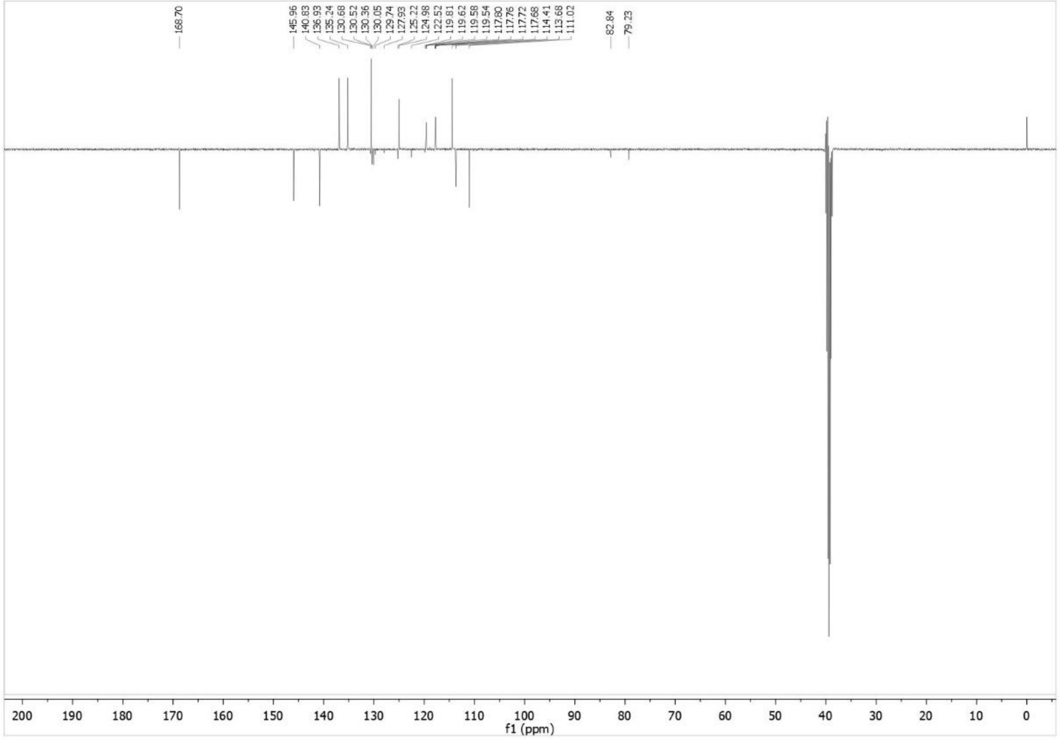

**LW209**, $^1$H NMR, DMSO-$d_6$, 400 MHz

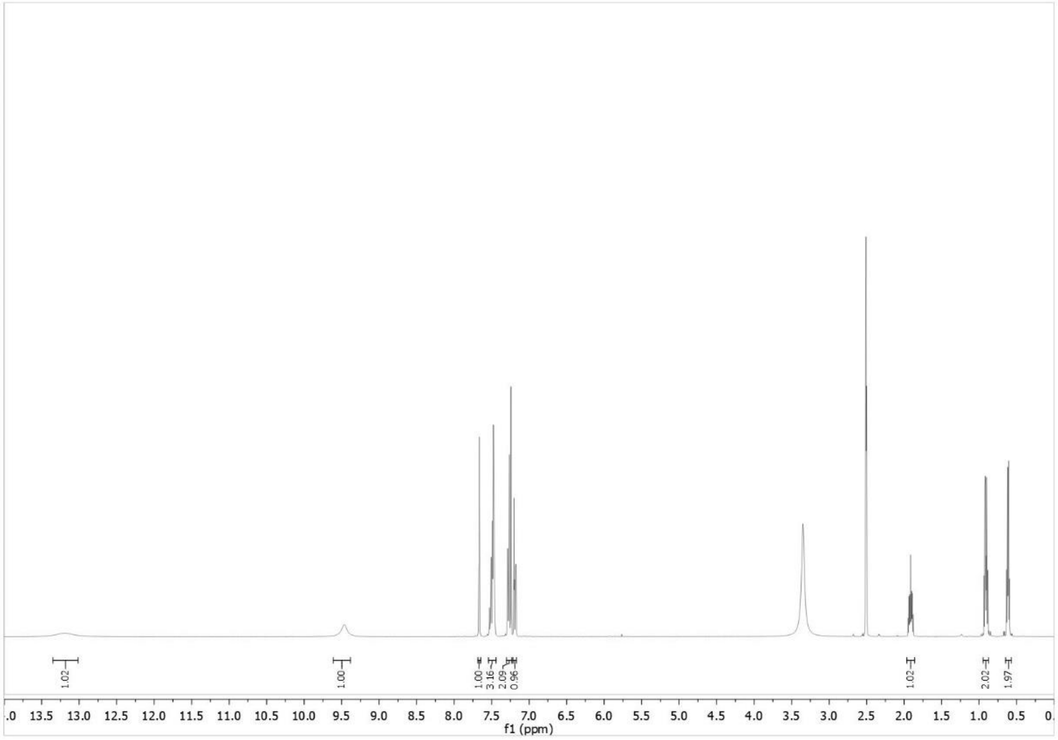

**LW209**, $^{13}$C NMR, DMSO-$d_6$, 101 MHz

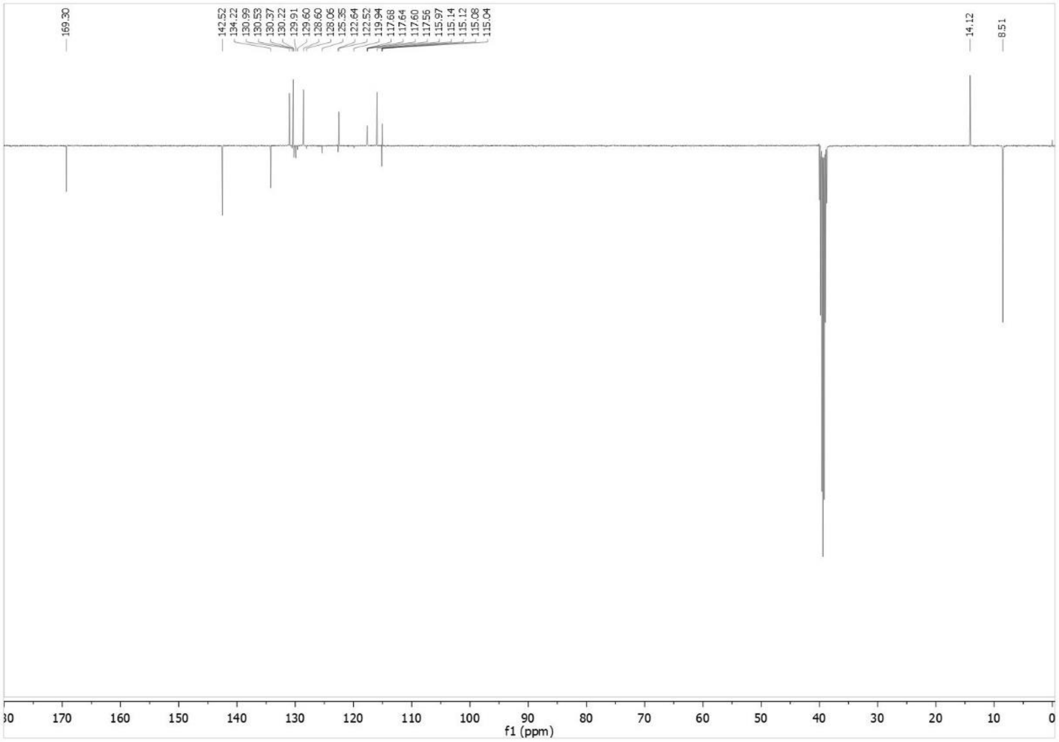

**<u>HPLC runs</u>** (system A: MeOH, system B: ACN)

**LW118**, system A, λ = 254 nm

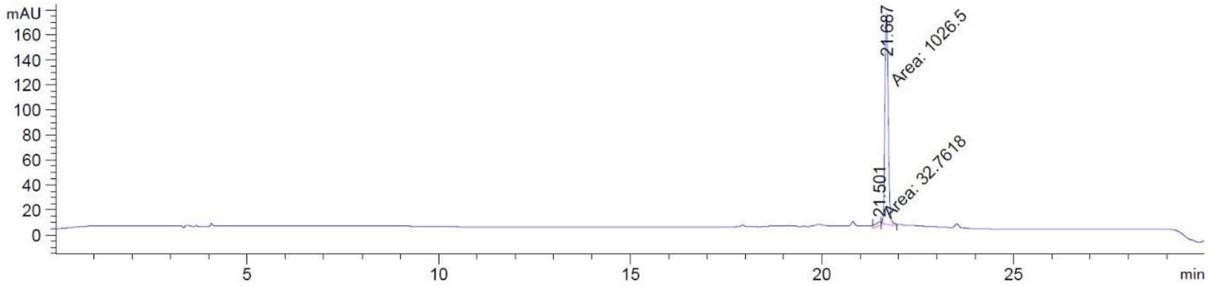


**LW118**, system B, λ = 254 nm

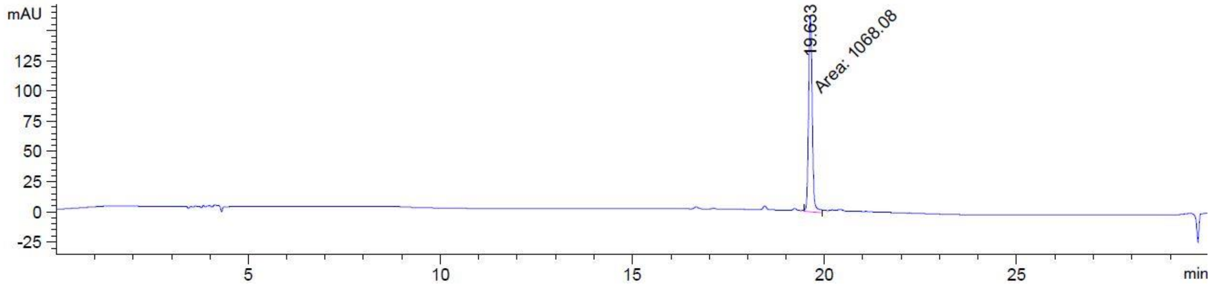


**LW129**, system A, λ = 254 nm

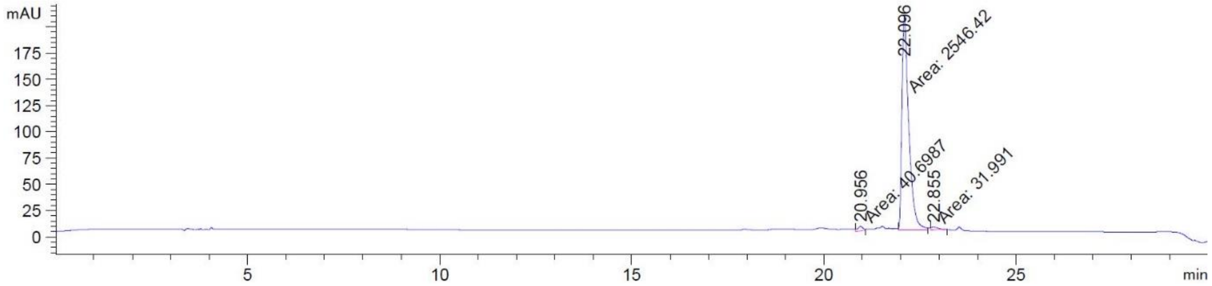


**LW129**, system B, λ = 254 nm

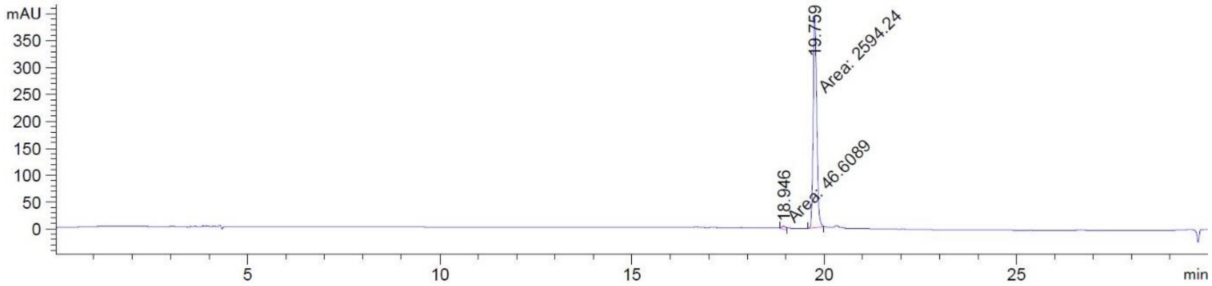

**LW131**, system A, λ = 254 nm

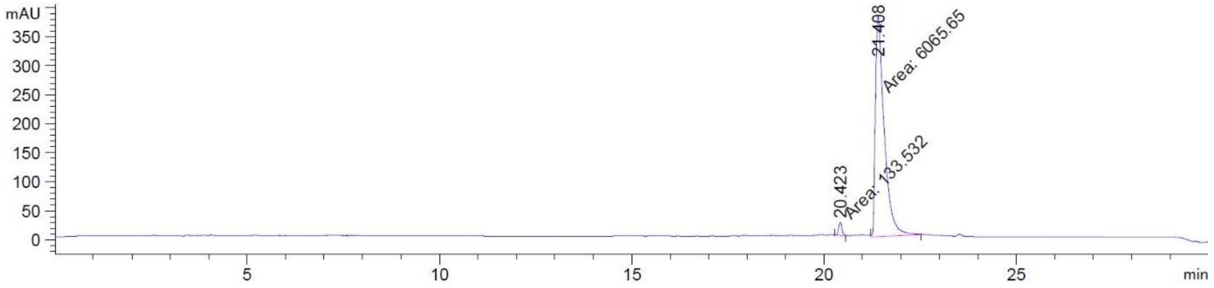


**LW131**, system B, λ = 254 nm

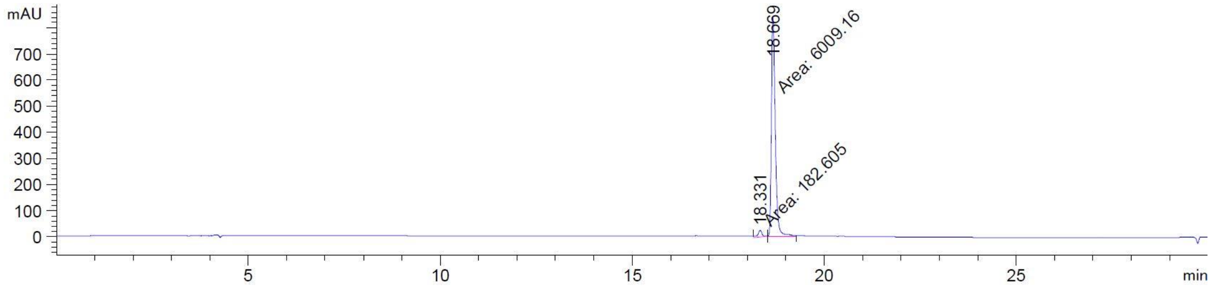


**LW145**, system A, λ = 254 nm

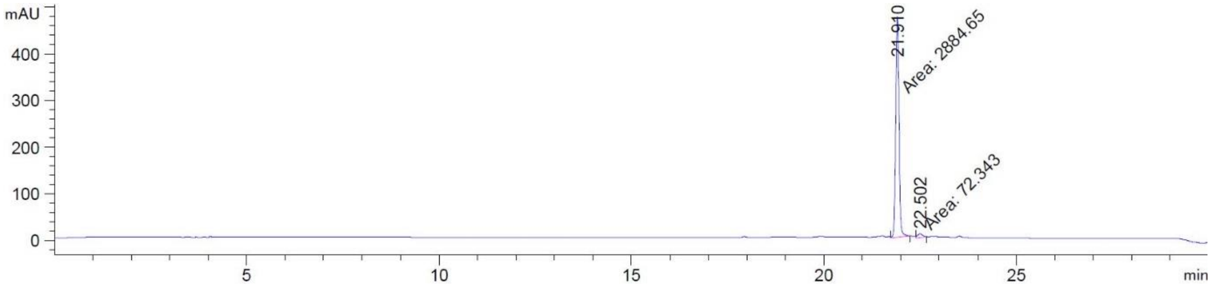


**LW145**, system B, λ = 254 nm

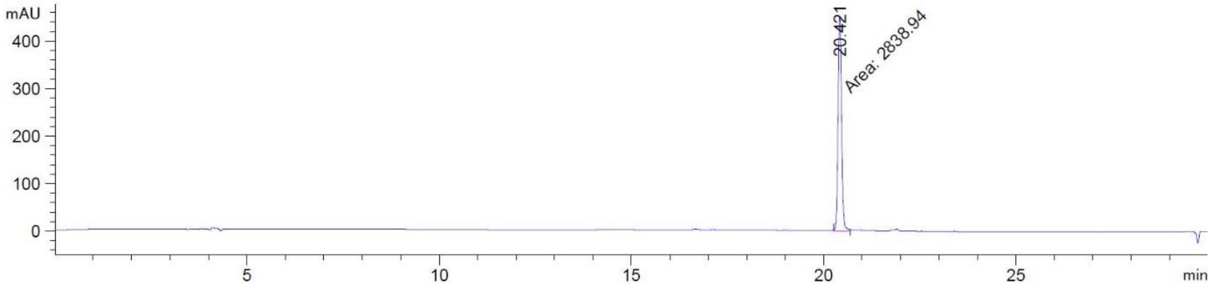

**LW148**, system A, λ = 254 nm

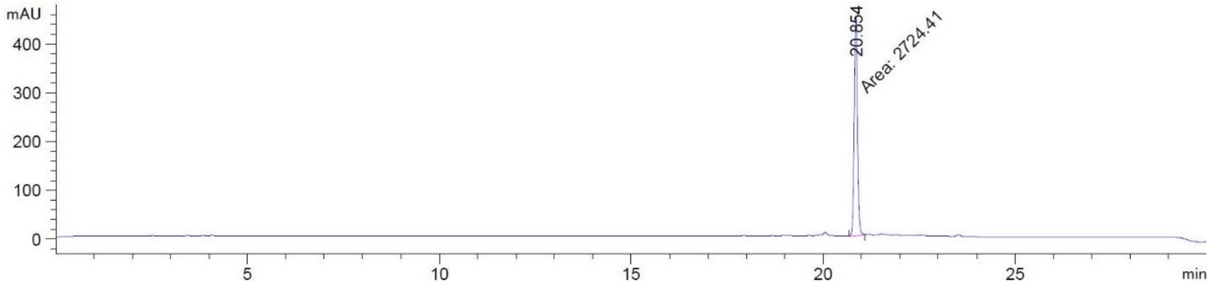


**LW148**, system A, λ = 254 nm

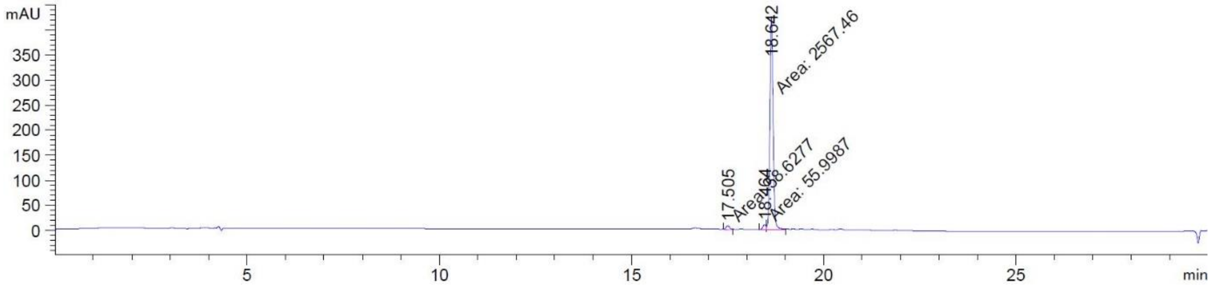


**LW209**, system A, λ = 254 nm

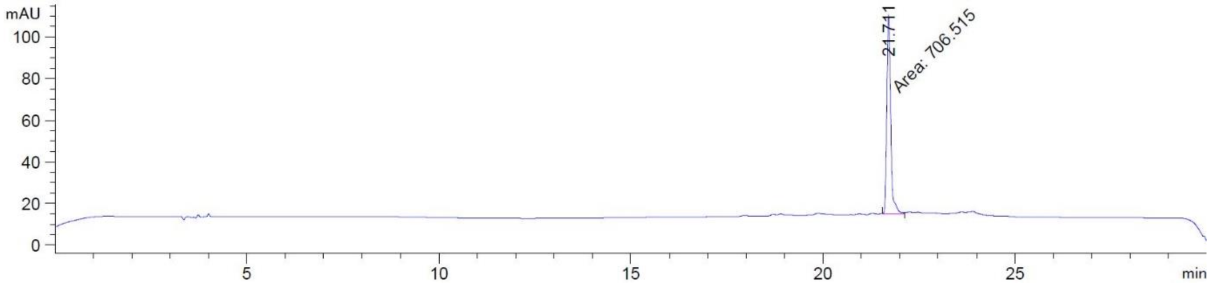


**LW209**, system B, λ = 254 nm

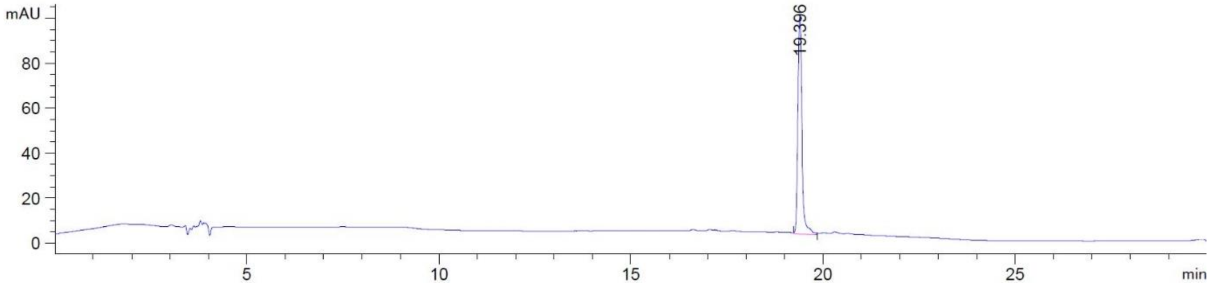

**Analytical data of additional library compounds:**

**2-((3-(Benzyloxy)phenyl)amino)benzenesulfonamide (LW019a)**

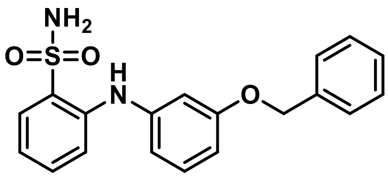

Mp: 104 °C

HR-MS (ESI): m/z $[M+Na]^+$ found: 377.0927, calcd. 377.0930 for $C_{19}H_{18}N_2O_3S$.

$^1$H-NMR (600 MHz, DMSO-d$_6$) δ 7.79 (dd, *J* = 8.0, 1.5 Hz, 1H), 7.68 (s, 1H), 7.56 (s, 2H), 7.46 – 7.43 (m, *J* = 7.1 Hz, 2H), 7.42 – 7.36 (m, 3H), 7.34 (tt, *J* = 7.3 Hz, 1H), 7.29 – 7.26 (m, 1H), 7.24 (t, *J* = 8.1 Hz, 1H), 6.96 – 6.92 (m, 1H), 6.82 (t, *J* = 2.2 Hz, 1H), 6.77 (dd, *J* = 7.9, 1.5 Hz, 1H,), 6.71 (dd, *J* = 8.1, 2.1 Hz, 1H), 5.11 (s, 2H).

$^{13}$C-NMR (151 MHz, DMSO-d$_6$) δ 159.2, 142.3, 140.9, 137.1, 133.1, 130.1, 128.8, 128.5, 128.4, 127.8, 127.5, 118.9, 116.2, 112.8, 109.1, 106.7, 69.1.

**Methyl 2-((3-(trifluoromethyl)phenyl)amino)benzoate (LW050)**[2]

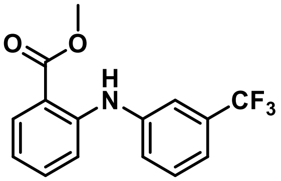

ESI-MS: m/z 296.05 $[M+H]^+$

$^1$H NMR: (400 MHz, $CDCl_3$) δ 9.60 (s, 1H), 8.00 (ddd, *J* = 8.0, 1.7, 0.5 Hz, 1H), 7.50 – 7.47 (m, 1H), 7.46 – 7.35 (m, 3H), 7.32 – 7.27 (m, 2H), 6.82 (ddd, *J* = 8.1, 7.0, 1.2 Hz, 1H), 3.91 (s, 3H).

$^{13}$CNMR: (101 MHz, $CDCl_3$) δ 168.8, 146.7, 141.6, 134.3, 132.4 – 131.1 (m), 129.9, 124.7 – 124.4 (m), 123.9 (q, *J* = 272.4 Hz), 119.5 (q, *J* = 3.8 Hz), 118.3, 118.0 (q, *J* = 3.9 Hz), 114.3, 112.9, 52.0.

HR-MS (ESI): m/z $[M+H]^+$ found 296.0894, calcd. 296.0893 for $C_{15}H_{13}F_3NO_2$.

**2-((3-(Trifluoromethoxy)phenyl)amino)benzoic acid (LW057)**[3]

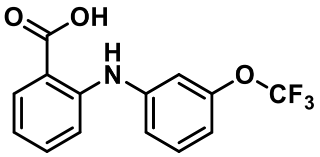

**ESI-MS:** m/z 297.97 $[M+H]^+$

**$^1$H NMR:** (400 MHz, DMSO-$d_6$) δ 13.23 (s, 1H), 9.67 (s, 1H), 7.96 – 7.91 (m, 1H), 7.49 – 7.41 (m, 2H), 7.33 (ddd, *J* = 8.4, 1.2, 0.4 Hz, 1H), 7.27 (ddd, *J* = 8.2, 2.2, 0.9 Hz, 1H), 7.22 – 7.19 (m, 1H), 7.02 – 6.96 (m, 1H), 6.89 (ddd, *J* = 8.1, 7.1, 1.1 Hz, 1H).

**$^{13}$C NMR:** (101 MHz, DMSO-$d_6$) δ 169.6, 149.2, 145.4, 143.0, 134.2, 131.9, 131.0, 124.1 – 116.1 (m), 115.0, 114.4, 114.2, 112.3.

**HR-MS (ESI):** m/z $[M+H]^+$ found 298.0686, calcd. 298.0686 for $C_{14}H_{11}F_3NO_3$.

**Purity:** λ = 254: 99.3%, $t_R$ = 21.0 min (eluent system 1)

λ = 254: 98.7%, $t_R$ = 18.7 min (eluent system 2).

**2-(1*H*-Tetrazol-5-yl)-*N*-(3-(trifluoromethoxy)phenyl)aniline (LW070)**

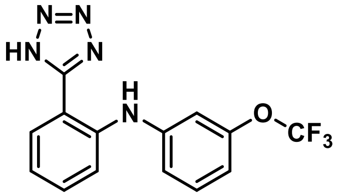

ESI-MS: m/z 322.07 $[M+H]^+$

$^1$H NMR: (400 MHz, DMSO-$d_6$) δ 9.06 (s, 1H), 7.89 (ddd, *J* = 7.8, 1.4, 0.6 Hz, 1H), 7.53 – 7.45 (m, 2H), 7.39 (dd, *J* = 8.2 Hz, 1H), 7.19 – 7.11 (m, 2H), 7.10 – 7.08 (m, 1H), 6.93 – 6.87 (m, 1H).

$^{13}$C NMR: (101 MHz, DMSO-$d_6$) δ 154.1, 149.3, 144.2, 141.2, 132.1, 131.0, 129.8, 121.2, 120.1 (q, *J* = 256.1 Hz), 117.9, 117.1, 113.1, 112.4, 110.5.

HR-MS (ESI): m/z $[M+H]^+$ found 322.0905, calcd. 322.0910 for $C_{14}H_{11}F_3N_5O$.

**N-(3-(Pentafluoro-λ⁶-sulfaneyl)phenyl)-2-(1H-tetrazol-5-yl)aniline (LW71)**

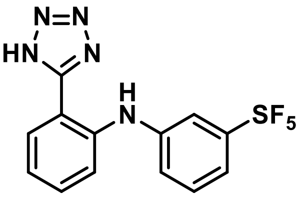


ESI-MS: m/z 364.05 $[M+H]^+$

$^1$H NMR: (400 MHz, $CD_3CN$) δ 9.02 (s, 1H), 7.87 (ddd, *J* = 7.9, 1.4, 0.7 Hz, 1H), 7.64 – 7.61 (m, 1H), 7.51 – 7.40 (m, 5H), 7.13 (ddd, *J* = 7.9, 6.2, 2.3 Hz, 1H).

$^{13}$C NMR: (101 MHz, $CD_3CN$) δ 155.3, 143.8, 142.8, 133.2, 130.9, 130.3, 123.8, 122.0, 120.1 – 119.9 (m), 117.8 – 117.5 (m), 112.2.

HR-MS (ESI): m/z $[M+H]^+$ found 364.0645, calcd. 364.0650 for $C_{13}H_{11}F_{5\,N5}S$.

**N-(3-Chlorophenyl)-2-(1H-tetrazol-5-yl)aniline (LW072)**[5]

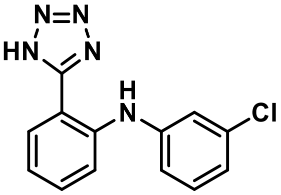


ESI-MS: m/z 271.98 $[M+H]^+$

$^1$H NMR: (400 MHz, DMSO-$d_6$) δ 9.04 (s, 1H), 7.92 – 7.86 (m, 1H), 7.53 – 7.43 (m, 2H), 7.30 (dd, *J* = 8.0 Hz, 1H), 7.19 (dd, *J* = 2.1 Hz, 1H), 7.15 – 7.09 (m, 2H), 6.99 (ddd, *J* = 8.0, 2.0, 0.9 Hz, 1H).

$^{13}$C NMR: (151 MHz, DMSO-$d_6$) δ 154.2, 143.7, 141.4, 133.7, 132.1, 130.9, 129.6, 121.1, 120.8, 118.1, 117.7, 117.2, 111.9.

HR-MS (ESI): m/z $[M+H]^+$ found 272.0694, calcd. 272.0697 for $C_{13}H_{11}ClN_5$.

**2-((4-Chloro-3-(trifluoromethyl)phenyl)amino)benzoic acid (LW076)**[6]

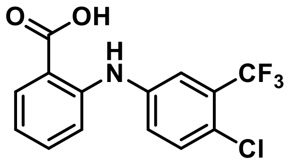


ESI-MS: m/z 316.02 $[M+H]^+$

$^{1}$H NMR: (400 MHz, DMSO-$d_6$) δ 13.23 (s, 1H), 9.64 (s, 1H), 7.94 (dd, *J* = 7.9, 1.7 Hz, 1H), 7.65 (d, *J* = 2.7 Hz, 1H), 7.61 (d, *J* = 8.7 Hz, 1H), 7.53 (dd, *J* = 8.7, 2.7 Hz, 1H), 7.48 (ddd, *J* = 8.7, 7.2, 1.7 Hz, 1H), 7.33 (dd, *J* = 8.4, 1.1 Hz, 1H), 6.94 (ddd, *J* = 8.1, 7.1, 1.1 Hz, 1H).

$^{13}$C NMR: (151 MHz, DMSO-$d_6$) δ 169.3, 144.7, 141.0, 134.1, 132.5, 131.9, 127.4 (q, *J* = 30.7 Hz), 124.2, 122.7 (q, *J* = 273.2 Hz), 122.3 – 122.1 (m), 119.6, 118.7 (q, *J* = 5.4 Hz), 115.7, 115.4.

HR-MS (ESI): m/z [M+H]$^+$ found 316.0348, calcd. 316.0347 for $C_{14}H_{10}ClF_3NO_2$.

**2-((3-Methoxy-5-(trifluoromethyl)phenyl)amino)benzoic acid (LW077)**

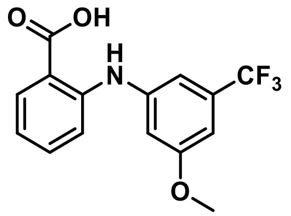


ESI-MS: m/z 312.07 [M+H]$^+$

$^{1}$H NMR: (400 MHz, DMSO-$d_6$) δ 13.22 (s, 1H), 9.65 (s, 1H), 7.94 (dd, *J* = 8.0, 1.7 Hz, 1H), 7.48 (ddd, *J* = 8.7, 7.1, 1.7 Hz, 1H), 7.36 (dd, *J* = 8.5, 1.1 Hz, 1H), 7.12 – 7.10 (m, 1H), 7.08 – 7.06 (m, 1H), 6.94 – 6.88 (m, 1H), 6.87 – 6.85 (m, 1H), 3.82 (s, 3H).

$^{13}$C NMR: (151 MHz, DMSO-$d_6$) δ 169.5, 160.7, 145.1, 143.5, 134.1, 131.9, 131.1 (q, *J* = 31.8 Hz), 123.9 (q, *J* = 272.6 Hz), 119.1, 115.6, 114.9, 108.7, 108.4 (q, *J* = 4.0 Hz), 104.3 (q, *J* = 3.8 Hz), 55.7.

HR-MS (ESI): m/z [M+H]$^+$ found 312.0838, calcd. 312.0842 for $C_{15}H_{13}F_3NO_3$.

***N*-(3-(Difluoromethoxy)phenyl)-2-(1*H*-tetrazol-5-yl)aniline (LW083)**

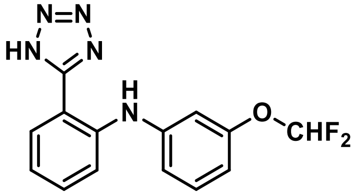


ESI-MS: m/z 304.08 [M+H]$^+$

$^{1}$H NMR: (400 MHz, DMSO-$d_6$) δ 9.11 (s, 1H), 7.92 – 7.87 (m, 1H), 7.49 – 7.47 (m, 2H), 7.34 (dd, *J* = 8.2 Hz, 1H), 7.24 (t, *J* = 74.2 Hz, 1H), 7.14 – 7.02 (m, 2H), 6.97 (dd, *J* = 2.2 Hz, 1H), 6.77 (dd, *J* = 8.1, 2.3 Hz, 1H).

$^{13}$C NMR: (101 MHz, DMSO-$d_6$) δ 154.7, 152.4 (t, *J* = 3.2 Hz), 144.0, 142.0, 132.5, 131.2, 130.0, 121.0, 117.6, 116.8 (t, *J* = 257.1 Hz), 116.0, 111.9, 111.7, 109.3.

HR-MS (ESI): m/z $[M+H]^+$ found 304.1006, calcd. 304.1004 for $C_{14}H_{12}F_{2}N_{5}O$.

**_N_-(2-(1_H_-Tetrazol-5-yl)phenyl)-4-chloro-3-(trifluoromethyl)aniline (LW084)**

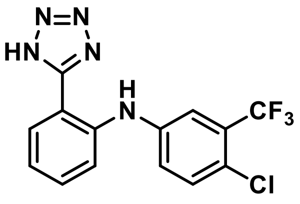


ESI-MS: m/z 340.05 $[M+H]^+$

$^1$H NMR: (400 MHz, DMSO-$d_6$) δ 9.02 (s, 1H), 7.89 (dd, *J* = 7.9, 1.5 Hz, 1H), 7.56 – 7.46 (m, 5H), 7.36 (dd, *J* = 8.8, 2.8 Hz, 1H), 7.23 – 7.16 (m, 1H).

$^{13}$C NMR: (101 MHz, DMSO-$d_6$) δ 154.4, 142.8, 141.0, 132.8, 132.6, 130.5, 127.7 (q, *J* = 30.5 Hz), 123.2 (q, *J* = 273.1 Hz), 122.5, 122.5, 121.2, 119.5, 117.0 (q, *J* = 5.4 Hz), 114.2.

HR-MS (ESI): m/z $[M+H]^+$ found 340.0575, calcd. 340.0571 for $C_{14}H_{10}ClF_3N_5$.

**_N_-(2-(1_H_-Tetrazol-5-yl)phenyl)-3-methoxy-5-(trifluoromethyl)aniline (LW085)**

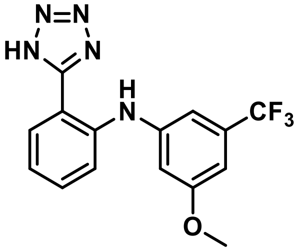


ESI-MS: m/z 336.08 $[M+H]^+$

$^1$H NMR: (600 MHz, $CD_3CN$) δ 8.99 (s, 1H), 7.86 (ddd, *J* = 7.9, 1.6, 0.5 Hz, 1H), 7.51 (ddd, *J* = 8.4, 1.4, 0.5 Hz, 1H), 7.47 (ddd, *J* = 8.5, 7.1, 1.5 Hz, 1H), 7.12 (ddd, *J* = 7.9, 7.1, 1.3 Hz, 1H), 7.06 – 7.05 (m, 1H), 6.97 – 6.96 (m, 1H), 6.85 – 6.83 (m, 1H), 3.82 (s, 3H).

$^{13}$C NMR: (151 MHz, $CD_3CN$) δ 162.2, 155.6, 145.4, 143.0, 133.4 – 132.6 (m), 130.4, 125.2 (q, *J* = 271.7 Hz), 122.0, 118.8, 112.3, 109.3 – 109.1 (m), 105.5 (q, *J* = 3.9 Hz), 56.5.

HR-MS (ESI): m/z $[M+H]^+$ found 336.1066, calcd. 336.1067 for $C_{15}H_{13}F_3N_5O$.

**N-(2-(1H-Tetrazol-5-yl)phenyl)-2-methyl-3-(trifluoromethyl)aniline (LW086)**

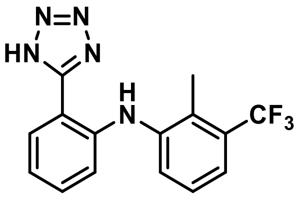

ESI-MS: m/z 320.97 $[M+H]^+$

$^1$H NMR: (600 MHz, $CD_3CN$) δ 9.09 (s, 1H), 7.82 (dd, *J* = 8.2, 1.6 Hz, 1H), 7.58 – 7.56 (m, 1H), 7.50 – 7.48 (m, 1H), 7.39 – 7.33 (m, 2H), 7.01 – 6.98 (m, 2H), 2.42 (q, *J* = 1.6 Hz, 3H).

$^{13}$C NMR: (151 MHz, $CD_3CN$) δ 155.9, 145.1, 142.5, 133.3, 131.8 – 131.7 (m), 130.0, 128.0, 127.8, 122.6 (q, *J* = 5.9 Hz), 120.1, 116.3, 109.6, 14.3 (q, *J* = 2.5 Hz).

HR-MS (ESI): m/z $[M+H]^+$ found 320.1120, calcd. 320.1118 for $C_{15}H_{13}F_3N_5$.

**2-((4-Morpholino-3-(trifluoromethyl)phenyl)amino)benzoic acid (LW101)**

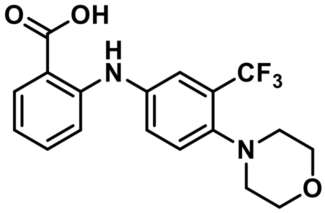

ESI-MS: m/z 367.09 $[M+H]^+$

$^1$H NMR: (400 MHz, DMSO-*d*$_6$) δ 13.16 (s, 1H), 9.63 (s, 1H), 7.92 (dd, *J* = 8.0, 1.7 Hz, 1H), 7.59 – 7.52 (m, 2H), 7.49 (d, *J* = 2.3 Hz, 1H), 7.43 (ddd, *J* = 8.6, 7.1, 1.7 Hz, 1H), 7.22 (dd, *J* = 8.5, 1.0 Hz, 1H), 6.85 (ddd, *J* = 8.1, 7.1, 1.1 Hz, 1H), 3.74 – 3.66 (m, 4H), 2.87 – 2.79 (m, 4H).

$^{13}$C NMR: (101 MHz, DMSO-*d*$_6$) δ 170.1, 147.1 – 146.7 (m), 146.6, 138.7, 134.7, 132.4, 127.5 (q, *J* = 28.1 Hz), 126.6, 125.9, 124.2 (q, *J* = 273.5 Hz), 119.4 (q, *J* = 5.5 Hz), 118.8, 114.8, 114.1, 67.1, 54.0.

HR-MS (ESI): m/z $[M+H]^+$ found 367.1267, calcd. 367.1264 for $C_{18}H_{18}F_3N_2O_3$.

**2-((3-Cyclopropoxy-5-(trifluoromethyl)phenyl)amino)benzoic acid (LW103)**

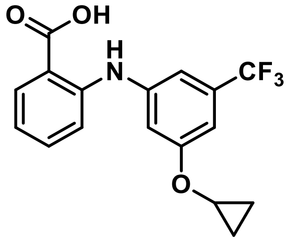

ESI-MS: m/z 338.03 $[M+H]^+$

$^1$H NMR: (400 MHz, DMSO-$d_6$) δ 13.23 (s, 1H), 9.66 (s, 1H), 7.93 (dd, *J* = 7.9, 1.7 Hz, 1H), 7.48 (ddd, *J* = 8.7, 7.1, 1.7 Hz, 1H), 7.39 – 7.34 (m, 1H), 7.20 – 7.17 (m, 1H), 7.15 – 7.11 (m, 1H), 6.96 – 6.89 (m, 2H), 3.95 (dddd, *J* = 5.9, 2.9 Hz, 1H), 0.83 – 0.66 (m, 4H).

$^{13}$C NMR: (101 MHz, DMSO-$d_6$) δ 169.5, 160.1, 145.0, 143.5, 134.1, 132.0, 131.0 (q, *J* = 31.7 Hz), 123.9 (q, *J* = 272.5 Hz) 119.3, 115.7, 115.0, 109.3, 108.8 (q, *J* = 3.9 Hz), 105.1 (q, *J* = 3.8 Hz) 51.3, 5.9.

HR-MS (ESI): m/z $[M+H]^+$ found 338.1000, calcd. 338.0999 for $C_{17}H_{15}F_3NO_3$.

**2-((3-Oxo-6-(trifluoromethyl)-3,4-dihydro-2H-benzo[b][1,4]oxazin-8-yl)amino)benzoic acid (LW104)**

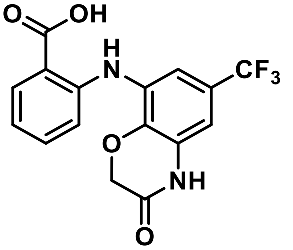


ESI-MS: m/z 353.00 $[M+H]^+$

$^1$H NMR: (400 MHz, DMSO-$d_6$) δ 13.22 (s, 1H), 10.98 (s, 1H), 9.72 (s, 1H), 7.94 (dd, *J* = 8.0, 1.7 Hz, 1H), 7.48 (ddd, *J* = 8.7, 7.2, 1.7 Hz, 1H), 7.32 (dd, *J* = 2.2, 0.8 Hz, 1H), 7.23 (dd, *J* = 8.5, 1.0 Hz, 1H), 6.92 – 6.87 (m, 2H), 4.76 (s, 2H).

$^{13}$C NMR: (101 MHz, DMSO-$d_6$) δ 169.7, 164.4, 145.1, 137.1, 134.2, 131.9, 130.2, 128.6, 124.0 (q, *J* = 271.6 Hz), 118.8, 122.7 (q, *J* = 32.3 Hz), 114.8, 114.1, 110.1 (q, *J* = 3.6 Hz), 106.1 (q, *J* = 4.2, 3.8, 3.8 Hz), 66.9.

HR-MS (ESI): m/z $[M+H]^+$ found 353.0747, calcd. 353.0744 for $C_{16}H_{12}F_3N_2O_4$.

**2-((3-(Benzyloxy)-5-(trifluoromethyl)phenyl)amino)benzoic acid (LW105)**

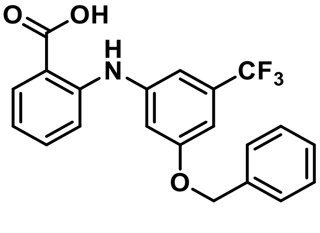


ESI-MS: m/z 387.97 $[M+H]^+$

$^1$H NMR: (600 MHz, $CDCl_3$) δ 9.43 (s, 1H), 8.09 (dd, *J* = 8.1, 1.6 Hz, 1H), 7.47 – 7.36 (m, 7H), 7.25 (dd, *J* = 8.6, 1.1 Hz, 1H), 7.14 – 7.10 (m, 1H), 7.07 – 7.04 (m, 1H), 6.99 – 6.96 (m, 1H), 6.87 (ddd, *J* = 8.1, 7.1, 1.1 Hz, 1H), 5.12 (s, 2H).

$^{13}$C NMR: (151 MHz, $CDCl_3$) δ 172.4, 159.9, 147.3, 142.5, 136.1, 135.3, 132.8 (q, *J* = 32.5 Hz), 132.7, 128.7, 128.3, 127.5, 123.8 (q, *J* = 272.7 Hz), 118.5, 114.6, 111.5, , 111.2 (q, *J* = 3.9 Hz), 111.1, 106.8 (q, *J* = 3.9 Hz), 70.4.

HR-MS (ESI): m/z $[M+H]^+$ found 388.1157, calcd. 388.1155 for $C_{21}H_{17}F_3NO_3$.

**_N_-(2-(1_H_-Tetrazol-5-yl)phenyl)-2-methyl-5-(trifluoromethyl)aniline (LW113)**

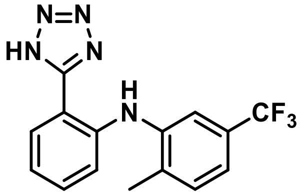


ESI-MS: m/z 319.94 $[M+H]^+$

$^{1}$H NMR: (400 MHz, DMSO-$d_6$) δ 9.18 (s, 1H), 7.97 – 7.92 (m, 1H), 7.55 – 7.43 (m, 3H), 7.36 – 7.29 (m, 1H), 7.20 – 7.15 (m, 1H), 7.11 – 7.04 (m, 1H), 2.38 (s, 3H).

$^{13}$C NMR: (101 MHz, DMSO-$d_6$) δ 154.5, 142.3, 140.5, 134.7 – 134.3 (m), 132.2, 132.0, 129.3, 127.6 (q, *J* = 31.7 Hz), 119.9, 119.2 (q, *J* = 3.9 Hz), 116.0, 115.7 (q, *J* = 3.9 Hz), 110.0, 17.9.

HR-MS (ESI): m/z $[M+H]^+$ found 320.1119, calcd. 320.1118 for $C_{15}H_{13}F_3N_5$.

**_N_-(2-(1_H_-Tetrazol-5-yl)phenyl)-4-morpholino-3-(trifluoromethyl)aniline (LW114)**

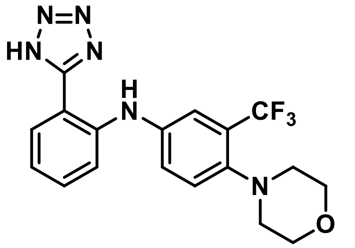


ESI-MS: m/z 390.98 $[M+H]^+$

$^{1}$H NMR: (400 MHz, $CD_3CN$) δ 9.06 (s, 1H), 7.86 – 7.81 (m, 1H), 7.50 – 7.37 (m, 5H), 7.09 – 7.03 (m, 1H), 3.76 – 3.72 (m, 4H), 2.88 – 2.84 (m, 4H).

$^{13}$C NMR: (101 MHz, $CD_3CN$) δ 155.5, 147.8 – 147.6 (m), 143.8, 140.1, 133.3, 130.2, 128.9 (q, *J* = 28.5 Hz), 126.9, 125.9 – 125.8 (m), 125.0 (q, *J* = 272.7 Hz), 121.0, 119.7 (q, *J* = 5.6 Hz), 117.2, 110.9, 68.0, 54.7.

HR-MS (ESI): m/z $[M+H]^+$ found 391.1491, calcd. 391.1489 for $C_{18}H_{18}F_3N_6O$.

**2-((3-Nitro-5-(trifluoromethyl)phenyl)amino)benzoic acid (LW119)**

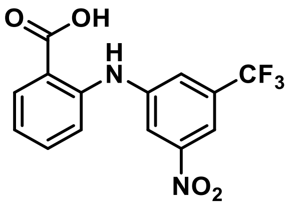

ESI-MS: m/z 326.93 $[M+H]^+$

$^1$H NMR: (600 MHz, DMSO-$d_6$) δ 13.25 (s, 1H), 9.75 (s, 1H), 8.21 (dd, *J* = 2.1 Hz, 1H), 7.97 (dd, *J* = 7.9, 1.7 Hz, 1H), 7.93 – 7.91 (m, 2H), 7.56 (ddd, *J* = 8.7, 7.1, 1.7 Hz, 1H), 7.49 (dd, *J* = 8.3, 1.1 Hz, 1H), 7.09 (ddd, *J* = 8.1, 7.2, 1.2 Hz, 1H).

$^{13}$C NMR: (151 MHz, DMSO-$d_6$) δ 168.7, 149.1, 144.8, 142.7, 133.8, 131.8, 131.1 (q, *J* = 33.1 Hz), 122.9 (q, *J* = 273.0 Hz), 121.4, 120.0 (q, *J* = 3.7 Hz), 118.1, 117.8, 114.8, 111.1 (q, *J* = 4.0 Hz).

HR-MS (ESI): m/z $[M+H]^+$ found 327.0590, calcd. 327.0587 for $C_{14}H_{10}F_3N_2O_4$.

**2-((3-(4-Methyl-1*H*-imidazol-1-yl)-5-(trifluoromethyl)phenyl)amino)benzoic acid (LW122)**

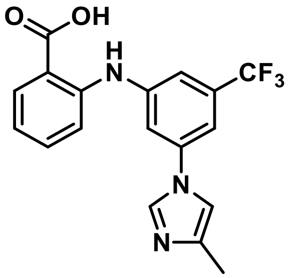

**ESI-MS:** m/z 361.94 $[M+H]^+$

**$^1$H NMR:** (400 MHz, DMSO-$d_6$) δ 9.93 (s, 1H), 8.29 (d, *J* = 1.4 Hz, 1H), 7.96 (dd, *J* = 7.9, 1.7 Hz, 1H), 7.76 – 7.72 (m, 1H), 7.60 – 7.57 (m, 1H), 7.57 – 7.54 (m, 1H), 7.50 (ddd, *J* = 8.7, 7.0, 1.7 Hz, 1H), 7.47 – 7.46 (m, 1H), 7.43 (dd, *J* = 8.4, 1.2 Hz, 1H), 6.96 (ddd, *J* = 8.1, 7.0, 1.2 Hz, 1H), 2.16 (d, *J* = 1.0 Hz, 3H).

**$^{13}$C NMR:** (101 MHz, DMSO-$d_6$) δ 169.5, 144.5, 143.9, 138.7, 138.6, 135.1, 134.0, 131.9, 131.5 (q, *J* = 32.2 Hz), 128.4 – 118.6 (m), 119.7, 116.0, 115.8, 114.3, 113.8, 113.1 (q, *J* = 4.0 Hz), 109.3 (q, *J* = 4.1 Hz), 13.6.

**HR-MS (ESI):** m/z $[M+H]^+$ found 362.1114, calcd. 362.1111 for $C_{18}H_{15}F_3N_3O_2$.

**4-Chloro-*N*-(3-methoxy-5-(trifluoromethyl)phenyl)-2-(1*H*-tetrazol-5-yl)aniline (LW128)**

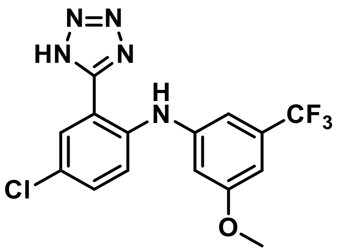


ESI-MS: m/z 369.85 $[M+H]^+$

$^1$H NMR: (400 MHz, $CD_3CN$) δ 9.06 (s, 1H), 7.86 – 7.81 (m, 1H), 7.50 – 7.37 (m, 5H), 7.09 – 7.03 (m, 1H), 3.76 – 3.72 (m, 4H), 2.88 – 2.84 (m, 4H).

$^{13}$C NMR: (101 MHz, $CD_3CN$) δ 155.5, 147.8 – 147.6 (m), 143.8, 140.1, 133.3, 130.2, 128.9 (q, *J* = 28.5 Hz), 126.9, 125.9 – 125.8 (m), 125.0 (q, *J* = 272.7 Hz), 121.0, 119.7 (q, *J* = 5.6 Hz), 117.2, 110.9, 68.0, 54.7.

HR-MS (ESI): m/z $[M+H]^+$ found 370.0679, calcd. 370.0677 for $C_{15}H_{12}ClF_3N_5O$.

***N*-(2-(1*H*-Tetrazol-5-yl)phenyl)-3-nitro-5-(trifluoromethyl)aniline (LW130)**

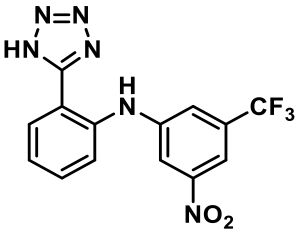


ESI-MS: m/z 351.86 $[M+H]^+$

$^1$H NMR: (400 MHz, $CD_3CN$) δ 9.08 (s, 1H), 8.14 (dd, *J* = 2.2 Hz, 1H), 7.99 – 7.95 (m, 1H), 7.95 – 7.91 (m, 1H), 7.77 – 7.73 (m, 1H), 7.60 – 7.53 (m, 2H), 7.26 (ddd, *J* = 7.9, 6.7, 1.8 Hz, 1H).

$^{13}$C NMR: (101 MHz, $CD_3CN$) δ 155.4, 150.6, 146.3, 141.3, 133.5, 133.2 (q, *J* = 33.6 Hz), 130.8, 124.2 (q, *J* = 272.2 Hz), 124.0, 121.0 (q, *J* = 3.6 Hz), 120.3, 116.2, 114.6, 113.09 (q, *J* = 4.0 Hz).

HR-MS (ESI): m/z $[M+H]^+$ found 351.0813, calcd. 351.0812 for $C_{14}H_{10}F_3N_6O_2$.

**2-((3-(Methoxycarbonyl)-5-(trifluoromethyl)phenyl)amino)benzoic acid (LW153)**

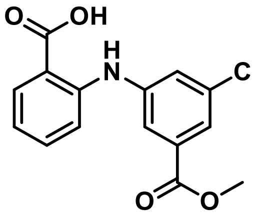


ESI-MS: m/z 339.92 $[M+H]^+$

$^1$H NMR: (400 MHz, DMSO-$d_6$) δ 13.26 (s, 1H), 9.71 (s, 1H), 8.01 (dd, *J* = 1.8 Hz, 1H), 7.96 (dd, *J* = 7.9, 1.7 Hz, 1H), 7.80 (dd, *J* = 2.0 Hz, 1H), 7.74 – 7.71 (m, 1H), 7.52 (ddd, *J* = 8.7, 7.2, 1.7 Hz, 1H), 7.38 (dd, *J* = 8.4, 1.1 Hz, 1H), 7.00 (ddd, *J* = 8.1, 7.2, 1.1 Hz, 1H), 3.88 (s, 3H).

$^{13}$C NMR: (101 MHz, DMSO-$d_6$) δ 169.7, 165.4, 144.6, 143.9, 134.5, 132.4, 132.3, 131.2 (q, *J* = 32.3 Hz), 128.1 – 119.7 (m), 122.7, 120.7, 120.1 (q, *J* = 3.7 Hz), 118.1 (q, *J* = 3.8 Hz), 116.9, 116.8, 53.2.

HR-MS (ESI): m/z $[M+H]^+$ found 340.0795, calcd. 340.0791 for $C_{16}H_{13}F_3NO_4$.

**2-((3-Carbamoyl-5-(trifluoromethyl)phenyl)amino)benzoic acid (LW154)**

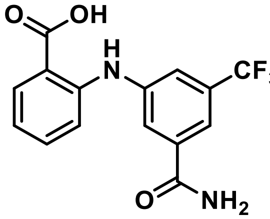


ESI-MS: m/z 346.94 $[M+Na]^+$

$^1$H NMR: (400 MHz, DMSO-$d_6$) δ 13.27 (s, 1H), 9.76 (s, 1H), 8.21 (s, 1H), 8.00 (dd, *J* = 1.8 Hz, 1H), 7.95 (dd, *J* = 7.9, 1.7 Hz, 1H), 7.82 – 7.79 (m, 1H), 7.72 – 7.68 (m, 1H), 7.61 (s, 1H), 7.50 (ddd, *J* = 8.7, 7.2, 1.7 Hz, 1H), 7.35 (dd, *J* = 8.4, 1.1 Hz, 1H), 6.94 (ddd, *J* = 8.1, 7.2, 1.1 Hz, 1H).

$^{13}$C NMR: (101 MHz, DMSO-$d_6$) δ 169.4, 166.1, 144.8, 142.4, 136.4, 134.1, 131.8, 130.23 (q, *J* = 31.9 Hz), 123.73 (q, *J* = 272.7 Hz), 121.9, 119.3, 118.41 (q, *J* = 3.6 Hz), 117.04 (q, *J* = 3.9 Hz), 115.3, 115.0.

HR-MS (ESI): m/z $[M+H]^+$ found 325.0797, calcd. 325.0795 for $C_{15}H_{12}F_3N_2O_3$.

**2-((3-Morpholino-5-(trifluoromethyl)phenyl)amino)benzoic acid (LW167)**

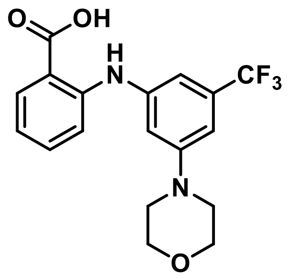

ESI-MS: m/z 366.98 $[M+H]^+$

$^1$H NMR: (400 MHz, DMSO-$d_6$) δ 13.06 (s, 1H), 9.66 (s, 1H), 7.92 (dd, *J* = 7.9, 1.7 Hz, 1H), 7.45 (ddd, *J* = 8.7, 7.1, 1.7 Hz, 1H), 7.29 (dd, *J* = 8.5, 1.0 Hz, 1H), 7.02 (dd, *J* = 2.1 Hz, 1H), 6.95 – 6.92 (m, 1H), 6.89 – 6.83 (m, 2H), 3.75 – 3.70 (m, 4H), 3.21 – 3.17 (m, 4H).

$^{13}$C NMR: (101 MHz, DMSO-$d_6$) δ 169.6, 152.5, 145.7, 142.5, 134.0, 131.8, 130.7 (q, *J* = 31.1 Hz), 124.1 (q, *J* = 272.8 Hz), 118.3, 114.8, 114.0, 109.7, 106.6 (q, *J* = 3.7 Hz), 105.2 (q, *J* = 3.7 Hz), 65.8, 47.6.

HR-MS (ESI): m/z $[M+H]^+$ found 367.1268, calcd. 367.1264 for $C_{18}H_{18}F_3N_2O_3$.

**Methyl 3-((2-(1*H*-tetrazol-5-yl)phenyl)amino)-5-(trifluoromethyl)benzoate (LW169)**

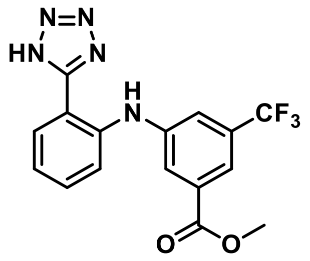

ESI-MS: m/z 364.00 $[M+H]^+$

$^1$H NMR: (400 MHz, $CD_3CN$) δ 9.05 (s, 1H), 7.97 – 7.95 (m, 1H), 7.93 – 7.89 (m, 1H), 7.82 – 7.80 (m, 1H), 7.66 – 7.63 (m, 1H), 7.52 – 7.49 (m, 2H), 7.17 (ddd, *J* = 7.9, 4.9, 3.5 Hz, 1H), 3.89 (s, 3H).

$^{13}$C NMR: (101 MHz, $CD_3CN$) δ 165.4, 154.5, 143.9, 141.3, 132.6, 132.3, 131.5 (q, *J* = 32.7 Hz), 129.5, 128.8 – 119.4 (m), 122.5 – 122.4 (m), 121.8, 119.1 (q, *J* = 3.8 Hz), 118.4 (q, *J* = 3.9 Hz), 118.1, 112.3, 52.2.

HR-MS (ESI): m/z $[M+H]^+$ found 364.1016, calcd. 364.1016 for $C_{16}H_{13}F_3N_5O_2$.

**3-((2-(1*H*-tetrazol-5-yl)phenyl)amino)-5-(trifluoromethyl)benzamide (LW183)**

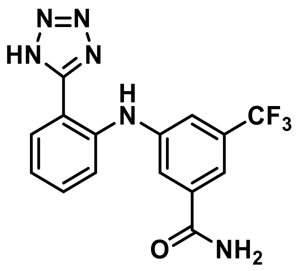

ESI-MS: m/z 349.05 $[M+H]^+$

$^1$H NMR: (400 MHz, DMSO-$d_6$) δ 9.21 (s, 1H), 8.19 (s, 1H), 7.92 (dd, $J$ = 7.8, 1.4 Hz, 1H), 7.88 (dd, $J$ = 1.8 Hz, 1H), 7.72 – 7.70 (m, 1H), 7.57 (s, 1H), 7.55 – 7.47 (m, 3H), 7.19 (ddd, $J$ = 8.1, 6.7, 1.7 Hz, 1H).

$^{13}$C NMR: (101 MHz, DMSO-$d_6$) δ 166.3, 154.2, 143.7, 140.5, 136.3, 131.8, 130.0 (q, $J$ = 31.8 Hz), 129.8, 123.8 (q, $J$ = 272.7 Hz), 121.7, 120.0, 118.5, 116.2 (q, $J$ = 3.5 Hz), 115.7 (q, $J$ = 3.7 Hz), 113.6.

HR-MS (ESI): m/z $[M+H]^+$ found 349.1019, calcd. 349.1019 for $C_{15}H_{12}F_3N_6O$.

***N*-(2-(1*H*-Tetrazol-5-yl)phenyl)-3-cyclopropoxy-5-(trifluoromethyl)aniline (LW190)**

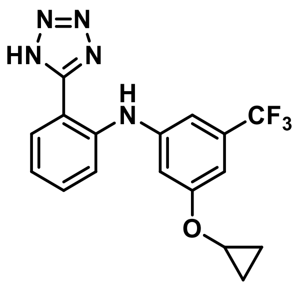

ESI-MS: m/z 362.07 $[M+H]^+$

$^1$H NMR: (400 MHz, $CD_3CN$) δ 9.03 (s, 1H), 7.87 (ddd, $J$ = 7.9, 1.5, 0.5 Hz, 1H), 7.52 (ddd, $J$ = 8.4, 1.4, 0.5 Hz, 1H), 7.47 (ddd, $J$ = 8.4, 7.0, 1.5 Hz, 1H), 7.15 – 7.09 (m, 2H), 7.08 – 7.06 (m, 1H), 6.97 – 6.94 (m, 1H), 3.83 (dddd, $J$ = 6.1, 3.0 Hz, 1H), 0.84 – 0.77 (m, 2H), 0.73 – 0.67 (m, 2H).

$^{13}$C NMR: (101 MHz, $CD_3CN$) δ 161.6, 155.6, 145.3, 142.9, 133.3, 132.90 (q, $J$ = 31.9 Hz), 130.4, 125.2 (q, $J$ = 272.2 Hz), 122.1, 118.8, 112.3, 110.1 – 109.8 (m), 109.5 (q, $J$ = 4.0 Hz), 106.5 (q, $J$ = 4.0 Hz), 52.3, 6.70.

HR-MS (ESI): m/z $[M+H]^+$ found 362.1225, calcd. 362.1223 for $C_{17}H_{15}F_3N_5O$.

**N-(2-(1H-Tetrazol-5-yl)phenyl)-3-methyl-5-(trifluoromethyl)aniline (LW201)**

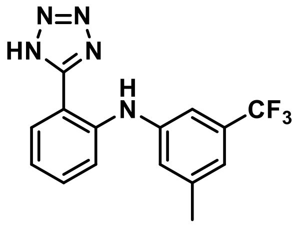

ESI-MS: m/z 320.08 $[M+H]^+$

$^1$H NMR: (400 MHz, $CD_3CN$) δ 9.02 (s, 1H), 7.84 (ddd, *J* = 7.9, 1.4, 0.7 Hz, 1H), 7.49 – 7.43 (m, 2H), 7.30 – 7.26 (m, 2H), 7.16 – 7.13 (m, 1H), 7.09 (ddd, *J* = 7.9, 6.1, 2.3 Hz, 1H), 2.37 (s, 3H).

$^{13}$C NMR: (101 MHz, $CD_3CN$) δ 155.5, 143.7, 143.3, 142.0, 133.4, 132.0 (q, *J* = 31.7 Hz), 130.3, 125.4 (q, *J* = 271.5 Hz), 124.8 – 124.7 (m), 121.6, 120.3 (q, *J* = 3.9 Hz), 118.2, 114.2 (q, *J* = 4.0 Hz), 111.6, 21.5.

HR-MS (ESI): m/z $[M+H]^+$ found 320.1119, calcd. 320.1118 for $C_{15}H_{13}F_3N_5$.

**5-Nitro-2-((3-(trifluoromethyl)phenyl)amino)benzoic acid (LW205)**

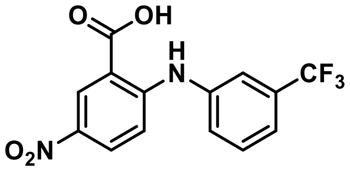

ESI-MS: m/z 327.05 $[M+H]^+$

$^1$H NMR: (400 MHz, DMSO-$d_6$) δ 13.96 (s, 1H), 10.41 (s, 1H), 8.72 (d, *J* = 2.8 Hz, 1H), 8.21 (dd, *J* = 9.4, 2.9 Hz, 1H), 7.76 – 7.56 (m, 4H), 7.19 (d, *J* = 9.4 Hz, 1H).

$^{13}$C NMR: (101 MHz, DMSO-$d_6$) δ 168.4, 151.6, 139.4, 137.3, 130.9, 130.5 (q, *J* = 31.9 Hz), 129.4, 128.3, 127.7 – 127.5 (m), 123.9 (q, *J* = 272.6 Hz), 121.9 (q, *J* = 3.7 Hz), 120.4 (q, *J* = 3.9 Hz), 113.7, 112.0.

HR-MS (ESI): m/z $[M+H]^+$ found 327.0588, calcd. 327.0587 for $C_{14}H_{10}F_3N_2O_4$.

**2-((3-(Trifluoromethyl)phenyl)amino)benzamide (LW208)**

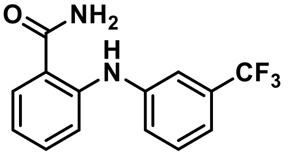

ESI-MS: m/z 281.00 $[M+H]^+$

$^{1}$H NMR: (400 MHz, $CDCl_3$) δ 9.90 (s, 1H), 7.74 – 7.69 (m, 1H), 7.69 – 7.66 (m, 1H), 7.65 – 7.53 (m, 4H), 7.47 – 7.44 (m, 1H), 7.09 – 7.03 (m, 1H), 6.08 (s, 2H).

$^{13}$C NMR: (101 MHz, $CDCl_3$) δ 171.6, 145.2, 142.0, 133.1, 131.7 (q, *J* = 32.2 Hz), 129.8, 128.4, 124.0 (q, *J* = 272.5 Hz), 123.6 – 123.5 (m), 118.8 (q, *J* = 3.9 Hz), 118.7, 116.9 (q, *J* = 3.8 Hz), 115.7.

HR-MS (ESI): m/z $[M+H]^+$ found 281.0895, calcd. 281.0896 for $C_{14}H_{12}F_3N_2O$.

**2-((5-Chloro-2-formylphenyl)amino)benzoic acid (LW231)**

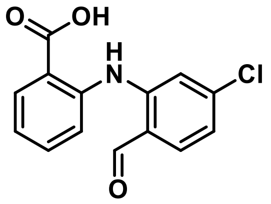


ESI-MS: m/z 275.96 $[M+H]^+$

$^{1}$H NMR: (400 MHz, DMSO-$d_6$) δ 13.29 (s, 1H), 11.20 (s, 1H), 9.94 (d, *J* = 0.7 Hz, 1H), 7.95 (dd, *J* = 7.8, 1.6 Hz, 1H), 7.85 (d, *J* = 8.3 Hz, 1H), 7.62 (dd, *J* = 8.3, 1.3 Hz, 1H), 7.57 (ddd, *J* = 8.4, 7.0, 1.7 Hz, 1H), 7.37 (d, *J* = 1.7 Hz, 1H), 7.14 (ddd, *J* = 8.1, 7.0, 1.4 Hz, 1H), 7.06 (dd, *J* = 8.3, 1.9 Hz, 1H).

$^{13}$C NMR: (101 MHz, DMSO-$d_6$) δ 193.7, 168.4, 145.3, 141.4, 140.7, 139.0, 133.9, 132.2, 123.0, 121.2, 120.8, 120.6, 119.7, 114.5.

HR-MS (ESI): m/z $[M+H]^+$ found 276.0422, calcd. 276.0422 for $C_{14}H_{11}ClNO_3$.

**3-(Methoxycarbonyl)-2-((3-(trifluoromethyl)phenyl)amino)benzoic acid (LW253)**

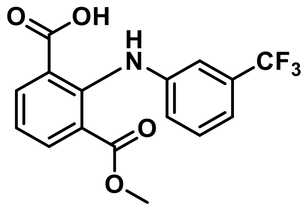


ESI-MS: m/z 339.99 $[M+H]^+$

$^{1}$H NMR: (400 MHz, DMSO-$d_6$) δ 13.38 (s, 1H), 9.61 (s, 1H), 8.06 (dd, *J* = 7.8, 1.7 Hz, 1H), 7.86 (dd, *J* = 7.7, 1.8 Hz, 1H), 7.47 – 7.38 (m, 1H), 7.23 – 7.19 (m, 1H), 7.19 – 7.15 (m, 2H), 7.13 (dd, *J* = 7.8 Hz, 1H), 3.42 (s, 3H).

$^{13}$C NMR: (101 MHz, DMSO-$d_6$) δ 168.6, 166.8, 144.5, 142.5, 135.1, 135.1, 130.1, 129.7 (q, *J* = 31.5 Hz), 124.0 (q, *J* = 272.4 Hz), 121.9, 121.1, 121.0 – 120.8 (m), 120.5, 117.3 (q, *J* = 3.9 Hz), 112.9 (q, *J* = 3.9 Hz), 51.7.

HR-MS (ESI): m/z $[M+H]^+$ found 340.0791, calcd. 340.0791 for $C_{16}H_{13}F_3NO_4$.

**2-((3-(Cyclopropylethynyl)-5-(trifluoromethyl)phenyl)amino)benzoic acid (LW271)**

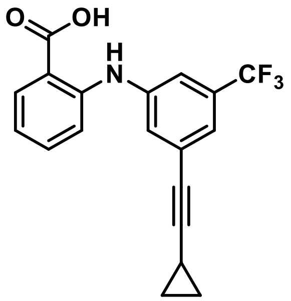


ESI-MS: m/z 346.06 $[M+H]^+$

$^1$H NMR: (400 MHz, $CDCl_3$) δ 9.37 (s, 1H), 8.08 (dd, *J* = 8.2, 1.7 Hz, 1H), 7.49 – 7.39 (m, 2H), 7.39 – 7.36 (m, 1H), 7.36 – 7.31 (m, 1H), 7.29 – 7.23 (m, 1H), 6.86 (ddd, *J* = 8.0, 7.1, 1.1 Hz, 1H), 1.49 – 1.41 (m, 1H), 0.95 – 0.80 (m, 4H).

$^{13}$C NMR: (101 MHz, $CDCl_3$) δ 172.9, 147.2, 141.3, 135.4, 132.8, 132.0 (q, *J* = 32.6 Hz), 127.5, 126.1, 123.5 (q, *J* = 272.7 Hz), 123.2 (q, *J* = 3.7 Hz), 118.6, 117.5 (q, *J* = 3.7 Hz), 114.4, 111.6, 95.6, 74.3, 8.7, 0.1.

HR-MS (ESI): m/z $[M+H]^+$ found 346.1051, calcd. 346.1049 for $C_{19}H_{15}F_3NO_2$.

**2-Butyl-1-(3-(trifluoromethyl)phenyl)-1*H*-indole-7-carboxylic acid (LW273)**

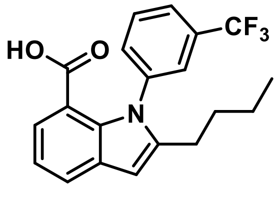


ESI-MS: m/z 262.05 $[M+H]^+$

$^1$H NMR: (600 MHz, $CDCl_3$) δ 7.78 (dd, *J* = 7.8, 1.2 Hz, 1H), 7.61 – 7.56 (m, 2H), 7.52 (dd, *J* = 7.9 Hz, 1H), 7.49 (dd, *J* = 1.9 Hz, 1H), 7.40 – 7.36 (m, 1H), 7.18 (dd, *J* = 7.6 Hz, 1H), 6.51 (t, *J* = 1.0 Hz, 1H), 2.51 – 2.45 (m, 3H), 1.60 – 1.53 (m, 2H), 1.36 – 1.29 (m, 2H), 0.85 (t, *J* = 7.4 Hz, 3H).

$^{13}$C NMR: (151 MHz, $CDCl_3$) δ 171.6, 144.2, 141.0, 135.5, 131.5 (q, *J* = 32.8 Hz), 131.1, 130.8, 129.4, 125.0 – 124.7 (m), 124.4 (q, *J* = 3.7 Hz), 123.6 (q, *J* = 272.5 Hz), 119.7, 114.8, 101.5, 30.4, 27.0, 22.3, 13.8.

HR-MS (ESI): m/z $[M+H]^+$ found 362.1361, calcd. 362.1362 for $C_{20}H_{19}F_3NO_2$.

**5-(2-((3-(Trifluoromethyl)phenyl)amino)phenyl)isoxazol-3-ol (LW306)**

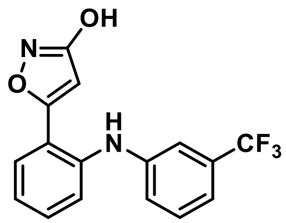

ESI-MS: m/z 321.10 $[M+H]^+$

$^1$H NMR: (400 MHz, DMSO-$d_6$) δ 11.29 (s, 1H), 8.22 (s, 1H), 7.79 (dd, *J* = 7.8, 1.5 Hz, 1H), 7.48 (ddd, *J* = 8.1, 7.2, 1.6 Hz, 1H), 7.42 – 7.36 (m, 2H), 7.27 (ddd, *J* = 7.8, 7.2, 1.3 Hz, 1H), 7.14 – 7.10 (m, 1H), 7.10 – 7.05 (m, 2H), 6.26 (s, 1H).

$^{13}$C NMR: (101 MHz, DMSO-$d_6$) δ 170.5, 166.5, 145.5, 138.8, 131.2, 130.2, 129.9 (q, *J* = 31.1 Hz), 128.3 – 119.9 (m), 128.2, 124.1, 123.8, 121.7, 118.4 – 118.1 (m), 114.9 (q, *J* = 3.7 Hz), 111.0 (q, *J* = 3.9 Hz), 94.4.

HR-MS (ESI): m/z $[M+H]^+$ found 321.0843, calcd. 321.0845 for $C_{16}H_{12}F_3N_2O_2$.

**2-((4-(Hex-1-yn-1-yl)-3-(trifluoromethyl)phenyl)amino)benzoic acid (LW308)**

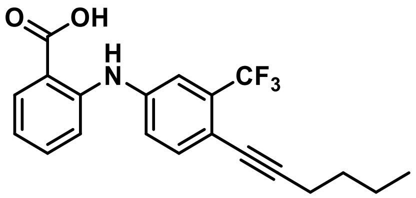

ESI-MS: m/z 362.29 $[M+H]^+$

$^1$H NMR: (600 MHz, DMSO-$d_6$) δ 13.22 (s, 1H), 9.73 (s, 1H), 7.94 (dd, *J* = 7.9, 1.7 Hz, 1H), 7.52 – 7.46 (m, 3H), 7.44 (dd, *J* = 8.5, 2.3 Hz, 1H), 7.36 (dd, *J* = 8.4, 1.0 Hz, 1H), 6.94 (ddd, *J* = 8.0, 7.2, 1.0 Hz, 1H), 2.45 (t, *J* = 6.9 Hz, 2H), 1.56 – 1.49 (m, 2H), 1.48 – 1.41 (m, 2H), 0.91 (t, *J* = 7.3 Hz, 3H).

$^{13}$C NMR: (151 MHz, DMSO-$d_6$) δ 169.2, 144.2, 141.2, 135.1, 133.8, 131.8, 130.9 (q, *J* = 29.5 Hz), 123.3 (q, *J* = 273.5 Hz), 121.6, 119.6, 116.1 (q, *J* = 5.3 Hz), 116.0, 115.8, 113.3 – 113.2 (m), 94.7, 76.5, 30.0, 21.1, 18.4, 13.3.

HR-MS (ESI): m/z $[M+H]^+$ found 362.1360, calcd. 362.1362 for $C_{20}H_{19}F_3NO_2$.

**2-((4-(Prop-1-yn-1-yl)-3-(trifluoromethyl)phenyl)amino)benzoic acid (LW310)**

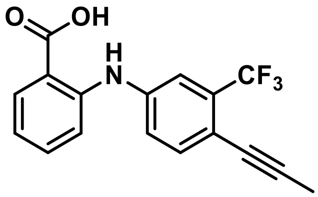

ESI-MS: m/z 320.09 $[M+H]^+$

$^1$H NMR: (400 MHz, DMSO-$d_6$) δ 13.25 (s, 1H), 9.67 (s, 1H), 7.94 (dd, $J$ = 8.0, 1.7 Hz, 1H), 7.55 – 7.43 (m, 4H), 7.37 (dd, $J$ = 8.4, 1.1 Hz, 1H), 6.95 (ddd, $J$ = 8.1, 7.2, 1.1 Hz, 1H), 2.08 (s, 3H).

$^{13}$C NMR: (101 MHz, DMSO-$d_6$) δ 169.2, 144.2, 141.2, 135.2, 134.0, 131.8, 130.8 (q, $J$ = 29.4 Hz), 123.3 (q, $J$ = 273.5 Hz), 121.6, 119.6, 116.2 – 115.9 (m), 115.5, 113.3 (q, $J$ = 2.2 Hz), 90.5, 75.5, 4.0.

HR-MS (ESI): m/z $[M+H]^+$ found 320.0890, calcd. 320.0893 for $C_{17}H_{13}F_3NO_2$.

**2-((4-(Pentylcarbamoyl)-3-(trifluoromethyl)phenyl)amino)benzoic acid (LW317)**

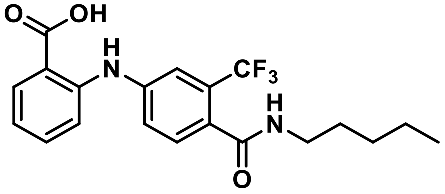

ESI-MS: m/z 365.21 $[M+H]^+$

$^1$H NMR: (400 MHz, DMSO-$d_6$) δ 13.24 (s, 1H), 9.70 (s, 1H), 8.38 (dd, $J$ = 5.7 Hz, 1H), 7.95 (dd, $J$ = 8.0, 1.7 Hz, 1H), 7.55 – 7.47 (m, 3H), 7.46 – 7.39 (m, 1H), 7.33 (dd, $J$ = 8.4, 1.1 Hz, 1H), 6.94 (ddd, $J$ = 8.1, 7.2, 1.1 Hz, 1H), 3.24 – 3.14 (m, 2H), 1.55 – 1.41 (m, 2H), 1.36 – 1.22 (m, 5H), 0.93 – 0.85 (m, 3H).

$^{13}$C NMR: (101 MHz, DMSO-$d_6$) δ 169.3, 166.6, 144.6, 142.3, 134.0, 131.9, 129.9, 129.9 – 129.6 (m), 127.7 – 119.2 (m), 127.2 (q, $J$ = 31.2 Hz), 121.7, 119.4, 116.7 (q, $J$ = 5.1 Hz), 115.4, 115.2, 38.8, 28.4, 21.7, 13.8.

HR-MS (ESI): m/z $[M+H]^+$ found 395.1583, calcd. 395.1577 for $C_{20}H_{22}F_3N_2O_3$.

***N*-(2-(1*H*-tetrazol-5-yl)phenyl)-4-phenyl-6-(trifluoromethyl)pyrimidin-2-amine (LW341)**

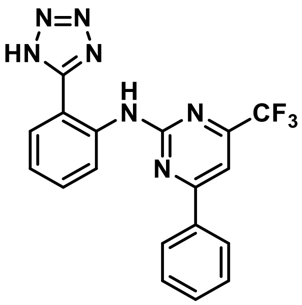

ESI-MS: m/z 384.16 $[M+H]^+$

$^1$H NMR: (600 MHz, DMSO-$d_6$) δ 10.92 (s, 1H), 8.53 (dd, *J* = 8.4, 1.1 Hz, 1H), 8.28 – 8.24 (m, 2H), 7.98 (dd, *J* = 7.9, 1.5 Hz, 1H), 7.92 (s, 1H), 7.67 (ddd, *J* = 8.6, 7.3, 1.6 Hz, 1H), 7.65 – 7.57 (m, 3H), 7.33 (ddd, *J* = 7.6, 1.2 Hz, 1H).

$^{13}$C NMR: (151 MHz, DMSO-$d_6$) δ 167.0, 159.8, 155.8 (q, *J* = 34.9 Hz), 137.6, 135.1, 131.9, 131.6, 129.0, 128.9, 127.4, 123.1, 121.6, 120.6 (q, *J* = 275.6 Hz), 114.0 – 112.9 (m), 104.2 (q, *J* = 2.8 Hz).

HR-MS (ESI): m/z $[M+H]^+$ found 384.1176, calcd. 384.1179 for $C_{18}H_{13}F_3N_7$.

***N*-(2-(1*H*-Tetrazol-5-yl)phenyl)-4-iodo-6-(trifluoromethyl)pyrimidin-2-amine (LW360)**

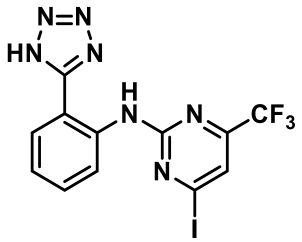

ESI-MS: m/z 434.25 $[M+H]^+$

$^1$H NMR: (600 MHz, $CD_3CN$) δ 11.02 (s, 1H), 8.66 (dd, *J* = 8.5, 1.1 Hz, 1H), 7.95 (dd, *J* = 7.9, 1.5 Hz, 1H), 7.70 (s, 1H), 7.65 (ddd, *J* = 8.7, 7.3, 1.6 Hz, 1H), 7.33 (ddd, *J* = 8.1, 7.4, 1.1 Hz, 1H).

$^{13}$C NMR: (151 MHz, $CD_3CN$) δ 159.7, 155.4 (q, *J* = 36.2 Hz), 138.6, 133.2, 132.4, 129.7, 124.4, 123.7 – 117.9 (m), 121.7, 120.6 (q, *J* = 2.9 Hz), 113.1.

HR-MS (ESI): m/z $[M+H]^+$ found 433.9832, calcd. 433.9833 for $C_{12}H_8F_3IN_7$.

**2-(1*H*-1,2,3-Triazol-5-yl)-*N*-(3-(trifluoromethyl)phenyl)aniline (LW-TP09)**

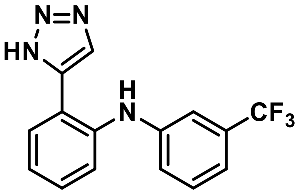

HR-MS: found: 305.1010 $[M+H]^+$

[1]H-NMR: (400 MHz, CDCl3) δ 11.86 (s, 1H), 8.78 (s, 1H), 8.05 (s, 1H), 7.67 (dd, *J* = 7.8 Hz, 1.6 Hz, 1H), 7.50 – 7.27 (m, 5H), 7.23 – 7.16 (m, 1H), 7.06 – 6.95 (m, 1H).

[13]C-NMR: (101 MHz, $CDCl_3$) δ 143.2, 140.7, 130.0, 129.7, 129.0, 122.3, 120.7, 118.2, 117.0, 115.9.

**2-(Isoxazol-4-yl)-*N*-(3-(trifluoromethyl)phenyl)aniline (LW-PH05)**

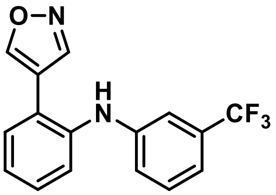

HR-MS: found 327.0716 $[M+Na]^+$

[1]H-NMR: (400 MHz, $CDCl_3$) δ 8.65 (s, 1H), 8.51 (s, 1H), 7.43 -7.39 (m, 1H), 7.37 - 7.31 (m, 3H), 7.19 (ddt, J = 7.6 Hz, 5.0 Hz, 4.0 Hz, 1H), 7.14 (ddt, J = 7.6 Hz, 1.7 Hz, 0.9 Hz, 1H), 7.12 - 7.09 (m, 1H), 7.04 - 7.00 (m, 1H), 5.53 (s, 1H).

[13]C-NMR: (101 MHz, $CDCl_3$) δ б 155.3, 149.3, 144.3, 139.3, 132.1, 131.9, 130.1, 130.1, 129.5, 124.3, 126.74 -121.19 (m), 122.0, 119.6, 117.5, 117.3 (q, *J* = 3.8 Hz), 112.9 (q, *J* = 3.8 Hz).

**2-((2-Chloro-3,5-bis(trifluoromethyl)phenyl) amino) benzoic acid (LW-JK05)**

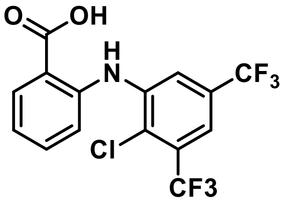

HR-MS: found: 382.0063 $[M-H]^-$

| | |
|---|---|
| $^{1}$H NMR: | (400 MHz, DMSO-$d_6$) δ 13.55 (s, 1H), 10.17 (s, 1H), 8.01 (dd, *J* = 8.0 Hz, 1.5 Hz, 2H), 7.70 (d, *J* = 1.3 Hz, 1H), 7.58 (ddd, *J* = 8.7 Hz, 7.4 Hz, 1.6 Hz, 1H), 7.42 (dd, *J* = 8.3 Hz, 0.7 Hz, 1H), 7.10-7.06 (m, 1H). |
| $^{13}$C-NMR | (101 MHz, DMSO-$d_6$) δ 169.4, 143.1, 141.4, 129.7 -127.8 (m), 124.8, 121.1, 127.2 -117.6 (m), 118.4, 118.2, 116.5, 116.0. |

**2-((2-Chloro-3,5-bis(trifluoromethyl))-2-(1*H*-terazol-5-yl) aniline (LW-JK06)**

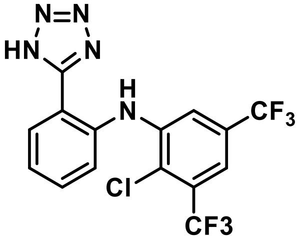


| | |
|---|---|
| HR-MS: | found: 406.0300 [M-H]$^-$ |
| $^{1}$H NMR: | (400 MHz, DMSO-$d_6$) δ 9.60 (s, 1H), 8.01 (dd, *J* = 7.8 Hz, 1.2 Hz, 1H), 7.76 (d, *J* = 1.4 Hz, 1H), 7.64 - 7.55 (m, 3H), 7.32 (ddd, *J* = 8.2 Hz, 7.0 Hz, 1.5 Hz, 1H). |
| $^{13}$C-NMR | (101 MHz, DMSO-$d_6$) δ 142.4, 139.0, 132.1, 129.6, 129.5 - 128.0 (m), 123.5, 123.2, 127.0 -119.0 (m), 119.9, 117.0, 115.1, 114.2. |

**2-((3-(4-Chlorobenzyloxy)phenyl)amino) benzoic acid (LW-MR04)**

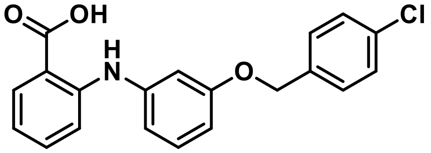


| | |
|---|---|
| HR-MS: | found: 352.0740 [M-H]$^-$ |
| $^{1}$H NMR: | (400 MHz, $CDCl_3$) δ 9.32 (s, 1H), 8.04 (dd, *J* = 8.1 Hz, 1.4 Hz, 1H), 7.37 (s, 3H), 7.33 (dt, *J* = 10.5 Hz, 2.5 Hz, 1H), 7.29 - 7.24 (m, 3H), 7.22 (dd, *J* = 8.5 Hz, 0.7 Hz, 1H), 6.89 - 6.85 (m, 2H), 6.77 (ddd, *J* = 8.1 Hz, 7.1 Hz, 1.1 Hz, 1H), 6.72 (ddd, *J* = 8.3 Hz, 2.3 Hz, 0.9 Hz, 1H), 5.04 (s, 2H). |
| $^{13}$C-NMR | (101 MHz, $CDCl_3$) δ 172.5, 159.4, 148.4, 141.7, 135.3, 135.3, 135.2, 135.2, 133.8, 132.5, 128.8, 128.7, 117.4, 115.5, 114.4, 110.5, 110.4, 110.3, 109.1, 69.2. |

**2-((3-(4-Chlorobenzyloxy)phenyl)amino) benzoic acid (LW-MR05)**

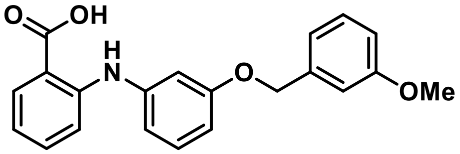


HR-MS: found: 348.1235 $[M-H]^-$

$^1$H NMR: (400 MHz, $CDCl_3$) δ 9.33 (s, 1H), 8.04 (dt, *J* = 8.1 Hz, 1,6 Hz, 1H), 7.37 - 7,30 (m, 2H), 7.29 - 7.25 (m, 2H), 7.25 - 7.21 (m, 1H), 7.03 - 6,98 (m, 2H}, 6.91 - 6.84 (m, 3H), 6.79 - 6.73 (m, 2H), 5.06 (d, *J* = 6.6 Hz, 2H), 3,85 (*d*, *J* = 20,0 Hz, 3H).

$^{13}$C-NMR (101 MHz, $CDCl_3$) δ 173.2, 159.8, 159.6, 148.5, 141.6, 138.4, 135.2, 132.6, 130.2, 129.7, 119.5, 117.4, 115.3, 114.4, 113.6, 112.8, 110.4, 109.2, 69.9, 69.1, 55.3.

**2-((3-(4-Chlorobenzyloxy)phenyl)amino) benzoic acid (LW-MR06)**

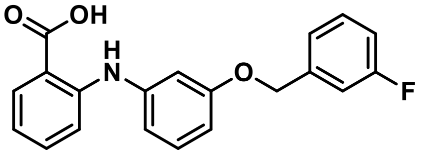


HR-MS: found: 336.1035 $[M-H]^-$

$^1$H NMR: (400 MHz, $CDCl_3$) δ 9.33 (s, 1H), 8.04 (dt, *J* = 8.1 Hz, 1.6 Hz, 1H), 7.37 - 7.30 (m, 2H), 7.29 - 7.25 (m, 2H), 7.25 - 7.21 (m, 1H), 7.03 - 6.98 (m, 2H), 6.91 - 6.84 (m, 3H), 6.79 - 6.73 (m, 2H), 5.06 (d, *J* = 6.6 Hz, 2H), 3.85 (d, *J* = 20.0 Hz, 3H).

$^{13}$C-NMR (101 MHz, $CDCl_3$) δ 173.2, 159.4, 148.4, 141.7, 135.2, 132.6, 130.2, 130.1, 122.7, 117.5, 115.5, 114.9, 114.7, 114.4, 114.3, 114.1, 110.6, 110.3, 109.1, 69.1.

**2-((3-((4-Chlorobenzyl)oxy)-4-methoxyphenyl)amino) benzoic acid (LW-BK19)**

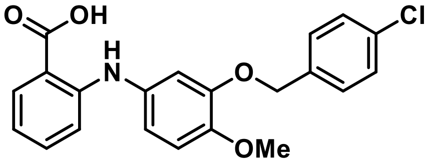


HR-MS: found: 382.0849 $[M-H]^-$

| | |
|---|---|
| $^1$H NMR: | (400 MHz, $CDCl_3$) δ 12.98 (s, 1H), 9.45 (s, 1H), 7.86 (dd, J = 8.0 Hz, 1.6 Hz, 1H), 7.51-7.42 (m, 4H), 7.27 (ddd, *J* = 8.7 Hz, 7.1 Hz, 1.7 Hz, 1H), 6.98 (d, *J* = 8.6 Hz, 1H), 6.91 - 6.85 (m, 2H), 6.79 (dd, *J* = 8.5 Hz, 2.4 Hz, 1H), 6.73-6.66 (m, 1H), 5.10 (s, 2H), 3.78 (s, 3H). |
| $^{13}$C-NMR | (101 MHz, $CDCl_3$) δ 170.5 (s), 148.9, 148.4. 146.5, 136.6, 134.6, 133.6, 132.8, 132.2, 129.9, 128.9, 116.9, 116.2, 113.4, 111.8, 110.6. 69.4, 56.3. |

**2-((3-((3-Fluorobenzyl)oxy)-4-methoxyphenyl)amino) benzoic acid (LW-BK20)**

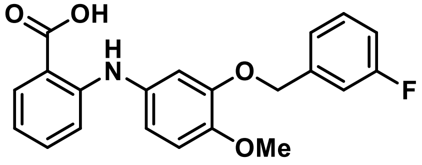


| | |
|---|---|
| HR-MS: | found: 366.1146 [M-H]$^-$ |
| $^1$H NMR: | (400 MHz, DMSO-$d_6$) δ 12.98 (s, 1H), 9.46 (s, 1H), 7.86 (dd, *J* = 8.0 Hz, 1.6 Hz, 1H), 7.48 - 7.42 (m, 1H), 7.29-7.23 (m, 3H), 7.18 (ddd, *J* = 8.2 Hz, 2.7 Hz, 1.3 Hz, 1H), 7.02 - 6.97 (m, 1H), 6.93 (d, *J* = 2.5 Hz, 1H), 6.89-6.85 (m, 1H), 6.80 (dd, *J* = 8.5 Hz, 2.5 Hz, 1H), 6.69 (ddd, *J* = 8.0 Hz, 7.1 Hz, 1.0 Hz, 1H), 5.13 (s, 2H), 3.79 (s, 3H). |
| $^{13}$C-NMR | (101 MHz, DMSO-$d_6$) δ 170.5, 162.6, 148.9, 148.4, 146.5, 140.5, 134.6, 133.6, 132.3, 130.9, 123.9, 116.9, 116.2, 115.0, 114.7, 113.4, 113.4, 111.8, 110.5, 69.4, 56.3. |

**2-((4-Methoxy-3-(3-methoxybenzyl)oxy)phenyl)amino) benzoic acid (LW-BK21)**

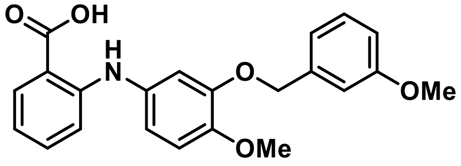


| | |
|---|---|
| HR-MS: | found: 378.1344 [M-H]$^-$ |
| $^1$H NMR: | (400 MHz, DMSO-$d_6$) δ 12.97 (s, 1H), 9.45 (s, 1H), 7.86 (dd, *J* = 8.0 Hz, 1.6 Hz, 1H), 7.34 - 7.24 (m, 2H), 7.02 - 6.95 (m, 3H), 6.93 - 6.85 (m, 3H), 6.78 (dd, *J* = 8.5 Hz, 2.4 Hz, 1H), 6.69 (ddd, *J* = 8.0 Hz, 7.1 Hz, 1.0 Hz, 1H), 5.08 (s, 2H), 3.78 (s, 3H), 3.74 (s, 3H). |
| $^{13}$C-NMR | (101 MHz, DMSO-$d_6$) δ 170.5, 159.7, 148.8, 148.6, 146.5, 139.1, 134.7, 133.6, 132.2, 130.0, 120.2, 116.9, 115.9, 113.6, 113.4, 111.7. 110.5, 70.1, 56.4, 55.5. |

**2-((3-((4-Chlorobenzyl)oxy)-4-methyphenyl)amino) benzoic acid (LW-LB17)**

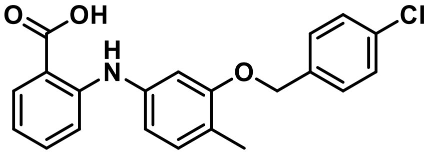


HR-MS: found: 366.0894 [M-H]$^-$

$^1$H NMR: (400 MHz, DMSO-$d_6$) δ 13,05 (s, 1H): 9.58 (s, 1H); 7.88 (dd, *J* = 8.0 Hz, 1.6 Hz, 1H); 7.47 (s, 4H), 7.32 – 7.27 (m, 1H); 7.15 - 7,11 (m. 1H), 7.05 – 6.99 (m, 1H), 6.87-6.85 (m, 1H), 6.76-6.70 (m, 2H), 5.14 (s, 2H), 2.19 (s, 3H).

$^{13}$C-NMR (101 MHz, DMSO-$d_6$) δ 170.0, 156.6, 147.4, 139.1, 136.4, 134.1, 132.2, 131.8, 131.0, 129.0, 128.5, 121.0, 117.0, 113.9, 113.4, 112.0, 106.5, 68.2, 15.7.

**2-((3-((3-Methoxybenzyl)oxy)-4-methyphenyl)amino) benzoic acid (LW-LB18)**

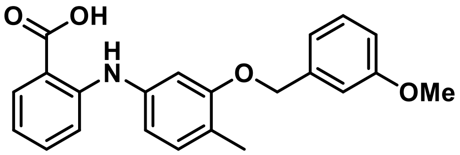


HR-MS: found: 362.1398 [M-H]$^-$

$^1$H NMR: (400 MHz, DMSO-$d_6$) δ 13.04 (s, 1H), 9.58 (s, 1H), 7.88 (dd, *J* = 8.0 Hz, 1.6 Hz, 1H), 7.35 - 7.26 (m, 2H), 7,14 – 7,11 (m, 1H), 7.05 – 7.00 (m, 3H), 6.92 – 6.85 (m, 2H), 6.77 – 6.70 (m, 2H), 5.12 (s, 2H), 3.75 (s, 3H), 2.19 (s, 3H).

$^{13}$C-NMR (101 MHz, DMSO-$d_6$) δ 170.0, 159.3, 156.7, 147.4, 139.1, 139.0, 134.2, 131.8, 130.9, 129.8, 121.0, 119.2, 116.9, 113.7, 113.4, 113.0, 112.7, 111.9, 106.4, 68.8, 55.0, 15.7.

**2-((3-((3-Fluorobenzyl)oxy)-4-methyphenyl)amino) benzoic acid (LW-LB19)**

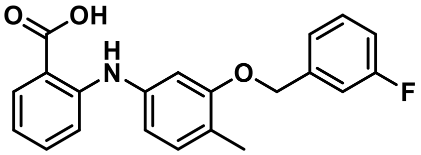


HR-MS: found: 350.1198 [M-H]$^-$

$^1$H NMR: (400 MHz, DMSO-$d_6$) δ 13.04 (s, 1H), 9.58 (s, 1H), 7.88 (dd, *J* = 8.0 Hz, 1.6 Hz, 1H), 7.52 - 7.40 (m, 1H), 7,34 – 7,24 (m, 3H), 7.21 – 7.11 (m, 2H), 7.06 – 6.98 (m, 1H), 6.88 – 6.86 (m, 1H), 5.17 (s, 2H), 2.20 (s, 3H).

$^{13}$C-NMR (101 MHz, DMSO-$d_6$) δ 170.0, 162.2, 156.5, 147.4, 140.4, 139.1, 134.2, 131.8, 131.0, 130.5, 123.0, 121.1, 117.0, 114.4, 113.9, 113.8, 113.5, 112.0, 106.4, 68.2, 15.7.
