## Supplementary File 3 for "Inhibiting a promiscuous GPCR: iterative discovery of bitter taste receptor ligands"

### Supplementary File 3: FDA-approved drug library screening

#### Inhibiting a promiscuous GPCR: iterative discovery of bitter taste receptor ligands

Fabrizio Fierro[1], Lior Peri[1], Harald Hubner[2], Alina Tabor-Schkade[2], Lukas Waterloo[2], Stefan Löber[2], Tara Pfeiffer[2], Dorothee Weikert[2], Tamir Dingjan[1], Eitan Margulis[1], Peter Gmeiner[2], Masha Y Niv[1*]

1. The Institute of Biochemistry, Food Science and Nutrition, Robert H. Smith Faculty of Agriculture, Food and Environment, The Hebrew University of Jerusalem, Rehovot, Israel

2. Department of Chemistry and Pharmacy, Medicinal Chemistry, Friedrich-Alexander-Universität Erlangen-Nürnberg, Nikolaus-Fiebiger-Str. 10, 91058 Erlangen, Germany

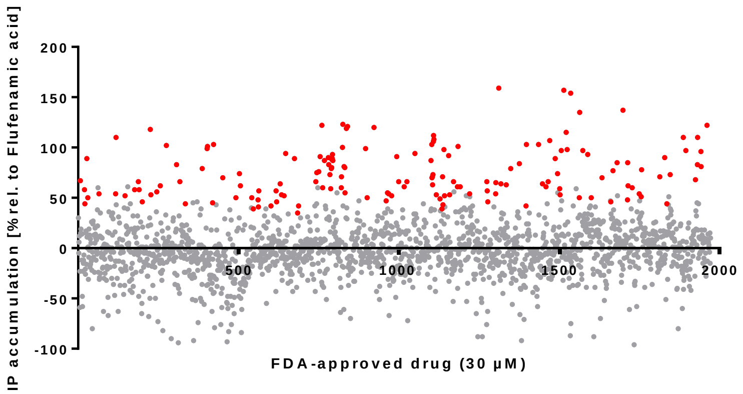


**Supplementary File 3 Fig. S1.** Primary screening of the FDA-approved drug library (DiscoveryProbe™, ApexBio). Dot plot of 1791 FDA-approved drugs screened at a concentration of 30 µM on HEK293T cells expressing TAS2R14 using the IP-One HTRF® assay. Highlighted are the initial active (red) and inactive (grey) compounds relative to 30 µM Flufenamic acid (100 % activation). The threshold for hit identification of an active compound was set as the mean of the unstimulated control plus three times the standard deviation. Data represent mean of duplicate assay plates.

**Supplementary File 3 Table S1.** Results primary screen - list of the initial active library compounds, screened at a concentration of 30 µM on HEK293T cells expressing TAS2R14 using the IP-One HTRF® assay.

| Catalog number | Item name | CAS number | Activation (%)[a] (rel. to Flufenamic acid) Mean ± S.D. |
|---|---|---|---|
| B2145 | Fosaprepitant dimeglumine salt | 265121-04-8 | 159 ± 4 |
| B4976 | Desogestrel | 54024-22-5 | 157 ± 7 |
| B5965 | Tamoxifen | 10540-29-1 | 154 ± 3 |
| C4087 | Mebendazole | 31431-39-7 | 138 ± 29 |
| B6140 | Hexachlorophene | 70-30-4 | 136 ± 23 |
| B1506 | Estradiol valerate | 979-32-8 | 123 ± 46 |
| N2558 | Sclareol | 515-03-7 | 122 ± 11 |
| B1401 | Cabozantinib malate (XL184) | 1140909-48-3 | 122 ± 11 |
| B1530 | Niflumic acid | 4394-00-7 | 121 ± 45 |

| | | | |
|---|---|---|---|
| B1674 | Benzethonium chloride | 121-54-0 | 120 ± 29 |
| B1517 | Gestodene | 60282-87-3 | 119 ± 23 |
| A3787 | Salirasib | 162520-00-5 | 118 ± 5 |
| B5846 | Radotinib(IY-5511) | 926037-48-1 | 115 ± 29 |
| B1905 | Carbimazole | 22232-54-8 | 113 ± 8 |
| N2032 | Aristolochic acid A | 313-67-7 | 111 ± 28 |
| A2977 | Cabozantinib (XL184, BMS-907351) | 849217-68-1 | 110 ± 7 |
| N1742 | Artemether | 71963-77-4 | 110 ± 33 |
| B1906 | Cefoselis Sulfate | 122841-12-7 | 109 ± 18 |
| B3680 | Griseofulvin | 126-07-8 | 107 ± 1 |
| B1904 | Carbidopa | 28860-95-9 | 107 ± 11 |
| B3470 | Etonogestrel | 54048-10-1 | 103 ± 13 |
| B1899 | Bromhexine HCl | 611-75-6 | 103 ± 7 |
| A5710 | Tazarotene | 118292-40-3 | 103 ± 28 |
| B2300 | Tolvaptan | 150683-30-0 | 103 ± 1 |
| B1991 | Norethindrone | 68-22-4 | 101 ± 25 |
| A5343 | Ispinesib (SB-715992) | 336113-53-2 | 101 ± 21 |
| B1505 | Epiandrosterone | 481-29-8 | 100 ± 28 |
| A5331 | TSU-68 (SU6668,Orantinib) | 252916-29-3 | 100 ± 36 |
| B1634 | Fosbretabulin (Combretastatin A4 Phosphate (CA4P)) Disodium | 168555-66-6 | 99 ± 10 |
| B1941 | Estradiol Benzoate | 50-50-0 | 98 ± 8 |
| B5859 | Entrectinib | 1108743-60-7 | 98 ± 26 |
| B6298 | Bromocriptine mesylate | 22260-51-1 | 97 ± 14 |
| B4962 | Thonzonium bromide | 553-08-2 | 97 ± 16 |
| N1790 | Podophyllotoxin | 518-28-5 | 97 ± 23 |
| N2250 | Vinorelbine | 71486-22-1 | 97 ± 1 |
| A8809 | Vorapaxar | 618385-01-6 | 95 ± 9 |
| B1831 | Sertaconazole nitrate | 99592-39-9 | 94 ± 17 |
| B1459 | Valdecoxib | 181695-72-7 | 93 ± 11 |
| B7201 | α-Estradiol | 57-91-0 | 93 ± 28 |
| B1960 | Levonorgestrel | 797-63-7 | 92 ± 25 |
| B1393 | Nitrendipine | 39562-70-4 | 92 ± 2 |
| B1760 | Flufenamic acid | 530-78-9 | 91 ± 8 |
| B1444 | Etodolac | 41340-25-4 | 90 ± 16 |
| N1318 | Vitexicarpin | 479-91-4 | 90 ± 71 |
| B1458 | Lumiracoxib | 220991-20-8 | 90 ± 4 |
| B1198 | Cisapride | 81098-60-4 | 89 ± 4 |
| A1659 | Dutasteride | 164656-23-9 | 89 ± 54 |
| B4796 | Meclofenamate sodium | 6385-02-0 | 89 ± 7 |
| B1457 | Diclofenac sodium | 15307-79-6 | 88 ± 35 |
| B1415 | Clevidipine butyrate | 167221-71-8 | 87 ± 16 |
| B1897 | Bifonazole | 60628-96-8 | 87 ± 10 |
| B1460 | Zaltoprofen | 74711-43-6 | 87 ± 17 |
| C3558 | 4-Methylbenzylidene camphor | 36861-47-9 | 85 ± 52 |
| C4201 | Androsterone | 53-41-8 | 85 ± 22 |
| B2266 | LDE225 (NVP-LDE225,Erismodegib) | 956697-53-3 | 84 ± 18 |
| N1949 | Panaxtriol | 32791-84-7 | 84 ± 30 |
| B1445 | Flunixin meglumin | 42461-84-7 | 83 ± 42 |
| A4188 | 2-Methoxyestradiol (2-MeOE2) | 362-07-2 | 83 ± 0 |

| | | | |
|---|---|---|---|
| B1510 | Medroxyprogesterone acetate | 71-58-9 | 82 ± 23 |
| N2256 | Vinblastine | 865-21-4 | 81 ± 5 |
| B1455 | Tolfenamic acid | 13710-19-5 | 80 ± 14 |
| B1512 | Pregnenolone | 145-13-1 | 80 ± 37 |
| A5143 | Finasteride | 98319-26-7 | 80 ± 22 |
| B2224 | Etravirine (TMC125) | 269055-15-4 | 79 ± 23 |
| B1454 | Rofecoxib | 162011-90-7 | 79 ± 20 |
| C6108 | L-Alanine | 56-41-7 | 78 ± 37 |
| C3108 | Linoleic Acid | 60-33-3 | 78 ± 19 |
| B1389 | Amiodarone HCl | 19774-82-4 | 76 ± 1 |
| B1378 | Spironolactone | 52-01-7 | 75 ± 15 |
| A8379 | Betamethasone Valerate | 2152-44-5 | 74 ± 21 |
| B4826 | Isavuconazole | 241479-67-4 | 74 ± 7 |
| N1486 | Lathyrol | 34420-19-4 | 74 ± 5 |
| B1903 | Carbazochrome sodium sulfonate (AC-17) | 51460-26-5 | 74 ± 11 |
| B1449 | Mefenamic acid | 61-68-7 | 73 ± 15 |
| C6358 | Fluorescein | 2321-07-5 | 72 ± 2 |
| B1937 | Econazole nitrate | 24169-02-6 | 71 ± 17 |
| B1901 | Butenafine HCl | 101827-46-7 | 71 ± 17 |
| B1900 | Budesonide | 51333-22-3 | 71 ± 1 |
| A8202 | Abiraterone acetate | 154229-18-2 | 70 ± 4 |
| B8214 | Dihydrotestosterone(DHT) | 521-18-6 | 70 ± 32 |
| N1901 | Z-Ligustilide | 4431-01-0 | 69 ± 28 |
| A1296 | Exemestane | 107868-30-4 | 67 ± 8 |
| B1375 | Dehydroepiandrosterone (DHEA) | 53-43-0 | 67 ± 1 |
| B3668 | Sodium monensin | 22373-78-0 | 67 ± 6 |
| B2104 | Itraconazole | 84625-61-6 | 67 ± 4 |
| B1798 | Nicardipine HCl | 54527-84-3 | 66 ± 14 |
| B1977 | Miconazole | 22916-47-8 | 66 ± 16 |
| A4240 | Abiraterone | 154229-19-3 | 66 ± 4 |
| A3543 | LDE225 diphosphate | 1218778-77-8 | 66 ± 19 |
| B1767 | Halobetasol propionate | 66852-54-8 | 66 ± 3 |
| B2134 | Ulipristal | 159811-51-5 | 66 ± 8 |
| B3505 | Diclofenac | 15307-86-5 | 65 ± 19 |
| A8553 | PTC124 (Ataluren) | 775304-57-9 | 64 ± 4 |
| B2156 | Bindarit | 130641-38-2 | 64 ± 7 |
| B1902 | Butoconazole nitrate | 64872-77-1 | 64 ± 33 |
| B2203 | Candesartan cilexetil | 145040-37-5 | 63 ± 14 |
| C4274 | Quinestrol | 152-43-2 | 62 ± 14 |
| A4014 | Artemether (SM-224) | 71963-77-4 | 62 ± 16 |
| B3542 | Tetrabenazine | 58-46-8 | 61 ± 20 |
| B1999 | Oxibendazole | 20559-55-1 | 61 ± 6 |
| B1788 | Mevastatin | 73573-88-3 | 61 ± 8 |
| B1989 | Nisoldipine | 63675-72-9 | 61 ± 16 |
| B1496 | Icotinib | 610798-31-7 | 61 ± 18 |
| B1409 | Benidipine HCl | 91599-74-5 | 60 ± 6 |
| C5167 | Fusidic acid (sodium salt) | 751-94-0 | 60 ± 3 |
| B4885 | Lomitapide | 182431-12-5 | 60 ± 29 |
| A1435 | CP-945598 HCl | 686347-12-6 | 58 ± 18 |

| | | | |
|---|---|---|---|
| A3549 | Lesinurad | 878672-00-5 | 58 ± 4 |
| A3403 | Etofenamate | 30544-47-9 | 58 ± 13 |
| A8455 | Nilvadipine | 75530-68-6 | 58 ± 4 |
| A8532 | Teniposide | 29767-20-2 | 57 ± 10 |
| B2106 | Loteprednol etabonate | 82034-46-6 | 57 ± 8 |
| A3921 | Vinorelbine ditartrate | 125317-39-7 | 56 ± 11 |
| B1724 | Deoxycorticosterone acetate | 56-47-3 | 55 ± 17 |
| B1728 | Difluprednate | 23674-86-4 | 55 ± 6 |
| A2174 | Lenvatinib (E7080) | 417716-92-8 | 55 ± 2 |
| A2974 | Foretinib (GSK1363089) | 849217-64-7 | 55 ± 5 |
| C6100 | Glycine | 56-40-6 | 54 ± 10 |
| B2133 | Suprofen | 40828-46-4 | 54 ± 21 |
| B2036 | Sulconazole nitrate | 61318-91-0 | 54 ± 4 |
| B1915 | Clobetasol propionate | 25122-46-7 | 53 ± 13 |
| A3791 | Saquinavir mesylate | 149845-06-7 | 53 ± 16 |
| B4888 | Obeticholic acid | 459789-99-2 | 53 ± 1 |
| A8623 | TOK-001 | 851983-85-2 | 53 ± 1 |
| A3222 | Baricitinib phosphate | 1187595-84-1 | 53 ± 5 |
| B1943 | Fenofibrate | 49562-28-9 | 53 ± 8 |
| B1740 | Ebastine | 90729-43-4 | 53 ± 12 |
| A8651 | LY2940680 | 1258861-20-9 | 52 ± 6 |
| C6107 | p-Anisaldehyde | 123-11-5 | 51 ± 14 |
| B1649 | Racecadotril | 81110-73-8 | 51 ± 9 |
| A8426 | Estrone | 53-16-7 | 51 ± 2 |
| B6139 | (+/-)-Sulfinpyrazone | 57-96-5 | 50 ± 7 |
| A8358 | Albendazole | 54965-21-8 | 50 ± 3 |
| B7456 | Allopregnanolone | 516-54-1 | 50 ± 6 |
| A1684 | Aprepitant | 170729-80-3 | 50 ± 25 |
| B1928 | Diclofenac diethylamine | 78213-16-8 | 49 ± 14 |
| A8453 | Isradipine (Dynacirc) | 75695-93-1 | 49 ± 2 |
| C4195 | Menaquinone 4 | 863-61-6 | 49 ± 6 |
| B1718 | Cyclandelate | 456-59-7 | 47 ± 7 |
| A3631 | Monomethyl auristatin E | 474645-27-7 | 47 ± 4 |
| B2108 | Lubiprostone | 136790-76-6 | 46 ± 4 |
| B8702 | Flunisolide | 3385-03-3 | 46 ± 6 |
| A8534 | Terfenadine | 50679-08-8 | 46 ± 6 |
| A5618 | Daclatasvir (BMS-790052) | 1214735-16-6 | 45 ± 11 |
| A4329 | Cilomilast | 153259-65-5 | 44 ± 10 |
| A1447 | Cilnidipine | 132203-70-4 | 44 ± 13 |
| B1938 | Erdosteine | 84611-23-4 | 43 ± 29 |
| B2296 | Topotecan HCl | 119413-54-6 | 42 ± 6 |
| A8509 | Progesterone | 57-83-0 | 42 ± 4 |
| B1248 | Beclomethasone dipropionate | 5534-09-8 | 42 ± 8 |
| A8454 | Istradefylline (KW-6002) | 155270-99-8 | 41 ± 14 |
| A8484 | Nimodipine | 66085-59-4 | 39 ± 7 |
| A8431 | Famprofazone | 22881-35-2 | 40 ± 4 |
| B1935 | Dronedarone HCl | 141625-93-6 | 39 ± 16 |
| B1896 | Betamethasone | 378-44-9 | 38 ± 13 |

| | | | |
|---|---|---|---|
| B1929 | Diclofenac potassium | 15307-81-0 | 38 ± 22 |
| N1500 | Germacrone | 6902-91-6 | 37 ± 6 |
| B1995 | (+,-)-Octopamine HCl | 770-05-8 | 36 ± 19 |
| B1240 | (+)-Ketoconazole | 142128-59-4 | 35 ± 2 |
| A3496 | Indacaterol | 312753-06-3 | 32 ± 11 |
| A8479 | Noscapine HCl | 912-60-7 | 32 ± 0 |

*[a]Activation (%): relative to 30 µM Flufenamic acid (100 % activation). Activation threshold (%) for the identification of an active compound, or "hit," was fixed as the mean of the unstimulated control plus three times the standard deviation (S.D.). Data represent mean ± S.D. of duplicate assay plates.*

**Supplementary File 3 Table S2.** Results confirmation screens - list of active library compounds screened at a concentration of 3 µM, 1 µM and 0.3 µM on HEK293T cells expressing TAS2R14 using the IP-One HTRF® assay.

| **Item name** | **CAS number** | **Screening Concentration** | | |
|---|---|---|---|---|
| | | ***3 µM[a]*** | ***1 µM[a]*** | ***0.3 µM [a]*** |
| | | **Activation (%) (rel. to Flufenamic acid)** *Mean ± S.D.* | | |
| Cabozantinib (XL184, BMS-907351) | 849217-68-1 | 133 ± 29 | 55 ± 2 | - |
| Cabozantinib malate (XL184) | 1140909-48-3 | 132 ± 25 | 77 ± 1 | - |
| Aristolochic acid A | 313-67-7 | 120 ± 1 | 89 ± 4 | 49 ± 13 |
| Bromocriptine mesylate | 22260-51-1 | 113 ± 35 | 109 ± 4 | 83 ± 12 |
| Etonogestrel | 54048-10-1 | 110 ± 3 | 81 ± 2 | 58 ± 11 |
| Vinorelbine | 71486-22-1 | 107 ± 4 | 74 ± 21[c] | 61 ± 15 |
| Podophyllotoxin | 518-28-5 | 105 ± 24 | 82 ± 31 [c] | 76 ± 11 |
| Levonorgestrel | 797-63-7 | 104 ± 40 | 79 ± 3 | 51 ± 7 |
| Vinblastine | 865-21-4 | 101 ± 16 | 81 ± 36 [c] | 74 ± 14 |
| Fosbretabulin (Combretastatin A4 Phosphate (CA4P)) Disodium | 168555-66-6 | 98 ± 16 | 68 ± 13 [c] | 80 ± 6 |
| Desogestrel | 54024-22-5 | 96 ± 21 | 54 ± 8 | 40 ± 11 |
| Flufenamic acid | 530-78-9 | 95 ± 23 | 78 ± 12 | 45 ± 10 |
| Mebendazole | 31431-39-7 | 95 ± 27 | - | - |
| Butoconazole nitrate | 64872-77-1 | 95 ± 24 | 71 ± 5 | 44 ± 20 |
| Niflumic acid | 4394-00-7 | 94 ± 9 | 61 ± 13 | - |
| TSU-68 (SU6668,Orantinib) | 252916-29-3 | 93 ± 14 | 47 ± 6 | - |
| Tolfenamic Acid | 13710-19-5 | 91 ± 18 | - | - |
| Meclofenamate Sodium | 6385-02-0 | 86 ± 12 | | - |
| Isavuconazole | 241479-67-4 | 84 ± 39 | 54 ± 5 | |
| Gestodene | 60282-87-3 | 82 ± 6 | 58 ± 13 | |
| Vorapaxar | 618385-01-6 | 80 ± 40 | 68 ± 8 | 48 ± 11 |
| Mefenamic acid | 61-68-7 | 79 ± 18 | | |
| Vinorelbine ditartrate | 125317-39-7 | 78 ± 8 | 55 ± 8 [c] | 61 ± 15 |
| Monomethyl auristatin E | 474645-27-7 | 78 ± 5 | - | - |
| TOK-001 | 851983-85-2 | 76 ± 11 | - | - |
| Artemether | 71963-77-4 | 74 ± 15 | - | - |
| Griseofulvin | 126-07-8 | 74 ± 5 | - | - |
| Sulconazole nitrate | 61318-91-0 | 69 ± 11 | 44 ± 7 | 34 ± 18 |
| Albendazole | 54965-21-8 | 68 ± 6 | - | - |
| Irisflorentin | 41743-73-1 | 66 ± 47 | - | - |

| | | | | |
|---|---|---|---|---|
| Halobetasol propionate | 66852-54-8 | 64 ± 6 | - | - |
| Oxibendazole | 20559-55-1 | 62 ± 32 | 68 ± 2 | - |
| Foretinib (GSK1363089) | 849217-64-7 | 56 ± 6 | - | - |
| Medroxyprogesterone acetate | 71-58-9 | 53 ± 6 | - | - |
| Exemestane | 107868-30-4 | 52 ± 5 | - | - |
| Flunixin meglumin | 42461-84-7 | 51 ± 1 | - | - |
| Dutasteride | 164656-23-9 | 50 ± 5 | - | - |
| Deoxycorticosterone acetate | 56-47-3 | 49 ± 6 | - | - |
| LDE225 Diphosphate | 1218778-77-8 | 49 ± 4 | - | - |
| Loteprednol etabonate | 82034-46-6 | 46 ± 1 | - | - |
| Norethindrone | 68-22-4 | 44 ± 1 | 32 ± 7 | - |
| Itraconazole | 84625-61-6 | 43 ± 16 | 51 ± 2 | - |

[a]*Activation (%): relative to 30 µM Flufenamic acid (100 % activation). Activation threshold (%) for the identification of an active compound, or "hit," was fixed as the mean of the unstimulated control plus three times the standard deviation (S.D.). Data represent mean ± (S.D.) of duplicate assay plates* [a] *and of two independent experiments.* [c] *Unspecific activation (see Supplementary Figure S2).*

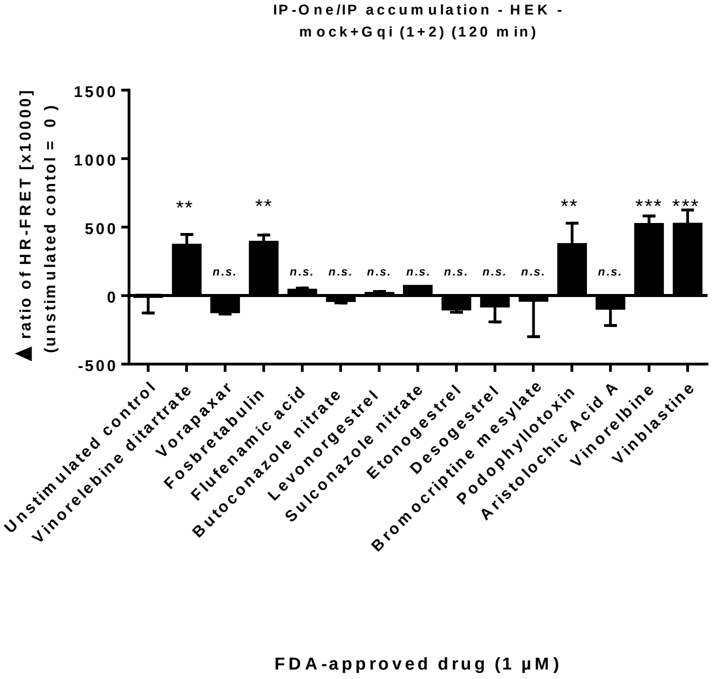


**Supplementary File 3 Fig. S2: Specificity of initial active library compounds selected from the screen at 0.3 µM.** Fourteen compounds identified as active compounds of the FDA-approved drug library on HEK293T cells expressing TAS2R14 were tested again (1 µM) using mock-transfected HEK293T cells to determine whether the compounds specifically activate TAS2R14. Data represent mean ± (S.D.) of one representative experiment ($n$=2) performed in duplicate. Statistical analysis was performed by an unpaired $t$-test (n.s. not significant, **p-value < 0.01 and ***p-value < 0.001 versus unstimulated control (0.3 % (v/v) DMSO), respectively)
